# Direction, Not Distance: A Mortality-Associated Physiological Axis Maps the Sex–Mortality Gap and Informs Sex-Balanced Trial Recruitment

**DOI:** 10.64898/2026.03.12.711462

**Authors:** Angus Silas Harding, Jim Coward, Jinping Feng, Tianhai Tian

## Abstract

Geroscience has more candidate interventions than it can afford to test. Trials powered for mortality or functional decline require large cohorts and long follow-up, and the persistent female survival advantage could further increase sample-size requirements if women and men age through different biology. Existing ageing biomarkers — including frailty indices, biological-age clocks, allostatic load and homeostatic dysregulation — compress multidimensional physiology to a scalar, discarding which combination of biomarkers changed. In 29,053 US adults measured on 38 routine laboratory and examination markers, we decomposed displacement from sex-specific young-healthy physiology into total magnitude and alignment with a cross-fitted mortality-associated direction. Women were no closer to young-health than men but had substantially lower disease mortality. Adjustment for magnitude widened the female survival advantage; adjustment for mortality-directed projection attenuated 87% of the sex difference on the log-hazard scale. Sex-specific directions transferred almost without loss, indicating a substantially shared predictive geometry with different positions along it. Across the population, a median 68% of the positive mortality-associated contribution came from measurements lying within conventional clinical limits and 57% from within the young-healthy range; these shares reflect how numerous such measurements are, not greater information per measurement. Ten routine measurements reproduced the 38-marker direction, and seven of eight predictions registered before analysis were confirmed in NHANES III. High-sensitivity C-reactive protein was strongly prognostic alone but showed no incremental information beyond the routine panel, whereas measuring lung-function added more held-out prognostic information than 33 of 34 individual blood and urine markers in the replication cohort. Retrospective sorting on this geometry produced mortality-enriched and physiologically displaced candidate trial groups; pooled thresholds were strongly sex-imbalanced, whereas within-sex thresholds restored balance with only a 1.5–3.8% reduction in expected five-year events. These findings suggest that inexpensive routine biomarkers plus simple functional testing could support smaller, more informative and sex-balanced geroscience trials, but prospective validation of treatment enrichment is required.

## Introduction

Geroscience aims to delay age-related disease and functional decline by targeting shared biological processes of ageing, thereby extending healthspan and potentially lifespan ^1,2^. Clinical testing is difficult because mortality, multimorbidity, disability and loss of physical or cognitive function accumulate slowly and heterogeneously. Trials powered for these outcomes therefore require large cohorts, long follow-up and substantial investment ^3,4^. Biomarkers could reduce this burden by identifying participants with relevant physiology, enriching event rates where appropriate, supporting mechanistic stratification and providing earlier evidence that an intervention changed biological state ^5–7^.

A second constraint is the persistent sex difference in mortality ^8^. Women outlive men across diverse populations while often reporting more diagnosed chronic disease — the morbidity–mortality paradox — and the gap is not fully explained by health behaviour, socioeconomic position, reported morbidity, cause-of-death distribution or the menopausal transition ^9–11^. For geroscience, the distinction matters. If women and men reach mortality risk through fundamentally different physiology, trials may require substantially larger sex-stratified samples and, potentially, sex-specific intervention development. If they occupy different positions within a largely shared physiological system, one instrument and one intervention pipeline may serve both, provided recruitment accounts for the offset.

Ageing instruments such as frailty indices ^12–14^, biological-age clocks ^15,16^, allostatic load ^17^ and homeostatic dysregulation ^18^ use different inputs and algorithms, but each compresses multidimensional physiology to a scalar score. That compression is useful for predicting age-related mortality, disability and disease, as well as informing clinical decisions, but it discards configuration. Two people can be equally far from young-healthy physiology — or have the same aggregate score — while differing in which biomarkers changed and whether that combination resembles physiology associated with later death. Magnitude answers how much physiology has changed it does not specify the pattern of change. Whether this missing directional information carries prognostic signal beyond magnitude has not, to our knowledge, been tested directly in a large human cohort.

There is reason to expect that routine measurements can recover such structure. Ageing reflects interactions among coupled physiological systems ^19,20^; organs age asynchronously, dysfunction can propagate across systems, and organ-specific biological age predicts mortality beyond chronological age ^21,22^. Mathematical models based on cascading subsystem failure ^23^, network damage ^24^ and senescent-cell feedback ^25^ likewise show that sparse observations of interconnected systems can recover organism-level ageing dynamics, consistent with empirical work on sparsely sampled physiological networks ^26^. Coupling therefore supports sparse measurement, but also makes configuration important: which systems move, and in what combination.

The female survival advantage provides a strong test of this idea. Women are not consistently less dysregulated than men and, on some instruments, appear more dysregulated despite lower mortality ^9,10^. If magnitude alone were sufficient, this pattern would be difficult to explain. Sex therefore provides an external grouping variable with which to ask whether physiological direction contains information that total dysregulation misses.

We tested two linked hypotheses. The primary hypothesis was that the female–male mortality difference is better captured by the configuration of physiological displacement than by its magnitude. The secondary hypothesis was that, if the mortality-associated configuration is substantially shared between sexes, it should compress to a small routine panel, replicate across survey eras, recover established adverse exposures not used to fit it, and support trial enrichment without forcing sex imbalance. We tested these hypotheses in 29,053 US adults using 38 routine laboratory and examination measurements referenced to sex-specific young-healthy physiology, with mortality supervision introduced only through cross-fitted Cox models.

We then asked where the resulting mortality-associated signal lies relative to clinical reference limits, how far it can be reduced without losing information, whether it replicates in NHANES III (1988– 1994), what additional information is contributed by inflammatory, metabolic and functional measurements, and how pooled versus within-sex thresholds alter the composition of candidate lifespan- and healthspan-oriented trial groups.

## Results

### 1. Women Had Lower Disease Mortality Despite Greater Reported Morbidity, and Dysregulation Magnitude Did Not Account for It

We first established the contrast motivating the study. Among 29,250 NHANES 1999–2018 participants aged 18–79 years (14,190 women, 15,060 men), women and men were closely matched on age, race and ethnicity, and income. Women nevertheless reported more diagnosed disease: a mean of 0.87 versus 0.73 of thirteen doctor-diagnosed conditions, and 48.2% versus 39.2% reported at least one among the 28,232 participants answering all thirteen items. Disease death occurred in 9.5% of women and 13.4% of men (male-to-female risk ratio 1.40, 95% confidence interval 1.32 to 1.50). In age-adjusted Cox regression among 29,248 participants with 3,369 disease deaths, the female hazard ratio was 0.644 (0.601 to 0.690) and changed little after adjustment for sociodemographic factors, health behaviours or morbidity count (Fig. 1a–c, Table 1).

**Figure 1.**
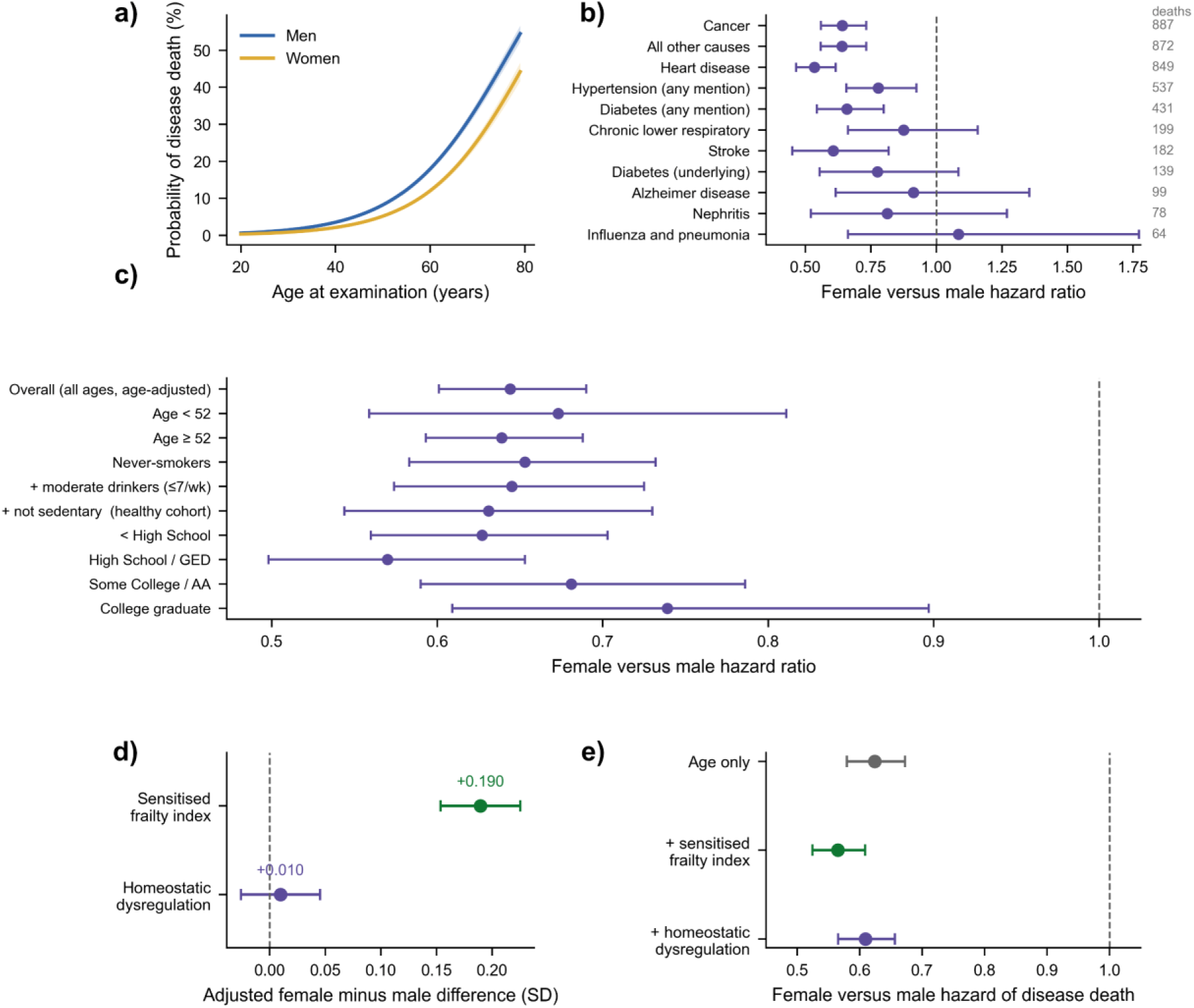
Women died of disease less often than men at every adult age, and neither measure of dysregulation magnitude accounted for it. **a**, Probability of disease death by age at examination, estimated separately in men (blue) and women (yellow); shaded bands are 95% confidence intervals. **b**, Female versus male hazard ratio for disease death by underlying cause, from Cox models adjusted for age, with causes ordered by the number of deaths contributing (given in grey at the right of the panel). Two causes are reported on any-mention rather than underlying-cause certification, as labelled. **c**, Female versus male hazard ratio for all disease death within pre-specified strata: overall, either side of the age-52 boundary, within a progressively restricted behaviourally healthy cohort, and within each educational stratum. Each estimate comes from a separate age-adjusted model fitted within that stratum. **d**, Adjusted female-minus-male difference in the sensitised frailty index (green) and in homeostatic dysregulation (purple) among participants aged 52 years and above (11,509 participants; 5,580 women, 5,929 men, twelve of whom lack linked mortality follow-up and so do not contribute to panel e), expressed in standard deviations of the analysis population and adjusted for age using a cubic B-spline with four degrees of freedom and for survey cycle as a fixed effect. Intervals are heteroskedasticity-consistent (HC3). **e**, Female versus male hazard of disease death adjusted for age alone (grey), and after further adjustment for the sensitised frailty index (green) or for homeostatic dysregulation (purple). Points are estimates and horizontal bars are 95% confidence intervals throughout. The dashed vertical line marks a hazard ratio of 1 in panels **b**, **c** and **e**, and a difference of zero in panel **d**. Cause-specific and stratified models (panels **a** to **c**) are estimated in 29,248 participants with 3,369 disease deaths; the survival models in panel e in 29,023 participants with 2,922 disease deaths, the smaller cohort reflecting both complete biomarker measurement and available mortality linkage. Accidental death is excluded from the disease-death endpoint and treated as a competing event.

**Table 1.** Characteristics of the corrected NHANES 1999–2018 analysis cohort by sex. Participants were non-pregnant adults aged 18–79 years without acute illness at examination and with complete data for the tabulated demographic, mortality, smoking, alcohol, and physical-activity fields (N=29,250; 14,190 women and 15,060 men). Continuous variables are mean (SD), except alcohol, which is median [Q1, Q3]; categorical variables are n (%). The historical morbidity count is the sum of 13 doctor-diagnosed conditions and is shown among participants answering all 13 items. Covariate differences are described using standardised mean differences and, for skewed or ordinal variables, Cliff’s delta. Mortality endpoints report men-to-women risk ratios with 95% confidence intervals. Disease death excludes accidental death; follow-up is measured from examination. Estimates are unweighted descriptions of the analytic cohort.

| Characteristic | Women (n = 14,190) | Men (n = 15,060) | Overall (n = 29,250) | Standardised difference |
| --- | --- | --- | --- | --- |
| Age, years - mean (SD) | 48.5 (16.4) | 48.1 (16.7) | 48.3 (16.6) | SMD 0.021 |
| Diagnosed conditions of 13 - mean (SD) | 0.9 (1.2) | 0.7 (1.2) | 0.8 (1.2) | SMD 0.112 |
| Disease death during follow-up | 1,354 (9.5) | 2,017 (13.4) | 3,371 (11.5) | RR 1.40 (1.32-1.50) |
| Accidental death during follow-up | 45 (0.3) | 71 (0.5) | 116 (0.4) | RR 1.49 (1.02-2.16) |
| Alcohol, drinks per week - median [Q1, Q3] | 0.1 [0.0, 1.0] | 0.9 [0.0, 6.0] | 0.2 [0.0, 3.0] | SMD 0.413 |
| *Smoking status* |  |  |  | SMD 0.360 |
| Never | 8,975 (63.2) | 6,835 (45.4) | 15,810 (54.1) |  |
| Former | 2,712 (19.1) | 4,466 (29.7) | 7,178 (24.5) |  |
| Current | 2,503 (17.6) | 3,759 (25.0) | 6,262 (21.4) |  |
| *Physical activity* |  |  |  | SMD 0.180 |
| Active | 10,222 (72.0) | 12,008 (79.7) | 22,230 (76.0) |  |
| Inactive (sedentary) | 3,968 (28.0) | 3,052 (20.3) | 7,020 (24.0) |  |
| *Education* |  |  |  | SMD 0.114 |
| <High School | 3,151 (22.2) | 3,837 (25.5) | 6,988 (23.9) |  |
| High School/GED | 3,200 (22.6) | 3,648 (24.2) | 6,848 (23.4) |  |
| Some College/AA | 4,476 (31.5) | 4,058 (26.9) | 8,534 (29.2) |  |
| College Graduate | 3,357 (23.7) | 3,510 (23.3) | 6,867 (23.5) |  |
| Non-responder | 6 (0.0) | 7 (0.0) | 13 (0.0) |  |
| *Poverty-to-income ratio* |  |  |  | SMD 0.070 |
| ≤100% FPL | 2,863 (20.2) | 2,744 (18.2) | 5,607 (19.2) |  |
| ≤185% FPL | 3,249 (22.9) | 3,355 (22.3) | 6,604 (22.6) |  |
| ≤300% FPL | 2,592 (18.3) | 2,781 (18.5) | 5,373 (18.4) |  |
| ≤500% FPL | 2,912 (20.5) | 3,105 (20.6) | 6,017 (20.6) |  |
| >500% FPL | 2,574 (18.1) | 3,075 (20.4) | 5,649 (19.3) |  |
| *Race/ethnicity* |  |  |  | SMD 0.019 |
| Hispanic | 3,467 (24.4) | 3,668 (24.4) | 7,135 (24.4) |  |
| Non-Hispanic Black | 3,081 (21.7) | 3,169 (21.0) | 6,250 (21.4) |  |
| Non-Hispanic White | 6,322 (44.6) | 6,830 (45.4) | 13,152 (45.0) |  |
| Other/Multiracial | 1,320 (9.3) | 1,393 (9.2) | 2,713 (9.3) |  |

We next asked whether women simply accumulated less physiological dysregulation. Two outcome-independent magnitude measures were used: covariance-adjusted Mahalanobis distance from a sex-specific young-healthy centroid (HD), and a sensitised linear frailty index (FI) equal to the mean adverse excursion across 38 routine markers.

Both measures strongly predicted mortality. Per cohort standard deviation, among 29,023 participants with 2,922 disease deaths, the hazard ratio was 1.71 (1.65 to 1.78) for FI and 1.67 (1.61 to 1.73) for HD, with no evidence of sex interaction (likelihood-ratio P = 0.273 and P = 0.746). Entered together, both retained associations with mortality, indicating overlapping but non-identical information.

Neither measure explained the female survival advantage. At age 52 years and above, the adjusted female-minus-male difference was +0.190 standard deviations (0.154 to 0.225) for FI but +0.010 (−0.026 to 0.045) for HD, compatible with no difference. Adjustment moved the female hazard ratio farther from unity: from 0.624 to 0.565 (0.524 to 0.609) for FI and to 0.609 (0.566 to 0.656) for HD (Fig. 1d–e). In these data, total dysregulation therefore widened rather than resolved the mortality paradox.

### 2. Women Were Markedly Less Aligned With a Shared Mortality-Associated Direction

Distance alone cannot distinguish different patterns of biomarker change. Two people can be equally far from young-health yet differ in which markers moved and whether those changes align with a pattern associated with later death. We therefore decomposed each participant’s displacement into magnitude and orientation.

Among 11,497 participants aged 52–79 years with 2,494 disease deaths, we estimated a mortality-associated direction by age- and sex-adjusted Cox regression on the 38-marker coordinate vector. Models were fitted leave-one-survey-cycle-out, so no participant’s score used outcomes from their own cycle. For each participant we then calculated magnitude m, alignment a — the cosine of the angle between their displacement and the fitted direction — and projection q = m × a, the amount of displacement lying along that direction.

Women were at least as far from young-health as men but were oriented differently. In Mahalanobis coordinates, adjusted female-minus-male differences were +0.232 standard deviations (95% bootstrap interval 0.199 to 0.268) for magnitude, −0.735 (−0.836 to −0.569) for alignment and −0.643 (−0.752 to −0.488) for projection; the frailty encoding was similar (+0.140, −0.743, −0.578). This ordering was preserved in all 252 directed survey-cycle partitions in both encodings (Fig. 2a–c).

**Figure 2.**
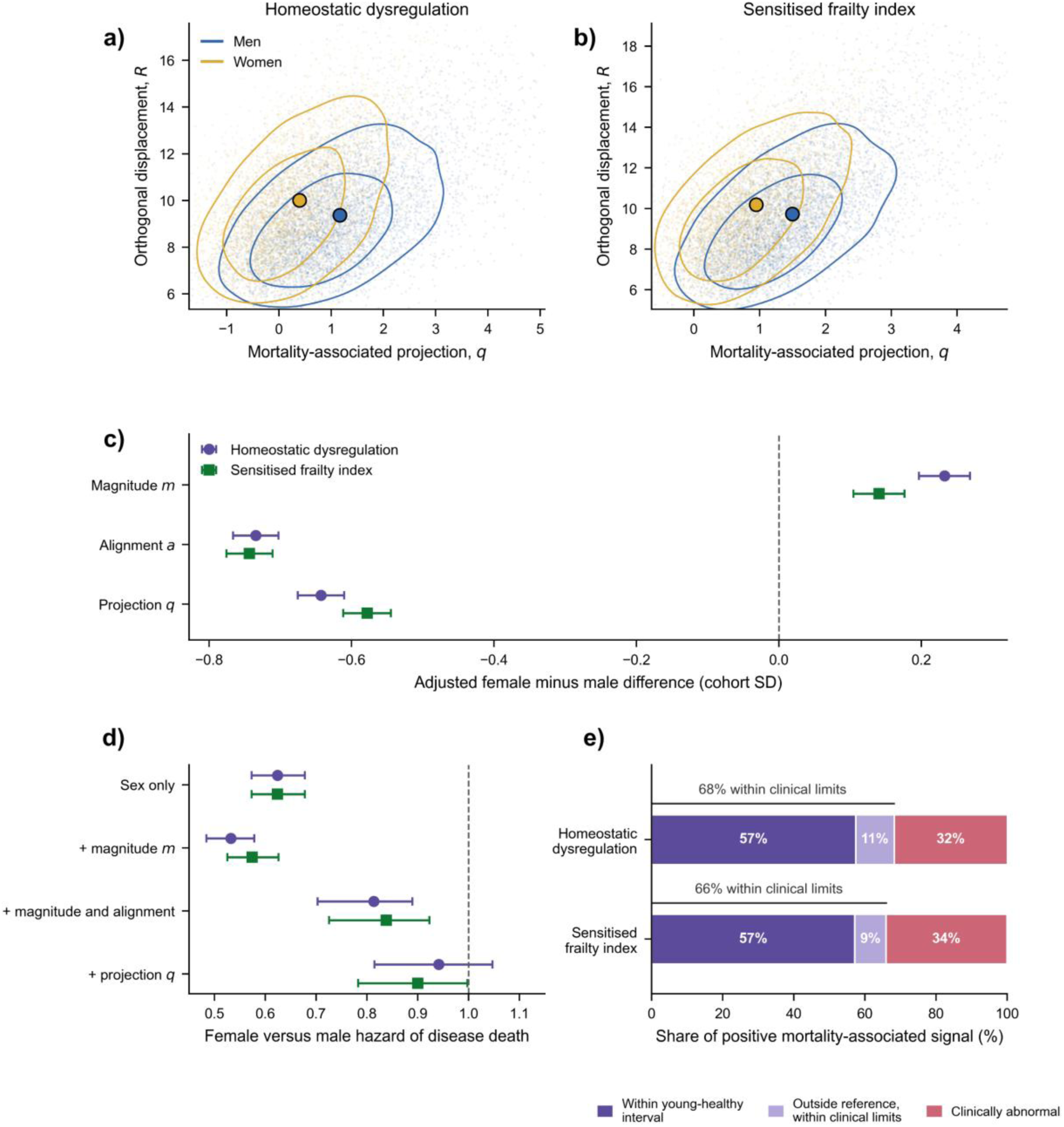
Women were displaced at least as far from young-health as men but in a direction less aligned with mortality, and projection along that direction attenuated most of the survival difference. **a**, **b**, Participant geometry among 11,497 participants aged 52–79 years with 2,494 disease deaths, in homeostatic-dysregulation coordinates (**a**) and sensitised frailty-index coordinates (**b**). The horizontal axis is the mortality-associated projection, the component of a participant’s displacement lying along the fitted mortality direction; the vertical axis is orthogonal displacement, the component lying at right angles to it. Each point is one participant, coloured blue for men and yellow for women. Contours enclose the densest 50% and 80% of each sex, estimated by kernel density. Large black-outlined circles mark sex centroids in these two coordinates; they are not mean 38-dimensional physiological vectors. **c**, Adjusted female-minus-male differences in displacement magnitude, alignment and projection, in standard deviations of the analysis population, for homeostatic dysregulation (purple circles) and the sensitised frailty index (green squares). Alignment is the cosine of the angle between a participant’s displacement and the fitted mortality direction, so it is scale-free; projection is magnitude multiplied by alignment. Bars in this panel are heteroskedasticity-consistent (HC3) intervals conditional on the fitted direction; the intervals quoted for these three quantities in the text come from the participant bootstrap, which refits the direction in every replicate and is correspondingly wider for alignment and projection. **d**, Female versus male hazard of disease death across a nested adjustment cascade, in both representations. Intervals are bootstrap percentile intervals. **e**, Where positive mortality-associated signal sits relative to conventional clinical thresholds, as a share of total positive contribution: within the same-sex young-healthy interval, outside that reference but still within conventional clinical limits, or clinically abnormal. Bars are medians across 20 survey-cycle partitions. The bracket above each bar gives the combined share falling within clinical limits. Points are estimates and horizontal bars are 95% intervals throughout. The mortality direction was fitted leave-one-survey-cycle-out, so no participant’s coordinates derive from any outcome observed in their own cycle. Where intervals overlapped, clinically abnormal classification took precedence, making the within-limits share in **e** conservative.

Projection attenuated the survival difference where magnitude did not. The female hazard ratio moved from 0.624 to 0.532 after adjustment for magnitude, but to 0.941 (95% bootstrap interval 0.815 to 1.047) after adjustment for projection, corresponding to an 87% attenuation of the sex difference on the log-hazard scale. In the frailty encoding, projection attenuated the difference by 78%, leaving a hazard ratio of 0.900 (0.783 to 0.997) (Fig. 2d).

The fitted direction was substantially shared between sexes. Sex-specific directions produced held-out scores correlated at 0.868 to 0.910, and applying the direction trained in the other sex retained 98.6% to 104.7% of the own-sex log-hazard association. Allowing separate sex-specific directions did not improve pooled held-out partial log-likelihood. The predictive axis was therefore largely shared, although women and men occupied different positions along it.

Two qualifications are important. Projection was more strongly associated with mortality in women than men (hazard ratio 2.068 versus 1.764 per standard deviation; multiplicative ratio 1.174 in both encodings), and its association declined with attained age, from 2.427 at ages 52–64 to 1.331 at age 85 and above. The reported per-standard-deviation hazard ratios are therefore cohort-averaged associations, not constant multipliers.

### 3. Most Mortality-Associated Information Arose Within Conventional Clinical Limits

We next located the signal relative to clinical thresholds. For each held-out participant and marker, we multiplied the coordinate by its training-derived mortality coefficient, retained positive contributions, and classified the raw value as within the same-sex young-healthy interval, outside that interval but within conventional clinical limits, or clinically abnormal. Coefficients were cross-fitted between halves of twenty survey-cycle partitions; where intervals overlapped, clinically abnormal classification took precedence, making the within-limits estimate conservative.

A median 68.1% of positive mortality-associated contribution arose while the raw value remained within conventional clinical limits, and 57.4% while it remained within the same-sex young-healthy interval. The alternative encoding gave closely similar values (66.0% and 57.1%) (Fig. 2e).

Thus, the positive contribution was not concentrated just beyond abnormality thresholds; most of it came from values within the clinically normal range, including its healthy centre. This is a statement about where contribution accumulates in a population rather than about individual risk: near-reference values contribute the majority of the mass because there are many of them, not because any individual near-reference value is as informative as an abnormal one. Because most measurements sit within limits, a threshold-based summary nevertheless discards a large share of the total continuous signal available in a population.

### 4. Ten Routine Measurements Reproduced the Mortality-Associated Direction

A 38-marker panel is feasible for analysis but cumbersome for deployment. We therefore tested how far the mortality direction could be reduced using nested cycle-grouped validation. Within each training half, elastic-net Cox models ranked all 38 markers; candidate panels of 3, 5, 7, 10, 12 and 15 markers were refitted and applied unchanged to the disjoint half. All 126 five-versus-five cycle divisions were evaluated in both directions (252 directed evaluations), against eight adequacy criteria fixed before results were generated.

Performance improved with panel size. Seven markers failed three of the eight criteria; ten satisfied all eight, reproducing the full 38-marker projection at a median held-out correlation of 0.902 and retaining 0.913 of its mortality information. The binding criterion passed narrowly (0.9018 versus the prespecified 0.90 threshold); panels of eight and nine markers were not among the sizes fixed in advance (Fig. 3a–b).

**Figure 3.**
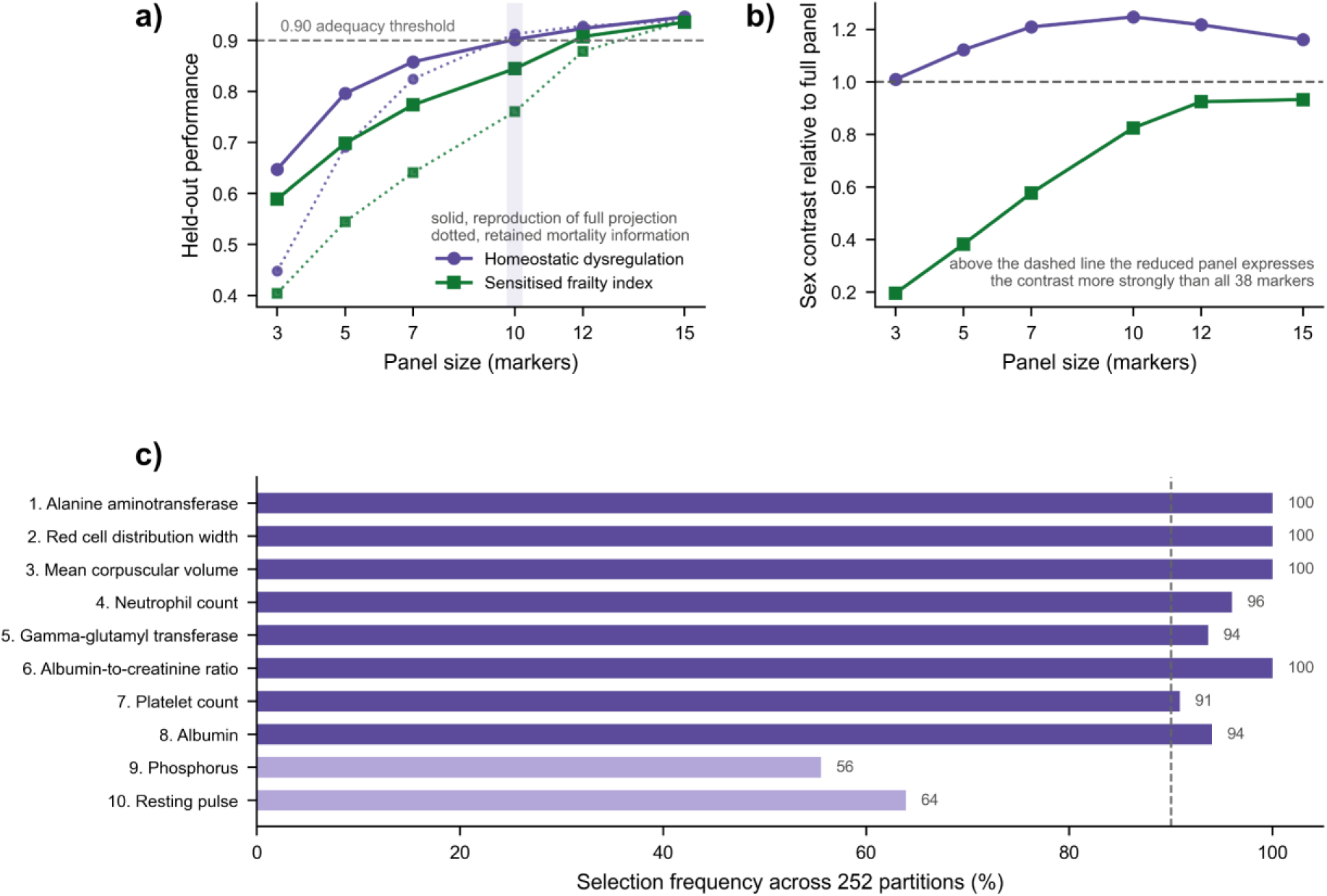
The mortality direction compressed to ten routine measurements, and reduction amplified rather than diluted the sex contrast. **a**, Held-out performance against panel size. Solid lines give the median correlation between the reduced-panel projection and the full 38-marker projection; dotted lines give the fraction of the full panel’s mortality information retained, both for homeostatic dysregulation (purple circles) and the sensitised frailty index (green squares). The dashed horizontal line marks the 0.90 adequacy threshold fixed before any result was generated; the shaded vertical band marks the selected panel size of ten. **b**, The adjusted sex contrast expressed by each reduced panel, relative to the contrast expressed by all 38 markers. Values above the dashed line at 1 indicate that the reduced panel expresses the contrast more strongly than the full panel. **c**, The ten selected markers, ranked by the magnitude of the penalised coefficient in the full development cohort, with the percentage of the 252 directed survey-cycle partitions in which each marker entered an independently fitted ten-marker panel. Bars are shaded darker where selection frequency reached 90% or above; the dashed line marks that level. Numerals at the end of each bar repeat the frequency. Panel sizes of 3, 5, 7, 10, 12 and 15 markers were fixed in advance; sizes of eight and nine were not evaluated. All 126 five-versus-five divisions of the ten survey cycles were evaluated in both directions, giving 252 directed evaluations, with markers ranked and panels refitted inside each training half and applied unchanged to the disjoint half. Selection frequency is an out-of-sample stability measure and is not a significance test. Marker identities, units, physiological systems and the direction on the raw measurement scale associated with higher mortality are given in Table 2.

**Table 2.** The ten markers selected by elastic-net penalised Cox regression in the full development cohort, ranked by the magnitude of the penalised coefficient. Direction is stated on the **raw measurement scale**: because three markers (alanine aminotransferase, gamma-glutamyl transferase, red cell distribution width) were analysed after a reciprocal transformation, which reverses the value axis, the raw-scale direction is the sign of the fitted coefficient multiplied by the recorded transformation orientation. Relative weight is the absolute penalised coefficient acting on the sex-standardised coordinate, so weights are comparable across markers and are expressed per standard deviation of the same-sex young-healthy reference; they are not clinical decision weights and are not intended for use as a bedside score without recalibration. Selected in (%) is the percentage of the 252 directed survey-cycle partitions in which the marker entered a ten-marker panel fitted independently within that partition’s training half — an out-of-sample stability measure, not a significance test. Lower alanine aminotransferase, rather than higher, marks elevated mortality in this age range, as observed previously ^27,28^. This is consistent with reduced hepatic mass and muscle mass in later life and runs opposite to the direction the same enzyme carries as a marker of acute hepatic injury. All ten markers are also contained in the fifteen selected by the same procedure run independently in the sensitised frailty-index encoding, whose adequate panel size was fifteen rather than ten.

| Rank | Measurement | NHANES variable | Units | System | Direction associated with higher mortality | Relative weight | Selected in (%) |
| --- | --- | --- | --- | --- | --- | --- | --- |
| 1 | Alanine aminotransferase | LBXSATSI | U/L | Hepatic | Lower | 0.23 | 100 |
| 2 | Red cell distribution width | LBXRDW | % | Red cell | Higher | 0.205 | 100 |
| 3 | Mean corpuscular volume | LBXMCVSI | fL | Red cell | Higher | 0.202 | 100 |
| 4 | Absolute neutrophil count | LBDNENO | 10<sup>9</sup>/L | Immune | Higher | 0.133 | 96 |
| 5 | Gamma-glutamyl transferase | LBXSGTSI | U/L | Hepatic | Higher | 0.127 | 94 |
| 6 | Urine albumin-to-creatinine ratio | ACR | mg/g | Renal | Higher | 0.122 | 100 |
| 7 | Platelet count | LBXPLTSI | 10<sup>9</sup>/L | Platelet | Lower | 0.119 | 91 |
| 8 | Serum albumin | LBXSAL | g/dL | Hepatic | Lower | 0.114 | 94 |
| 9 | Serum phosphorus | LBXSPH | mg/dL | Mineral | Higher | 0.089 | 56 |
| 10 | Resting pulse rate | BPXPLS | bpm | Cardiovascular | Higher | 0.087 | 64 |

Applied once to the complete development sample, the procedure selected alanine aminotransferase, red cell distribution width, mean corpuscular volume, neutrophil count, gamma-glutamyl transferase, urinary albumin-to-creatinine ratio, platelet count, albumin, phosphorus and resting pulse. The panel spans hepatic, red-cell, immune, renal, platelet, mineral and cardiovascular domains. The top eight markers were each selected in at least 90% of the 252 partitions; ranks 9 and 10 were less stable, identifying where the ten-marker panel begins to lose selection stability (Fig. 3c, Table 2).

Reduction amplified the sex contrast. At every candidate size, the adjusted female–male difference per standard deviation was larger than for all 38 markers, reaching 1.25 times the full-panel contrast at ten markers. Marker-wise attribution explained why: the selected ten contributed 108% of the full-panel contrast, whereas the remaining 28 contributed −8% overall. The largest opposing contributions came from uric acid, alkaline phosphatase, chloride, lactate dehydrogenase and systolic pressure, on which men deviated farther from their own young-healthy reference. Reduction therefore removed not only weakly informative markers but also markers that attenuated the sex contrast (Fig. 3b; Supplementary Information).

### 5. Established Adverse Exposures Mapped Differently Onto the Mortality Direction

The direction was fitted only to prospective disease mortality. To test whether it also tracked independently established adverse exposures, we examined current smoking, physical inactivity and poorer diet. None contributed to biomarker selection, direction estimation, panel size or panel identity, and predictions were recorded before these analyses were run.

Among 11,497 participants aged 52–79 years, each exposure was associated with magnitude, alignment and projection in both representations and both reduced panels. Coverage was 99.9% for smoking, 95.4% for physical activity and 65.8% for diet, because dietary recall was available only from 2005–2006 onward.

All three exposures were associated with higher projection: current smoking by 0.65 standard deviations (0.60 to 0.70), physical inactivity by 0.28 (0.24 to 0.32) and poorer diet by 0.17 (0.15 to 0.19). Thus, an axis fitted without exposure data recovered the expected ordering of adverse physiology.

The components differed sharply (Fig. 4a). Current smoking was associated primarily with alignment: 0.72 standard deviations (0.67 to 0.77) versus 0.15 (0.10 to 0.20) for magnitude. After socioeconomic adjustment, the magnitude association disappeared (−0.003, −0.06 to 0.05) while alignment remained 0.62 (0.57 to 0.67); after additional adjustment for the other exposures, magnitude was −0.10 (−0.17 to −0.03) and alignment 0.57 (0.50 to 0.63) (Fig. 4b). Physical inactivity showed the reverse ordering, with a stronger association with magnitude than alignment (0.36 versus 0.24), while poorer diet was weakly associated with both. Former smokers were close to never-smokers on both quantities (0.09 standard deviations), suggesting that the alignment-dominant pattern was specific to current smoking.

**Figure 4.**
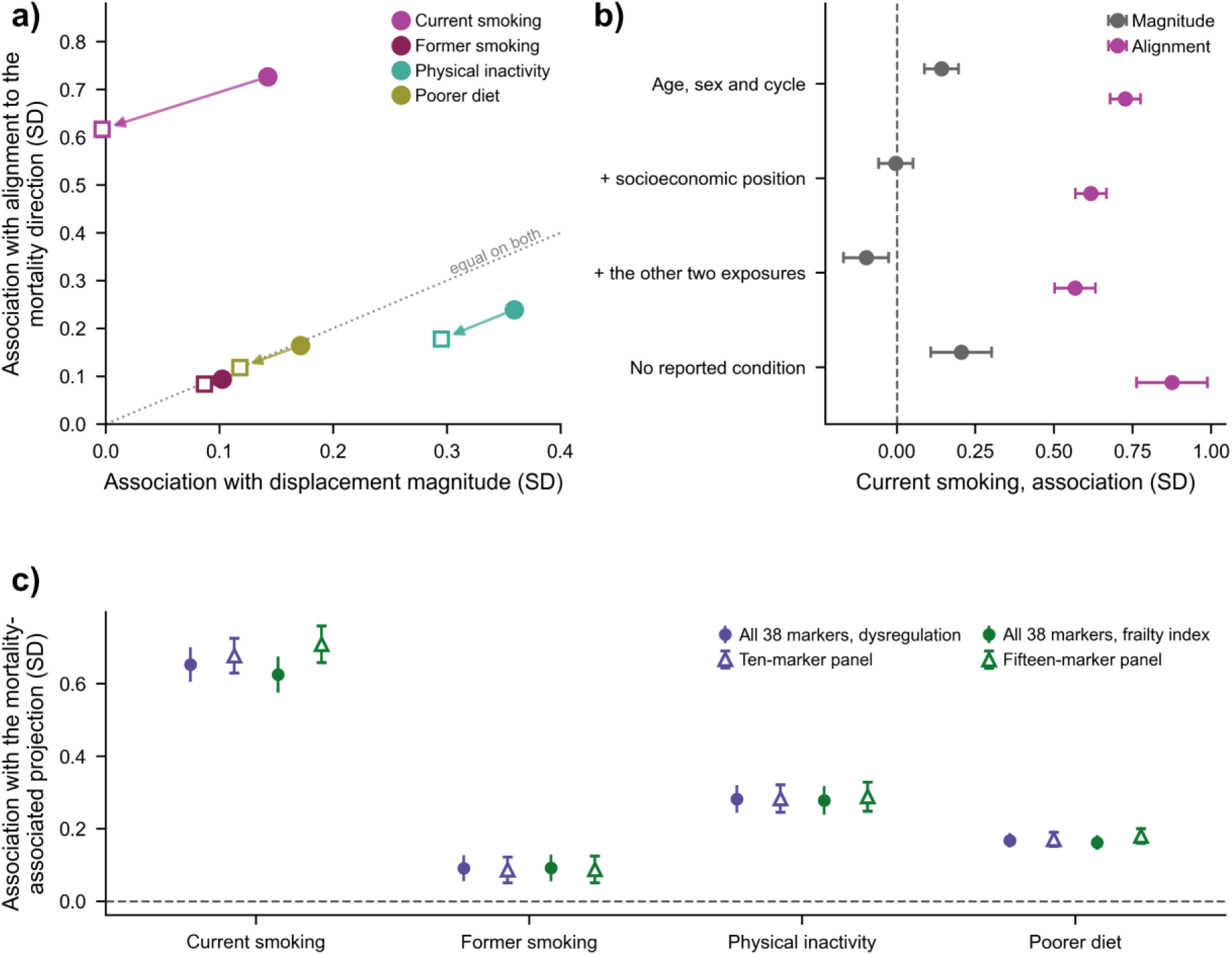
An axis fitted to mortality alone recovered three exposures it had never seen, and separated them by which component of displacement they moved. Smoking, physical inactivity and diet took no part in biomarker selection, in estimation of the mortality direction, or in the choice of panel size or identity. Colours denote exposures throughout and are used for nothing else in this paper: magenta for current smoking, dark wine for former smoking, teal for physical inactivity, olive for poorer diet. Associations are in standard deviations of the analysis population, among 11,497 participants aged 52–79 years. **a**, Each exposure placed on the plane of the two quantities the paper distinguishes: association with displacement magnitude on the horizontal axis, association with alignment to the mortality direction on the vertical. Filled circles are estimates adjusted for age, sex and survey cycle; open squares are the same estimates after further adjustment for socioeconomic position, and the arrow runs between them. The dotted diagonal marks equal association with both quantities; distance above it measures how far an exposure acts on direction rather than on distance. Current smoking sits far above the diagonal, physical inactivity below it, poorer diet on it. **b**, Current smoking across a nested adjustment cascade, and within the 1,950 participants reporting no chronic condition. Grey is the association with magnitude, magenta with alignment. Alignment remains substantially larger than magnitude after socioeconomic and mutual adjustment, although it is attenuated (0.72 to 0.62 to 0.57), and is larger in participants reporting no condition; the magnitude association is abolished by socioeconomic adjustment. **c**, The reduced panels of Table 2 carry the same exposure signatures as the full 38 markers. Filled circles are the full panel, open triangles the reduced panel, in homeostatic-dysregulation (purple) and sensitised frailty-index (green) representations. Points are estimates and bars are 95% confidence intervals, computed with heteroskedasticity-consistent (HC3) standard errors. The adjustment arrow in **a** begins from a model fitted on the same participants as the socioeconomically adjusted model, so it isolates adjustment from any change of sample. All associations are cross-sectional and are not intervention effects.

Prevalent disease did not explain the smoking pattern. Among 1,950 participants reporting no chronic condition, the smoking–alignment association was larger, not smaller, at 0.88 (0.76 to 0.99) (Fig. 4b).

Both reduced panels preserved these associations (Fig. 4c). For current smoking, projection was 0.68 (0.63 to 0.73) for the ten-marker panel and 0.71 (0.66 to 0.76) for the fifteen-marker panel, versus 0.65 for all 38 markers; estimates for physical inactivity and diet were similarly close to the full panel.

Five of six predictions were confirmed; physical inactivity was more magnitude-dominant than predicted. Overall, the mortality-derived axis distinguished exposures that were not used to fit it: smoking was associated mainly with orientation, inactivity more with total displacement, and diet weakly with both. These cross-sectional associations support construct validity but are not intervention effects.

### 6. The Finding Replicated in an Independent Cohort Under Predictions Registered in Advance

We repeated the analysis in NHANES III (1988–1994), a disjoint cohort measured on different assay platforms and linked to mortality through 2019. Eight predictions and falsification criteria were signed off before any mortality-linked analysis. The young-healthy reference, transformations and mortality direction were rebuilt within NHANES III, so replication tested the physiological pattern rather than transport of a fitted model.

The replication cohort comprised 13,195 adults aged 18–79 years with complete data on 34 of the 38 markers, of whom 4,474 were aged 52 years and above with 3,401 disease deaths and a median follow-up of 26.7 years.

Seven of eight predictions were confirmed. Women again reported more morbidity but had lower disease mortality (female hazard ratio 0.714, 0.678 to 0.753). They were not less displaced than men (+0.005 and +0.070 standard deviations) but were markedly lower in alignment (−0.526 and −0.594) and projection (−0.453 and −0.438). Magnitude adjustment again moved the female hazard ratio away from unity. A median 59.6% of positive mortality-associated contribution lay within conventional clinical limits. The eight locked markers available in NHANES III reproduced the 34-marker projection at 0.849, above the 95th percentile of 200 random eight-marker panels and at 95.7% of the ceiling achieved by eight markers selected natively within NHANES III; five locked markers were recovered in every native-selection fold (Fig. 5a–d, Table 3).

**Figure 5.**
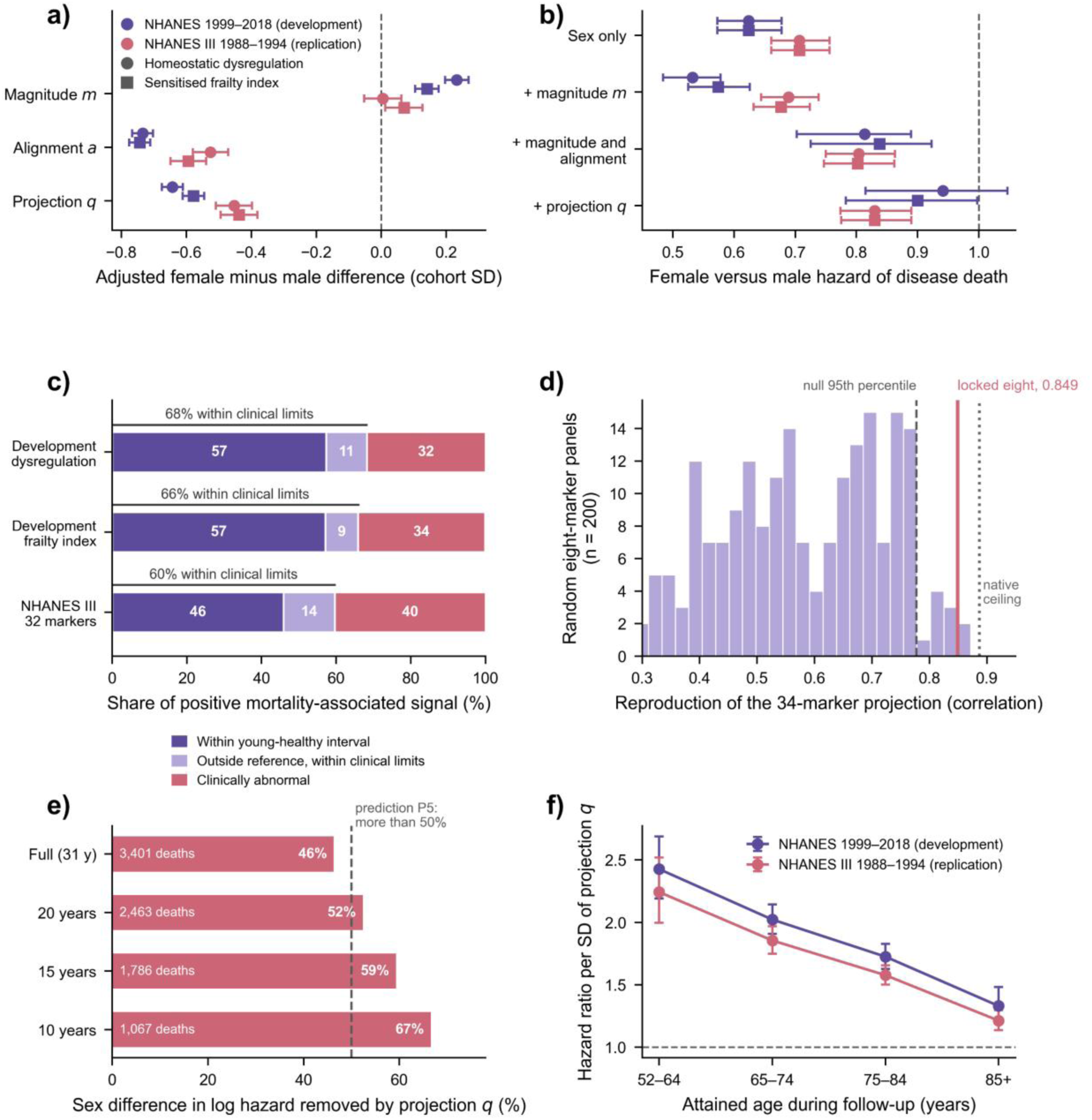
Seven of eight predictions registered before analysis were confirmed in an independent cohort measured three decades earlier. Throughout this figure, colour distinguishes cohort — purple for the NHANES 1999–2018 development cohort, rose for the NHANES III 1988–1994 replication cohort — and marker shape distinguishes representation, circles for homeostatic dysregulation and squares for the sensitised frailty index. **a**, Adjusted female-minus-male differences in displacement magnitude, alignment and projection, in standard deviations of the analysis population, in both cohorts and both representations. Magnitude straddles zero in the replication cohort while alignment and projection are strongly negative. **b**, Female versus male hazard of disease death across the same nested adjustment cascade in both cohorts. **c**, Where positive mortality-associated signal sits relative to conventional clinical thresholds, in both cohorts. The bracket above each bar gives the combined share falling within clinical limits. **d**, Whether the locked panel transported. The histogram is the distribution of reproduction achieved by 200 eight-marker panels drawn at random from the 34 markers available in NHANES III. The dashed line marks the 95th percentile of that null, the solid rose line the locked eight, and the dotted line the ceiling set by eight markers selected natively within NHANES III. **e**, Pre-specified follow-up truncation: the percentage of the sex difference in log hazard removed by adjustment for projection at each horizon, with the number of deaths contributing shown inside each bar. The dashed line marks the 50% threshold that prediction P5 required. **f**, The measured reason P5 fell short: the hazard ratio per standard deviation of projection within bands of attained age during follow-up, in both cohorts. Points are estimates and bars are 95% confidence intervals. The mortality direction, young-healthy reference and transformations were all rebuilt within NHANES III rather than imported. Predictions, criteria and verdicts are set out in Table 3.

**Table 3.** NHANES III prediction register (P1–P7b): predictions, falsification criteria and verdicts. The specification recording these predictions, their criteria, and the conditions that would have counted as a failure to replicate was signed off before any mortality-linked analysis was run in NHANES III and is archived unmodified with the analysis code. Two figures are reported where both representations were evaluated: homeostatic dysregulation first, sensitised frailty index second. Estimation populations differ by prediction and are stated with each: P1 is estimated in the broad age 18–79 cohort (n = 16,040; 5,747 deaths), while P2 to P7b are estimated in the 34-marker complete-case cohort aged 52 years and above (n = 4,474; 3,401 disease deaths). P5 is the one prediction not met on its stated criterion. The shortfall is measured rather than asserted: the projection association declines with attained age in both cohorts (Fig. 5f), 46% of person-time in NHANES III accrues above age 75, and truncating follow-up at 20 years — a sensitivity specified in advance rather than chosen after seeing the result — returns the figure to 52%. No prediction was recorded about the size of the projection association per standard deviation, because longer follow-up and an older attained-age distribution were expected in advance to attenuate it. Eight markers rather than ten are evaluated in P7: neutrophil count exists in NHANES III only as total granulocytes, and gamma-glutamyl transferase is 48% complete in Phase 1. Both were excluded rather than substituted, a decision fixed in the specification. Reported reproduction is a held-out correlation against the 34-marker mortality-directed displacement estimated within NHANES III itself; the null draws 200 random eight-marker panels from the same 34.

| No. | Prediction, recorded before any mortality-linked analysis | Criterion fixed in advance | Observed in NHANES III | Verdict |
| --- | --- | --- | --- | --- |
| P1 | Women will have lower disease mortality than men | Age-adjusted female hazard ratio below 1, interval excluding 1 | 0.714 (0.678–0.753) | Confirmed |
| P2 | Women will not be less physiologically displaced than men | Adjusted female–male difference in magnitude $m$ at or above 0 | +0.005 (–0.053 to +0.062) and +0.070 (+0.012 to +0.127) | Confirmed |
| P3 | Women will be lower in alignment and in projection | Both differences negative, intervals excluding 0 | $a$ –0.526 (–0.580 to –0.471); $q$ –0.453 (–0.509 to –0.398) | Confirmed |
| P4 | Adjusting for magnitude will not attenuate the female advantage | Hazard ratio moves away from 1, or is materially unchanged | 0.707 (0.661–0.756) → 0.689 (0.644–0.738) | Confirmed |
| P5 | Adjusting for projection will move the hazard ratio toward 1 | More than half the sex difference in log hazard removed | 46.3% at full follow-up; 52.4% at the pre-specified 20-year horizon | Partially met |
| P6 | Most positive mortality-associated signal will sit within conventional clinical limits | More than 50% within limits | 59.6% | Confirmed |
| P7 | The locked eight markers will transport | Reproduction above the 95th percentile of a random eight-marker null | 0.8488 against a null 95th percentile of 0.7774 (99th percentile) | Confirmed |
| P7b | The locked eight will approach a natively selected panel | At least 90% of the reproduction attained by eight markers selected within NHANES III | 95.7% of a ceiling of 0.8868 | Confirmed |

Prediction P5 was partially met. At full follow-up, projection adjustment attenuated 46% of the sex difference in log hazard rather than the prespecified >50%. The projection–mortality association declined with attained age in both cohorts — from 2.243 at ages 52–65 to 1.213 above age 85 in NHANES III, versus 2.427 to 1.331 in development (Fig. 5f). Because 46% of NHANES III person-time accrued above age 75, the longer follow-up placed more weight where the axis was weaker. The prespecified 20-year truncation restored attenuation to 52%; 15- and 10-year horizons gave 59% and 67%, respectively (Fig. 5e). The central contrast therefore replicated, but its average magnitude depended on attained-age distribution and follow-up horizon.

### 7. Retrospective Sorting Produced Two Candidate Trial Groups With Opposite Sex Composition

For event-driven lifespan trials, recruitment benefits from participants likely to accrue events within a feasible horizon. A healthspan-oriented trial may instead seek participants with measurable physiological disturbance but low near-term mortality. We therefore asked whether the projection– orthogonal plane could retrospectively identify candidate groups with these contrasting profiles.

Participants aged 52–79 years were assigned to a lifespan candidate group if projection was at or above the training 90th percentile, and to a physiologically displaced, lower-risk group if projection was at or below the 60th percentile and orthogonal displacement at or above the 60th. Thresholds were estimated in training cycles and applied unchanged to held-out cycles across 252 directed partitions. Sex was not used to choose thresholds, panels or representations.

The lifespan group contained about 10% of eligible participants and had a five-year disease-mortality rate 3.7 to 4.3 times the cohort rate. The physiologically displaced group contained 13% to 17% and had 0.44 to 0.50 times the cohort rate while remaining displaced across a median of three to five physiological systems. A usable rule was obtained in every partition (Fig. 6a–b).

**Figure 6.**
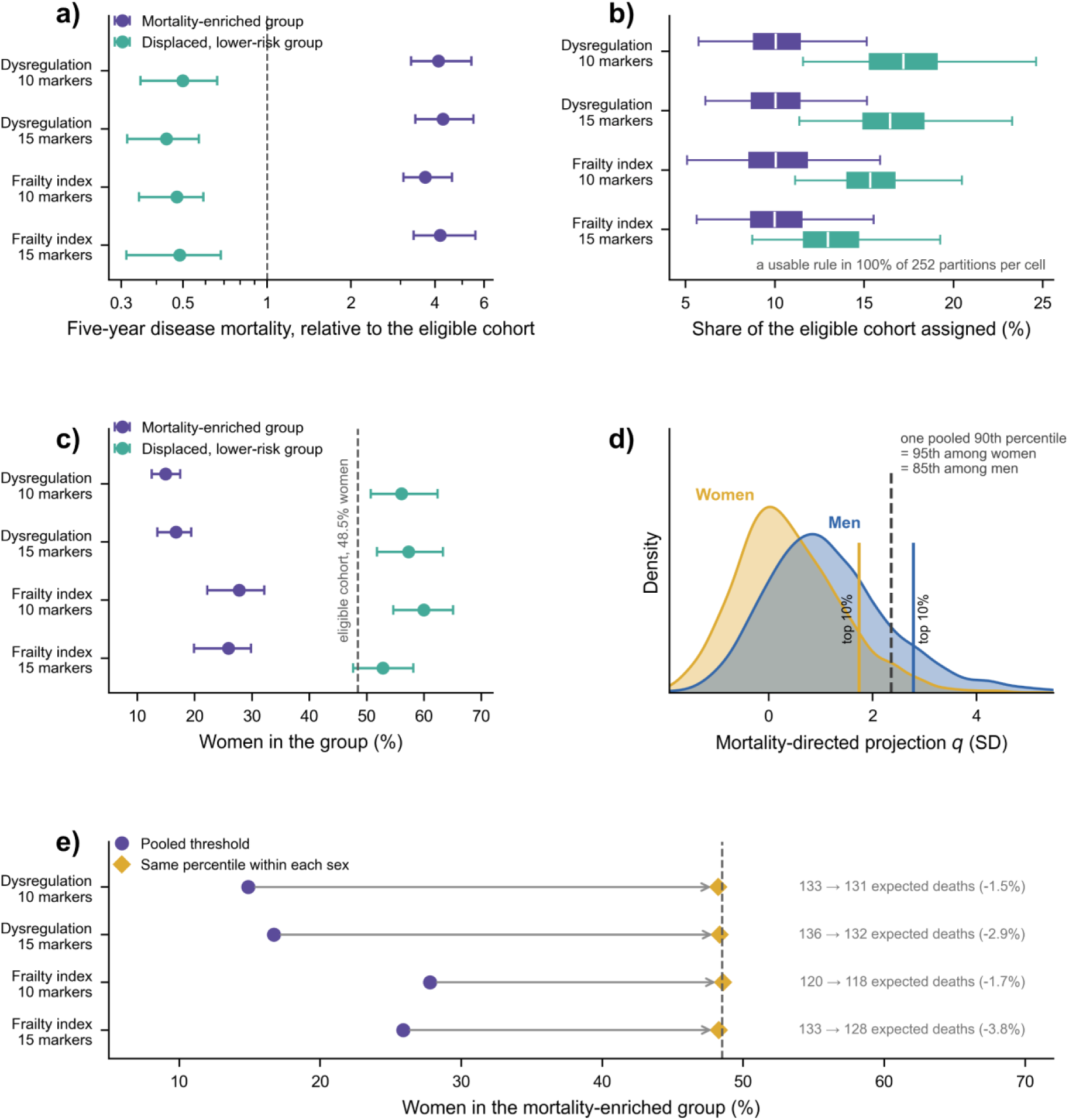
One geometric rule produced two candidate recruitment groups of opposite sex composition without using sex. Four configurations are shown throughout: each representation — homeostatic dysregulation and the sensitised frailty index — at panel sizes of ten and fifteen markers. Purple denotes the mortality-enriched group, teal the physiologically displaced, lower-risk group. Neither is a trial arm: both are retrospectively sorted observational groups, and no healthspan or functional outcome was measured in either cohort. **a,** Five-year disease mortality within each group relative to the eligible cohort, on a logarithmic scale. The mortality-enriched group is enriched roughly fourfold; the displaced, lower-risk group is actively depleted of near-term death. The dashed line marks parity with the cohort. **b,** The share of the eligible cohort assigned to each group, across all 252 directed survey-cycle partitions. Boxes span the interquartile range, the white line is the median, and whiskers extend to 1.5 times the interquartile range; outlying partitions are not drawn. **c,** The percentage of each group that was women, with the eligible cohort’s own composition marked by the dashed line. Sex was measured after assignment and was never used to select or adjust a threshold, a panel or a representation, so this composition follows from the geometry alone. **d,** Why a pooled threshold produces a male-dominated group. The held-out mortality-directed projection is shown separately for women (yellow) and men (blue) in the eligible cohort. Solid vertical lines mark the 90th percentile *within* each sex, each admitting exactly the top tenth of that sex. The dashed line is the single pooled 90th percentile the primary rule applies, and it is not the same percentile in the two distributions. This panel is drawn on the projection defined by all 38 markers — the geometry quantified in Fig. 2 — rather than on a locked-panel rule, because the reduced-panel projections are not written per participant; the locked-panel rules place the pooled cut at the 84th to 86th percentile among men and the 94th to 97th among women, closely matching the values shown. **e**, A pooled threshold compared with the same percentile applied within each sex. Arrows run from the pooled result (purple circles) to the within-sex result (yellow diamonds); expected five-year events under each rule, and the percentage change, are given at the right of the panel. Participants aged 52–79 years were assigned to the mortality-enriched group if projection reached the training 90th percentile, and to the displaced, lower-risk group if projection was at or below the 60th percentile while orthogonal displacement was at or above the 60th. Thresholds were estimated within training cycles and applied unchanged to held-out cycles. Intervals in **a** and **c** are 2.5th to 97.5th percentiles across partitions. The within-sex threshold in **e** is a post-hoc variant reported as a sensitivity, not a pre-specified rule, and is offered as a design option to be pre-specified in a future study rather than as a validated screen.

The groups had opposite sex composition. In a cohort that was 48.5% women, the lifespan group was 15% to 28% women and the physiologically displaced group 53% to 60% (Fig. 6c). Because sex was not used in assignment, the imbalance arose from the geometry itself. Configurations that resolved the sex difference most strongly produced the most male lifespan groups (14.9% and 16.7% women), whereas weaker configurations produced 25.9% and 27.8% women.

The imbalance reflected the pooled threshold rather than poor performance in women. Selected women had a five-year disease-mortality rate of 81.0 per 1,000 person-years versus 51.9 in selected men, corresponding to 8.2-fold enrichment over cohort women and 3.1-fold over cohort men. The pooled rule selected fewer women, but those selected were at very high risk.

The mechanism is arithmetic. Because women sat lower on the projection distribution, the pooled 90th percentile corresponded to the 84th–86th percentile in men but the 94th–97th in women, admitting 13.9%–16.4% of men and only 3.1%–5.8% of women (Fig. 6d). A threshold that never explicitly uses sex can therefore act as a sex filter when the underlying score differs by sex.

Applying the same percentile within each sex — a post-hoc sensitivity analysis — restored the cohort sex composition: 48.2%–48.6% women at the same total group size. Expected five-year events decreased only modestly, from 133 to 131, 136 to 132, 120 to 118 and 133 to 128 across the four configurations (1.5%–3.8%) (Fig. 6e). Thus, in this cohort, sex balance could be restored with a small loss of expected events, although the rule requires prospective validation.

### 8. C-Reactive Protein and Fasting Glycaemic Measures Added Little Beyond the Panel, Whereas FEV₁ Added Prognostic Information

The 38-marker matrix came from routine blood chemistry, haematology and urinalysis and contains no direct measure of organ mechanical capacity. We therefore asked two complementary questions: which commonly used assays add little incremental prognostic information once the panel is present, and which functional measurements capture information the blood panel misses?

In the development cohort, three candidates were specified before results were examined: high-sensitivity C-reactive protein; fasting glucose and insulin with HOMA-IR; and spirometric FEV₁. Four other candidates — cardiorespiratory fitness, grip strength, knee-extensor strength and apolipoprotein B — were excluded for prespecified availability or redundancy reasons (Methods).

Each retained candidate was evaluated in its eligible cycles on identical participant rows, against cross-fitted projections from both the full 38-marker panel and the ten-marker panel. Incremental information was assessed by change in held-out partial log-likelihood, with conditional hazard ratios reported alongside.

C-reactive protein was strongly associated with disease mortality on its own (hazard ratio 1.31 per standard deviation, 1.23 to 1.38; n = 8,886; 2,204 deaths) but showed no incremental association conditional on the panel: 0.98 (0.93 to 1.04) given all 38 markers and 1.01 given the ten, with held-out likelihood decreasing under both comparators (Fig. 7). Fasting insulin and HOMA-IR were similarly redundant in these data. Fasting glucose added no information beyond all 38 markers but retained a small association given the ten-marker panel (1.06, 1.02 to 1.11; held-out likelihood +3.4; six of nine folds improved), consistent with the omission of glycated haemoglobin from the reduced panel. This effect weakened under a stricter eight-hour fasting definition and is treated as suggestive. By contrast, reduced FEV₁ remained strongly associated with mortality conditional on both panels — 1.57 (1.34 to 1.83) given all 38 markers and 1.63 (1.39 to 1.90) given the ten — and improved held-out likelihood in all three available folds (Fig. 7).

**Figure 7.**
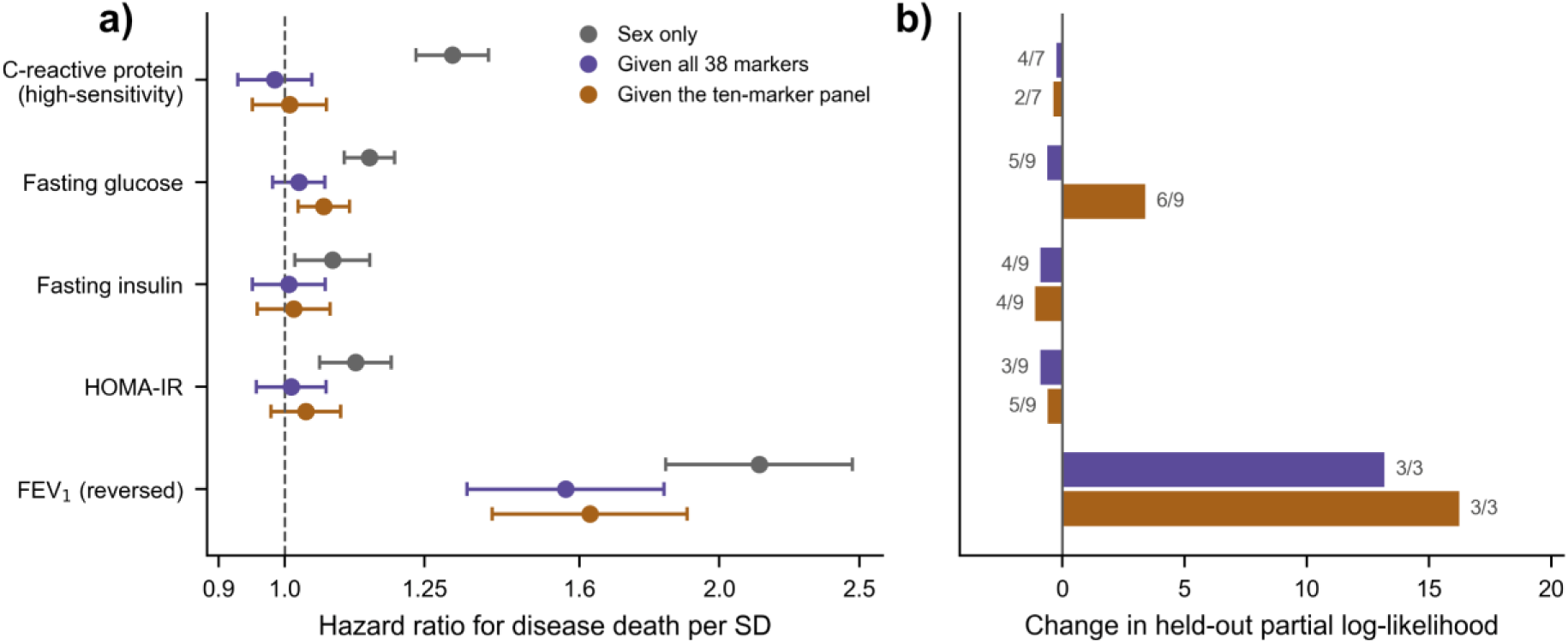
What the routine panel already captures, and what it misses (development cohort). Each candidate measurement was evaluated in its own eligible NHANES cycles, since no common complete-case cohort exists across them, and always on identical rows: models with and without the candidate were never fitted on different participants. Candidates are oriented so that higher values are more adverse; FEV₁ is therefore reversed. C-reactive protein and insulin are log-transformed, and C-reactive protein is standardised within assay era because the measurement changed between 2009–2010 and 2015–2016, with no data in 2011–2014. Spirometry is restricted to quality grades A and B and adjusted for standing height. **a**, Hazard ratio per standard deviation, with 95% confidence intervals, for each candidate under three specifications: sex only (grey), given the cross-fitted projection of all 38 markers (purple), and given the ten-marker panel projection (ochre). The dashed line marks no association. The grey estimates are a reference quantity, not evidence: a measurement can be strongly associated with mortality on its own and add nothing once the panel is present, which is precisely what C-reactive protein does. The question this figure asks is whether the coloured estimates remain away from 1. **b**, Change in held-out partial log-likelihood when the candidate is added, summed across leave-one-cycle-out folds within that candidate’s eligible cycles. Positive values indicate improved prediction in cycles not used for fitting. Labels give the number of folds improved out of the number available. Folds differ by candidate — seven for C-reactive protein, nine for the fasting measures, three for FEV₁ — and results are not pooled across candidates for that reason.

NHANES III provided a larger exploratory test of functional measurements. Conditional on the cross-fitted 34-marker projection, reduced FEV₁ had a hazard ratio of 1.21 per standard deviation (1.16 to 1.27) and improved held-out partial log-likelihood by 31.3 in 8 of 10 folds; forced vital capacity followed at 1.16 (1.10 to 1.21) and +15.0. Bioimpedance reactance added modestly (1.07, 1.03 to 1.11; +5.2) and tandem-stand time marginally (1.05, 1.01 to 1.09; +2.0). Femoral-neck bone mineral density added little (1.04, 1.00 to 1.08; +1.5), while total bone mineral content and bioimpedance resistance showed no incremental information (Fig. 8).

**Figure 8.**
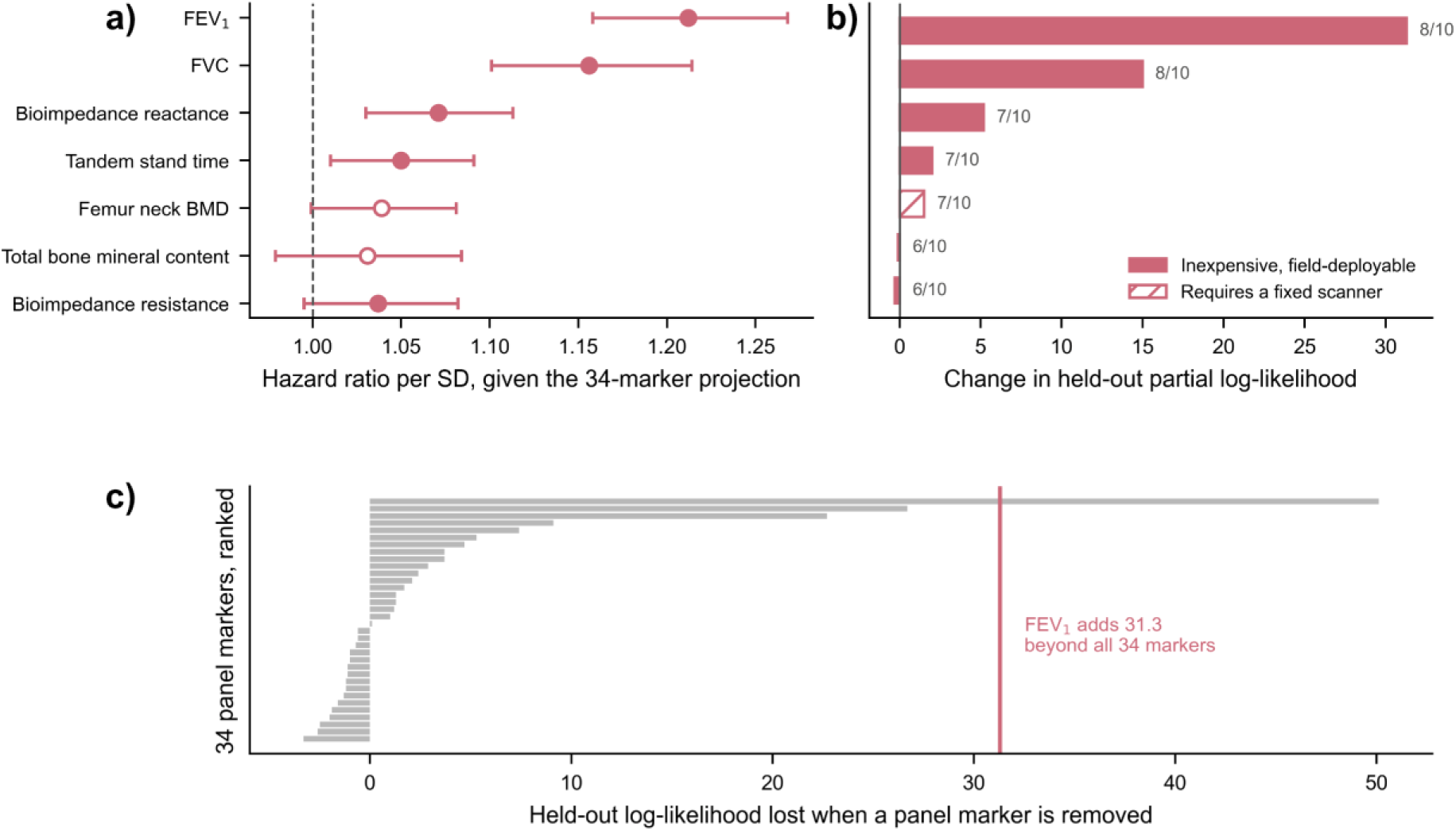
One inexpensive lung-function measurement carried more independent mortality information than almost every marker in the panel (replication cohort). Every measurement in this figure comes from the replication cohort, so rose carries its usual meaning throughout and deployability is shown by fill instead: solid markers and bars are measurements that are inexpensive and field-deployable, open and hatched ones require a fixed scanner. a, Hazard ratio for disease death per standard deviation of each candidate functional measurement, estimated with the 34-marker mortality-directed projection already in the model, so each estimate is the information the measurement adds beyond the routine panel rather than its association on its own. Points are estimates, bars are 95% confidence intervals, and the dashed line marks a hazard ratio of 1. b, Change in held-out partial log-likelihood when each candidate is added to the panel projection. Values to the right of zero indicate added information. The figure at the end of each bar is the number of cross-validation folds out of ten in which that candidate improved held-out fit, a stability check that a single pooled statistic cannot provide. c, The same quantity computed for the panel itself: held-out log-likelihood lost when each of the 34 markers is removed in turn, ranked. The vertical rose line marks the gain contributed by forced expiratory volume in one second beyond the entire panel, placing it above all but one individual marker. Analyses in this figure are exploratory: they were specified after the primary replication was complete, carry no registered prediction, and are reported as hypothesis-generating. Candidate measurements differ in availability, so the number of participants and deaths contributing differs between candidates and is reported for each in Supplementary Information.

Against the panel itself, the spirometry result was large. Removing each of the 34 markers in turn reduced held-out log-likelihood by a median of 0.6 units; the strongest marker, red cell distribution width, cost 50.1. FEV₁ added 31.3 units on top of the complete panel — more incremental information than 33 of the 34 individual markers contributed when removed (Fig. 8c).

Across two cohorts, the same pattern emerged: measurements from domains already represented in the blood panel, such as inflammation and glycaemia, contributed little incremental information, whereas a direct measure of pulmonary function contributed substantially. This pattern is consistent with, but does not establish, a distinction between measured physiological state and functional capacity: little incremental prediction shows conditional redundancy for this endpoint in this sample, not that the panel biologically represents inflammation or glycaemia. A sparse panel need not include each system’s most familiar assay, but it may benefit from direct functional measurements that blood cannot replace.

## Discussion

### More Informative Shots at Goal

Geroscience needs more informative shots at goal if we hope to meaningfully extend the human healthspan and lifespan. While the field contains many promising interventions, each definitive trial is expensive, slow, and vulnerable to failure through heterogeneous recruitment, low event rates, poorly matched endpoints, and biomarker batteries that are either too shallow to capture consequential physiology or too burdensome for broad deployment. The practical objective of this study was to determine whether inexpensive, routinely available measurements could be organised more strategically — to reduce the cost per informative participant, allow more promising interventions to proceed to trial, and do so for both sexes at once.

The main result is that total physiological displacement and its direction carry different information. Three observations support this conclusion. First, most positive mortality-associated signal occurred while individual biomarkers remained within conventional clinical limits. Second, women were not closer to young-health than men, but their displacement was substantially less aligned with the mortality-associated direction; projection attenuated most of the female–male log-hazard difference, whereas magnitude attenuated none. Third, this directional structure replicated across cohorts, compressed to a small routine panel and defined candidate groups with distinct mortality profiles.

### Direction, Not Distance

Both magnitude measures — the sensitised frailty index and homeostatic dysregulation — strongly predicted mortality, yet neither accounted for the female survival advantage. Re-expressing the same 38 measurements as displacement along a mortality-associated direction attenuated most of that difference. The point is not that distance is uninformative; it is that distance and direction answer different questions. A scalar measure of total dysregulation can be prognostic while still discarding the configuration that distinguishes two people — or two groups — at the same overall level of disturbance.

This distinction also changes how existing ageing instruments should be described. Frailty indices ^12–14^, biological-age clocks ^15,16^, allostatic load ^17^ and homeostatic dysregulation ^18^ are not mathematically equivalent, and not all are literal distance measures. What they share is scalar compression of a multidimensional physiological state. Our results suggest that this common compression can create a common blind spot: different configurations can map to similar scalar values while carrying different associations with mortality.

### Where the Signal Sits

Approximately two thirds of positive mortality-associated signal arose while raw biomarker values remained within conventional clinical limits, and more than half while they remained within the same-sex young-healthy interval. This does not mean that clinical reference intervals are wrong; they answer a different question, namely when a value is sufficiently unusual or disease-associated to warrant clinical attention. Nor does it make the young-healthy centre a treatment target. It shows that, at population level, continuous variation within the ‘normal’ region carries a large share of the total mortality-associated contribution, which threshold-based scoring discards. The share follows in part from how many measurements fall in that region, so it does not imply that a normal-range value is individually as informative as an abnormal one.

### What the Sex Finding Does — and Does Not — Mean

The sex result locates a difference within measured physiology; it does not identify its cause. The fitted direction is predictive and observational. It may reflect causal ageing processes, accumulated disease, treatment, exposure history, preclinical illness, physiological compensation, or unmeasured confounding. Lower alignment is not synonymous with health: orthogonal displacement may still contribute to morbidity, disability or mortality through pathways absent from this model. The supported claim is narrower: within these measured systems, women and men share a similar mortality-associated direction but occupy different positions along it.

### A Shared Axis Can Accommodate Biological and Behavioural Contributions

Biological and behavioural accounts of the female survival advantage need not be mutually exclusive. A female advantage persists under extreme mortality and is especially marked in infancy, where adult health behaviour cannot have a causal role ^11^. Yet the size of the gap changed markedly during the twentieth century, tracking sex differences in smoking and other exposures ^29,30^. A shared mortality-associated axis can accommodate both observations: baseline physiology may shift the distributions, while modifiable exposures move individuals along the same axis.

Current smoking was consistent with this interpretation: it was strongly associated with alignment but had essentially no residual association with magnitude after socioeconomic adjustment. In the full cohort, adjustment for smoking, alcohol and physical activity changed the female hazard ratio only from 0.64 to 0.63, and the survival advantage persisted in behaviourally restricted strata. These data therefore support a model in which modifiable exposures can alter position along the shared axis without accounting for the entire sex offset. Whether an intervention can move an individual away from the mortality-associated direction — and whether doing so changes clinical risk — remains a prospective experimental question.

### Replication Across an Assay Generation

Replication in NHANES III argues against the pattern being unique to one survey era. The young-healthy reference, transformations and mortality direction were rebuilt on a different assay generation, yet the sex configuration, signal location and attained-age gradient were reproduced under predictions fixed before mortality-linked analysis. The one partially met prediction was informative: full-follow-up attenuation was smaller because much more person-time occurred at older attained ages, where the projection–mortality association was weaker. The exact ten-marker identity, however, remains a development product. Eight markers transported well and approached a natively selected ceiling, but the final combination requires prospective and geographically external validation.

### Redundant Assays and Missing Capacity

The candidate analyses separate biological relevance from incremental information. High-sensitivity C-reactive protein was strongly prognostic alone but contributed no detectable held-out information beyond either routine panel in these data; fasting insulin and HOMA-IR behaved similarly. Fasting glucose added little beyond the full panel but recovered a small amount of information omitted by the ten-marker panel, which lacks glycated haemoglobin. This is a useful limitation rather than a nuisance result: reducing a panel changes what it can represent, which informs future panel design.

Direct functional measurements may meaningfully improve future panel performance. For example, reduced FEV₁ remained strongly associated with mortality after conditioning on the complete routine matrix in the development cohort and added more held-out prognostic information than 33 of 34 individual panel markers in NHANES III. This extends prospective evidence linking impaired pulmonary function and lower grip strength to mortality ^31–34^. Circulating biomarkers report regulation, inflammation, organ stress, turnover and damage; they do not necessarily measure integrated functional capacity. Blood can indicate pulmonary strain, but it cannot directly measure how much air a person can move.

The emergent design principle is therefore information density; not minimal marker counts or one assay per organ label. The most useful battery is the least burdensome combination that preserves the physiology required for the intended experiment. Here, routine blood and urine measurements captured much of the mortality-associated state, while spirometry supplied a complementary measure of capacity. The weak incremental value of the fixed-scanner measurements also suggests that cost and technological complexity need not track information gain.

### One Geometry, Two Trial Objectives

The two candidate trial groups address different recruitment objectives. The mortality-enriched group contained about 10% of eligible participants at 3.7–4.3 times the cohort’s five-year disease-mortality rate. The physiologically displaced group contained 13%–17% at 0.44–0.50 times the cohort rate while remaining displaced across several physiological systems. The second group is therefore better described as a candidate healthspan-enrichment population, not as proof that orthogonal displacement is benign or that it predicts functional decline.

Prognostic enrichment can reduce the number of participants or follow-up time needed to accrue a target number of events ^5^, but the gain does not scale mechanically as the inverse of the observed event-rate ratio: formal efficiency depends on treatment effect, censoring, recruitment rate, screening burden and generalisability. More importantly, prognostic enrichment is not predictive enrichment. Participants at high risk may not share an intervention’s target mechanism, their risk may not be reversible, and selection need not increase treatment effect. The central untested assumption is therefore whether this geometry identifies people whose outcomes are more modifiable by a given intervention. The lower-risk, physiologically displaced group poses the complementary question: whether orthogonal displacement predicts functional decline or recoverable reserve requires healthspan outcomes not available in NHANES.

### Better Enrichment Can Worsen Sex Imbalance — Unless Thresholds Account for It

Without using sex in assignment, the pooled rule produced strongly sex-skewed groups: the mortality-enriched group was 15%–28% women, whereas the physiologically displaced group was 53%–60% women. This imbalance follows directly from the sex offset on the projection axis.

A pooled high-risk threshold selects proportionally fewer women because women sit lower on the mortality-associated projection. The effect was strongest for configurations that resolved the sex difference most clearly: the better the score separated the sex distributions, the more male the mortality-enriched group became. This principle extends beyond this panel. Any prognostic enrichment score that differs systematically by sex can create an implicit sex filter when a single pooled threshold is used.

That does not mean the score performed poorly in women. Under the pooled rule, selected women had higher five-year disease mortality than selected men (81.0 versus 51.9 per 1,000 person-years) and greater enrichment over their sex-specific cohort baseline (8.2-fold versus 3.1-fold). The rule selected fewer women because it sampled a more extreme percentile of the female distribution.

A simple post-hoc sensitivity analysis showed how the imbalance could be managed. Applying the same 90th percentile within each sex produced groups that were 48.2%–48.6% women at the same total size, while expected five-year events fell by only 1.5%–3.8% across configurations. The trade-off is explicit: women enter at lower absolute risk and men at higher absolute risk than under the pooled rule, so baseline risk is not exchangeable across sex. A future trial using this strategy should therefore prespecify sex-stratified randomisation and sex-by-treatment interaction analyses. Because this analysis was post hoc, it is a design option, not a validated recruitment rule.

The diagnostic is simple: report where any proposed pooled cutoff falls within the score distribution of each sex. Here, the nominal pooled 90th percentile was only the 84th–86th percentile in men but the 94th–97th in women. Within-sex percentiles make the recruitment trade-off visible and controllable rather than allowing the score distribution to determine sex composition implicitly. Equal percentiles are only one option; thresholds can be chosen separately to target a prespecified balance between event yield, generalisability, and sex representation.

Two constraints remain. First, within-sex selection produced different absolute baseline risks in women and men (43.1 versus 67.3 per 1,000 person-years), so randomisation and analysis must account for sex. Second, the total screening burden is not reduced by balancing the selected group; the same number of candidates must still be measured.

The important reassurance is that sex-balanced recruitment does not require separate predictive axes. A direction fitted in one sex retained 98.6%–104.7% of its mortality association in the other, and sex-specific directions did not improve held-out fit. The data therefore support one substantially shared measured axis with sex-specific position, not proof of one universal ageing biology. Projection was also more strongly associated with mortality in women than men (2.07 versus 1.76 per standard deviation), providing no empirical basis here for treating women as a noisier or less informative trial population.

### Boundaries

Several limitations bound the interpretation. Both cohorts are observational, so none of the reported associations establishes causality or mediation; in particular, adjustment for a mortality-supervised physiological state that may lie downstream of sex changes a coefficient but does not estimate mediation or the proportion of the survival advantage explained. Because the direction analysis conditions on survival and examination at age 52 or older, its sex contrast pertains to survivors eligible for later-life recruitment and may be shaped by differential survival before entry; it should not be read as explaining the lifetime sex–mortality gap. The physiological analyses are also complete-case analyses: if panel completeness depends jointly on sex and mortality-related health, selection into the analytic cohort could alter both the observed sex contrast and its attenuation after adjustment for projection, and the direction and size of that bias are unknown. Development and replication come from the same national survey programme; replication in another country and health system is still needed. The reduced panel was selected by one algorithm, its binding adequacy criterion passed narrowly (0.9018 versus 0.90), and panels of eight and nine markers were not evaluated. Physiology was measured once, so the geometry describes cross-sectional position rather than within-person trajectory. NHANES records sex as a binary variable and cannot resolve chromosomal, gonadal, hormonal, reproductive or gender-related contributors. Most importantly, prognostic association is not surrogate validity or treatment-effect prediction ^35^: neither the panel nor the sorting rule has been shown prospectively to identify responders, increase treatment effects, reduce trial cost or improve clinical outcomes.

## Conclusion

Geroscience cannot test every plausible intervention with broad, expensive trials. More informative trials require extracting more value from inexpensive measurements and matching recruitment to the outcome a study is designed to observe.

Total displacement describes how much measured physiology differs from young-health; mortality-directed projection describes how much of that displacement follows a multivariable pattern associated with subsequent disease death. Much of this signal lay within conventional clinical limits. Women were not less dysregulated than men, but less of their displacement lay along a substantially shared mortality-associated direction, and this projection attenuated most of the female–male log-hazard difference where magnitude did not. Ten routine measurements preserved most of the directional information, the central geometry replicated across survey eras, and spirometry added a complementary measure of physiological capacity.

The practical hypothesis is now testable: a small routine pathology panel, brief functional testing and thresholds prespecified within sex may support mortality-enriched or physiologically displaced recruitment while maintaining sex balance. In this retrospective analysis, balancing the mortality-enriched group cost only 1.5%–3.8% of expected five-year events. The next step is prospective validation in an external population and, ultimately, a trial testing whether geometry-based selection improves efficiency or treatment-effect detection. The present study does not replace such a trial; it propose a tractable way to run it.

## Methods

### Study design and participants

We conducted a development–replication study using two independent components of the US National Health and Nutrition Examination Survey (NHANES), each linked to National Death Index records through 31 December 2019. Development used ten continuous NHANES cycles (1999–2018); replication used NHANES III (1988–1994), a disjoint sample collected on different assay platforms. All source data are public, and participants provided written informed consent under protocols approved by the National Center for Health Statistics Research Ethics Review Board. Analyses were unweighted because the objective was model development and transport across measurement eras, not nationally representative prevalence estimation.

The eligible development population comprised non-pregnant adults aged 18–79 years who were not acutely unwell at examination (n = 34,785); 29,053 had complete data for the 38-marker matrix, of whom 29,023 had linked mortality follow-up. Thirty participants were ineligible for National Death Index linkage; they are excluded from every survival model but retained in analyses that do not use follow-up time. Age was capped at 79 because NHANES top-codes older ages. Analysis-specific complete-case cohorts were used because questionnaire and examination availability differ; no imputation was performed. Descriptive and mortality-gap analyses used the broader eligible cohort rather than the biomarker-complete subset to avoid conditioning population comparisons on panel availability. Direction and panel analyses were restricted to 11,497 participants aged 52 years or older, including 2,494 disease deaths. The boundary was fixed before any mortality-linked analysis and was derived from questionnaire data: among women reporting a natural final menstrual period, with surgical menopause excluded, the median age at menopause was 50 years (interquartile range 45–52), and 52 — the upper quartile, and therefore the upper bound of the perimenopausal band — was taken so that the older stratum would lie beyond the transition for most women. It was subsequently observed, and played no part in setting the boundary, that women’s red-cell indices change course over the same period while men’s do not: the female-minus-male difference in haemoglobin and in red cell distribution width closes fastest during ages 45–55 (+0.061 g/dL and −0.068% per year) and much more slowly on either side, whereas the male trends run in one direction throughout — the pattern expected when menstrual iron loss ceases. Mean corpuscular volume shows no such change. This places the shift in the 45–55 decade, the interquartile range of reported menopausal age, rather than at age 52 specifically, and because it comes from the same cohort it is convergent internal evidence rather than independent validation. Age 52 is therefore an age threshold applied identically to women and men and chosen to sit beyond the menopausal transition at the population level; it is not a claim that any individual woman is post-menopausal at 52, and hormonal status was not measured. Mortality data were not used in choosing it, and the results do not depend on it: age floors of 50, 52 and 55 gave held-out reproduction of 0.920, 0.918 and 0.915, and the sex-by-stratum interaction is non-significant at every boundary from 45 to 55 (Supplementary Information §S1.3, §S4.4).

NHANES records participants as female or male. We use *women* and *men* for those recorded categories and *sex* for the corresponding binary variable, which does not resolve chromosomal, gonadal, hormonal, reproductive or gender-related dimensions that may contribute to the differences observed.

### Mortality, morbidity and behavioural variables

Follow-up began at examination. Disease mortality — any death other than accidental death — was the primary outcome. Accidental deaths were censored at their recorded time because participants remained at risk of disease death until that point. Cause-specific analyses treated the cause of interest as the event and censored other causes at their observed times.

Five-year disease mortality in the trial-sorting analysis was expressed as an incidence rate: disease deaths within five years divided by person-time truncated at five years. This avoids treating the 23.8% of participants with less than five years of linked follow-up as complete denominators. Accidental deaths contributed person-time only until death. Of 2,539 deaths in the relevant cohort, 45 were accidental and 10 occurred within five years, removing 22 of 52,363 person-years.

Morbidity was the count of thirteen questionnaire-defined doctor-diagnosed conditions, calculated only among participants answering all thirteen items. Smoking was classified as never, former or current; physical inactivity used the harmonised activity classification; diet used the Healthy Eating Index 2020 total score ^36,37^, reversed so that higher values indicate poorer diet.

### Biomarker preparation and sex-specific young-healthy reference states

The physiological matrix contained 38 routine haematological, biochemical, urinary, anthropometric and cardiovascular measurements available across all ten development cycles. Units were harmonised and confirmed implausible values set to missing. Extreme but physiologically possible values were retained because participants removed by trimming were substantially enriched for mortality (death-rate ratio 2.42).

For each marker, one transformation was selected from identity, square root, cube root, logarithm or reciprocal using a locked rule based on absolute skewness, excess kurtosis and quantile–quantile deviation. Identity was retained when the raw scale met all three criteria; otherwise the least aggressive adequate transformation was chosen. Selection was unsupervised — age, sex, reference status and mortality were not used — and was performed on the complete 29,053-person panel.

The young-healthy reference comprised participants under 30 years with body mass index 18.5–24.9 kg m⁻², who had never smoked, were not heavy drinkers, and reported no included chronic condition, functional impairment, diabetes or kidney failure, giving 1,238 participants (547 men, 691 women). Formal equivalence testing showed that most markers did not satisfy location- and scale-equivalence between healthy women and men, so each transformed marker was centred and scaled within sex. These coordinates express departure from sex-appropriate young physiology; they do not imply identical raw concentrations, clinical thresholds or biomarker–mortality gradients between sexes.

### Physiological distance

Two representations were used, neither of which reads mortality. *Homeostatic dysregulation* ^18^ is Mahalanobis distance ^38^ from the same-sex young-healthy centroid, using the ordinary unbiased sample covariance of the reference; it is direction-neutral and accounts for correlation between markers, so jointly improbable combinations of individually unremarkable values register as distant.

The *sensitised linear frailty index* expresses each transformed marker relative to its same-sex young-healthy median using a normal-consistent robust scale, the interquartile range divided by 1.349. The adverse excursion is taken according to each marker’s prespecified direction — the positive part for high-adverse markers, the negative part for low-adverse markers, the absolute deviation for bidirectional markers — and deficit severity is that excursion itself, with no lower dead zone and no upper cap. Severity is therefore proportional to displacement at every magnitude and the construction has no free parameters. The index is the unweighted mean across 38 markers. Because deficits are continuous, reference-relative and unbounded above rather than binary and clinically defined, values are not on the conventional zero-to-one frailty scale and are not comparable with published frailty index values.

Adverse directions were assigned as clinical priors before scoring, with a written physiological justification for each marker, and locked by hash. Directionality was not selected or revised using age, sex, mortality, disease or the observed distribution; doing so would make the coordinates outcome-selected before cross-validation.

### Mortality direction and decomposition

Mortality supervision entered only after both representations were defined. Among participants aged 52 years or older, Cox proportional-hazards models were fitted on either coordinate vector with sex included unpenalised and outside the vector. Writing β for the fitted biomarker coefficients, the unit mortality-associated direction is **u** = β/‖β‖₂, and for a participant with displacement **v** we define magnitude *m* = ‖**v**‖₂, alignment *a* = **v**ᵀ**u**/‖**v**‖₂, projection *q* = **v**ᵀ**u** = *m* × *a*, and orthogonal displacement *R* = √(*m*² − *q*²). Orthogonal displacement requires the full 38-dimensional space; computed within a panel selected for alignment with **u**, almost no orthogonal remainder exists by construction.

Projection is proportional to the biomarker component of the fitted Cox linear predictor. It is not an estimate of absolute risk. The fitted direction is predictive rather than causal, and orthogonal displacement is not assumed to be benign.

### Cross-fitting and prevention of outcome leakage

Held-out estimation grouped participants by whole survey cycle; no cycle was split between training and test data. Participant-level directions were fitted leave-one-cycle-out. For panel selection and trial sorting, all 126 five-versus-five cycle splits were evaluated in both directions, yielding 252 directed held-out evaluations. Complete enumeration avoids dependence on a favourable random split; a 20-partition subset produced near-identical medians. Signal-location analyses used 20 seeded partitions with coefficients cross-fitted between halves.

All information that could influence held-out predictions was confined to training data: reference parameters where relevant, direction estimation, tuning, marker ranking and selection, coefficient refitting, score scaling and threshold selection. Because the 252 evaluations overlap and complementary directions are correlated, they were not treated as independent studies. Medians and 2.5th–97.5th percentiles across partitions describe sensitivity to cycle allocation, not confidence intervals. Cross-fitting prevents direct outcome leakage and resubstitution optimism; it does not make a mortality-supervised score causal, outcome-independent or a validated surrogate.

### Reduced-panel development

Panel reduction used nested cycle-grouped validation in participants aged 52 years or older. Within each outer training half, elastic-net penalised Cox models were tuned using grouped inner folds and used to rank all 38 markers ^39^; sex was unpenalised and marker name provided a deterministic tie-break. Elastic net was chosen because correlated biomarkers can make pure L₁ selection unstable, whereas the combined L₁/L₂ penalty can retain groups of correlated predictors. Ranking by coefficient magnitude was appropriate because all coordinates used the same scale: standard deviations of the same-sex young-healthy reference.

Candidate panels of 3, 5, 7, 10, 12 and 15 markers were refitted in the outer training data and applied unchanged to held-out cycles. Eight adequacy criteria were fixed before results were generated, covering reproduction of the 38-marker projection, transfer across encodings, retention of mortality information, within-sex association and preservation of the female–male contrast. After panel size was fixed, the frozen algorithm was applied once to the complete development cohort to produce the locked identity. Nested validation therefore estimates the performance of the selection algorithm at a given size; the exact marker identity remains a development product requiring external validation.

### Trial-arm sorting

A single geometric rule assigned participants to a lifespan arm if projection was at or above the training 90th percentile, and to a physiologically displaced healthspan arm if projection was at or below the 60th percentile and orthogonal displacement at or above the 60th; the rules cannot overlap. Thresholds were estimated within training cycles and applied unchanged to held-out cycles across the same 252 directed partitions, and were carried over from previously qualified rules rather than searched. No weighted score was fitted on the panel markers; the geometry was used as computed. The sort is not mortality-independent, since the direction is fitted against disease mortality, but it is leakage-free: no participant’s assignment uses an outcome from their own cycle. Sex was measured after assignment and never used to select or adjust a threshold, panel or representation.

### External replication

All NHANES III predictions, falsification criteria and design decisions were recorded and signed off before mortality-linked analysis. The young-healthy reference, transformations, standardisation and mortality direction were rebuilt within NHANES III so that agreement reflected replication of the pattern rather than transport of a fitted model. The adverse-direction specification was transferred unchanged as a clinical prior, while transformation orientation was recalculated.

Of the 38 markers, 34 were available: the five-part automated differential was replaced by the three-part Coulter differential, and gamma-glutamyl transferase and globulin were excluded as they were fielded in fewer than half of Phase 1 participants. The replication cohort comprised 13,195 adults aged 18–79 with complete data, of whom 4,474 were aged 52 or older with 3,401 disease deaths; the rebuilt young-healthy reference contained 1,053 participants. Because NHANES III has two survey phases rather than ten cycles, cross-fitting was grouped ten-fold by survey primary sampling unit, preserving the principle that correlated sampling units are never split.

Panel transport imported the locked marker identity unchanged and refitted coefficients within NHANES III; no markers were reselected. Performance was judged against two comparators: 200 random eight-marker panels and a ceiling defined by eight markers selected natively within NHANES III. This distinguishes genuine transport of the mortality direction from the correlation expected simply from retaining eight dimensions.

### System-coverage candidates in the development cohort

Three candidate measurements were nominated in the module specification, on measurement rationale and available statistical power, before any result was examined: high-sensitivity C-reactive protein, because the panel contains no acute-phase reactant and this is the default inflammatory marker of ageing research; fasting glucose and insulin with HOMA-IR, because the panel contains glycated haemoglobin but no fasting glycaemic measure; and spirometric FEV₁, because the panel contains no measure of any organ’s mechanical capacity. Four further candidates were considered and excluded in the same document, with reasons recorded in advance so that the omissions are not silent. Cardiorespiratory fitness was measured only to age 49 and therefore has no participants at all in the age-52-or-older cohort. Grip strength was fielded in two cycles, yielding 263 deaths in this cohort and the shortest follow-up of any candidate, and knee-extensor strength in two cycles permitting reciprocal evaluation only; both are underpowered for a held-out test at the standard applied to the retained three. Apolipoprotein B was fielded in three cycles and correlates at r ≈ 0.93 with non-HDL cholesterol, which the panel already contains. None of the four is claimed to be uninformative; each is untestable in this cohort at a standard comparable to the three retained.

Each retained candidate was evaluated in its own eligible survey cycles, since no common complete-case cohort exists across them, and always on identical participant rows, so that models with and without the candidate were never fitted on different participants. Comparators were the cross-fitted projections of the full 38-marker panel and of the locked ten-marker panel. C-reactive protein and insulin were log-transformed, and C-reactive protein was standardised within assay era because the measurement changed between the 2009–2010 and 2015–2016 cycles, with none fielded in 2011– 2014; an era sensitivity is reported. Spirometry was restricted to quality grades A and B and adjusted for standing height. Evidence of added information was the change in held-out partial log-likelihood under leave-one-cycle-out cross-fitting, with the conditional hazard ratio reported alongside; the unconditional association is a reference quantity and is not evidence of added information. Because the number of eligible cycles differs by candidate — seven for C-reactive protein, nine for the fasting measures, three for FEV₁ — held-out results are not pooled across candidates.

### Functional measurements

Spirometry, bone densitometry, bioimpedance and tandem-stand balance were tested in NHANES III for information beyond the 34-marker projection. These analyses were specified after the primary replication and are exploratory. Models with and without each candidate used identical participant rows, were cross-fitted by sampling unit, and were compared by held-out partial log-likelihood and conditional hazard ratio. Candidates were oriented so that higher values were adverse. For scale, each of the 34 panel markers was removed in turn and the corresponding loss in held-out log-likelihood recorded.

### Statistics and reproducibility

All tests were two-sided with α = 0.05, and exact P values are reported. Survival models used Cox proportional hazards with Efron handling of ties. From the direction analysis onward, attained age with delayed entry at examination age was the primary time scale, allowing participants to be compared at the same age without parametric modelling of the age–hazard relationship. Descriptive mortality models used follow-up time with age adjustment; the alternative time scale is reported as a sensitivity. Baseline hazards were stratified by survey cycle or phase. Proportional hazards were assessed with scaled Schoenfeld residuals ^40^; because projection showed age-varying effects, its per-standard-deviation hazard ratio is interpreted as a cohort-averaged association and the attained-age gradient is reported explicitly.

Adjusted sex contrasts used ordinary least squares with a centred four-degree-of-freedom cubic spline in age and survey-cycle fixed effects, with HC3 heteroskedasticity-consistent standard errors.

The spline basis was centred to keep the design matrix full rank. Continuous outcomes were standardised within the analysis population where per-standard-deviation effects are reported. Marker distributions were examined before and after transformation using skewness, excess kurtosis and D’Agostino K² tests. Held-out predictive performance was compared by partial log-likelihood and discrimination by Harrell’s concordance statistic.

Direction-analysis uncertainty used 500 participant bootstrap replicates stratified by sex and survey cycle, with all leave-one-cycle-out directions refitted within each replicate. Cause-specific analyses controlled the false-discovery rate within that family using the Benjamini–Hochberg procedure ^41^. No single correction was applied across conceptually distinct prespecified analyses. Exploratory analyses are labelled as such and interpreted using conditional association, held-out transport and consistency rather than isolated P values. Cycle-split percentiles are distinguished from confidence intervals throughout.

Analyses used Python 3.10 with NumPy 1.26.4, pandas 2.2.3, SciPy 1.15.3, statsmodels 0.14.5, lifelines 0.30.0, patsy 1.0.2, scikit-learn 1.7.2 and Matplotlib. Analyses are deterministic with fixed recorded seeds. Each analysis module verifies SHA-256 hashes of its locked inputs before execution and refuses to run on changed inputs, and analysis fingerprints combine contract, configuration, input and code hashes so that results generated under different settings cannot be silently combined.

### Use of large language models

Large language models (Claude, Anthropic; ChatGPT, OpenAI) were used for code translation from R to Python, code clean-up and restructuring, analysis code implementation, verification of computed results against primary output files, and manuscript drafting, under author direction, and were additionally used to audit the manuscript against the analysis outputs. Method selection for one supplementary concordance analysis (Supplementary Information §S1.3) was model-proposed and author-approved. All other study design decisions, analysis specifications, predictions and interpretations were made by the authors, and pre-analysis specifications were recorded and signed off before the corresponding analyses were run. No model is an author: the models could not approve the submission or take responsibility for the work, and accountability for its integrity rests with the named authors. Documentation of the analysis pipeline and its outputs is provided in the analysis compendium.

## Supporting information

Revised Supplementary

## Code availability

All analysis code, configuration files, pre-analysis specifications and the executable pipeline are available in the analysis compendium deposited at GitHub: https://github.com/AngusHarding/physiological-direction.

## Data availability

All data analysed in this study are publicly available from the US National Center for Health Statistics.

Continuous NHANES (1999–2018) and NHANES III (1988–1994) survey files are available at https://www.cdc.gov/nchs/nhanes/, and the linked public-use mortality files at https://www.cdc.gov/nchs/data-linkage/mortality-public.htm.

## Acknowledgements

J.F. is supported by the National Natural Science Foundation of China (No. 12371486).

## Author Contributions

### Angus Harding (corresponding Author)

Study conception and design, data acquisition, data cleaning and analysis, data interpretation, initial code generation, creation of figures and tables, manuscript writing, manuscript submission.

### Jim Coward

Study conception and design.

### Jinping Feng

Manuscript writing and review.

### Tianhai Tian

Study conception and design, manuscript writing and review.

## Competing Interests

A.S.H. is director of BH Biotech Pty Ltd, which develops geroscience interventions that could benefit from more cost-effective trial design. An earlier analysis of related NHANES data was submitted as a Masters thesis by A.S.H. (Applied and Computational Mathematics, Monash University, conferred 05-Feb-2025).

J.C., J.F. and T.T. declare no competing interests.

## Abbreviations

AA: Associate of Arts
ACR: urine albumin-to-creatinine ratio
FEV₁: forced expiratory volume in one second
FI: sensitised frailty index
FPL: federal poverty level
GED: General Educational Development certificate
HC3: heteroskedasticity-consistent covariance estimator, type 3
HD: homeostatic dysregulation
HDL: high-density lipoprotein
HOMA-IR: homeostatic model assessment of insulin resistance
L₁ and L₂: the lasso and ridge penalty norms of the elastic net
NHANES: National Health and Nutrition Examination Survey
P1–P7b: the predictions registered before the replication analysis (Table 3)
Q1 and Q3: first and third quartiles
RR: risk ratio
SD: standard deviation
SMD: standardised mean difference. Marker names in Table 2 are listed with their NHANES variable codes.

## Notes

https://github.com/AngusHarding/harding-et-al-2026-biological-age

https://github.com/AngusHarding/physiological-direction

### Summary of Updates

This revision substantially restructures the manuscript following reanalysis after peer-review. The original study focused on mortality-supervised biological-age scores and interpreted sex differences primarily as evidence for sex-specific biomarker requirements. The revised study instead separates outcome-independent physiological displacement from a subsequently estimated, cross-fitted mortality-associated direction. Biomarker transformations are selected using an unsupervised rule independent of age, sex, health status and mortality, reducing outcome-dependent preprocessing and improving the portability of the analytical framework. This change produces a different and more directly translational conclusion. Women and men are not distinguished primarily by the amount of physiological dysregulation, but by their position along a substantially shared mortality-associated physiological axis. The revised study shows that this direction can be reproduced using a small panel of routine measurements and proposes a practical solution for geroscience trial recruitment: within-sex thresholds can maintain sex balance while preserving most of the expected mortality-event enrichment. The central findings were additionally tested in the disjoint NHANES III cohort under predictions and falsification criteria specified before mortality-linked replication analysis; seven of eight predictions were confirmed and one was partially met.

