## Supplementary material for "Direction, Not Distance: A Mortality-Associated Physiological Axis Maps the Sex–Mortality Gap and Informs Sex-Balanced Trial Recruitment": Revised Supplementary

Version 1.0 · 11 August 2026

This document accompanies the manuscript of the same name. It is written to be audited: every section states its hypothesis, its method and the reason for that method, the population it is estimated in, its result, and the code and output files the result comes from. `CODE_INDEX.md` in the analysis compendium maps each section to its scripts and outputs.

---

### About this document

This is the **source of truth** for the analysis. Every method, every number and every figure in the manuscript traces to a section here. It is written to be read by a human reviewer and audited by an AI collaborator, and it is exported to PDF for submission.

Its job is not to record *what* the code did — hashes, manifests and regression tests already prove that. Its job is to record **why each step was chosen and what it is for**. That is the check no test can automate, and its absence is what allowed two conceptual errors into the earlier analysis (§0.4).

#### The three-document rule

| Document | Answers | Authority |
|---|---|---|
| **SUPPLEMENTARY.md** (this file) | Why and how we did it; what we found | **Source of truth** for method and result |
| **MANUSCRIPT.md** | What we claim | Derived; generated from the submitted `.docx` |
| **STATUS.md** | Where the work currently stands | Internal, transient |

**If a finding or a result number appears in STATUS.md, that is a bug.** STATUS records state only — what is built, running, blocked or stale. This prevents three documents drifting into three versions of the truth.

#### How to audit a section

Each section below answers, in order: hypothesis, aim, method with rationale, **estimation-population declaration**, results, conclusion, code, outputs.

To audit, read the **estimation-population table** first and ask of every row: *is the population this parameter was estimated on the right one for the use it is put to?* Both historical errors are visible from that single question.

---

### §0. Global conventions

#### 0.1 Data source

NHANES 1999–2018, ten two-year survey cycles, public-use files linked to the National Death Index through 31 December 2019. All raw files are read but never modified; every derived file is written under `data_preparation/outputs/`.

#### 0.2 Cohorts

| Cohort | n | Defined in | Used by |
|---|---|---|---|
| Eligible adults | 34,785 | §S0 | attrition accounting |
| Complete-covariate (Table 1) | 29,250 | §S1 | Table 1, descriptive comparison |
| Survival | 29,248 | §S1 | all §S1 mortality models |
| 38-biomarker panel | 29,053 | §S0 | §S2 onward |
| Young-healthy reference | 1,238 (547 M, 691 F) | §S0 | reference parameters |
| Direction / selection (age 52–79) | 11,497 (5,920 M, 5,577 F) | §S3 | §S3–§S4 |

Superseded values that may appear in earlier drafts: **27,061 / 27,059** (pre-correction complete-covariate and survival cohorts).

#### 0.3 Recurring conventions

- **Disease death** excludes accidental death, which is censored.
- **Sex** refers to the male and female categories recorded in NHANES.
- **Cycle-split percentiles** describe sensitivity to how survey cycles were allocated between training and test. They are **not confidence intervals**; partitions overlap heavily and complementary directions are not independent.
- **Held-out** means the participants contributed nothing to marker selection, model tuning, or coefficient estimation.

#### 0.4 Analytical provenance and corrections

Two conceptual errors were identified in earlier versions of this analysis. In both cases the code was correct and every validation gate passed; the defect was that the step did not serve its stated purpose. Both are recorded here because they explain why published numbers differ from earlier drafts.

| # | Defect | Consequence | Status |
|---|---|---|---|
| 1 | Biomarker transformations were selected on the young-healthy reference rather than the analysis cohort | Transformations normalised a subgroup, not the distribution they were applied to. 2 of 38 choices changed on correction (eosinophils, HbA1c) | **Corrected.** `data_preparation/TRANSFORMATION_PR`[illegible] selects on all 29,053 |

| # | Defect | Consequence | Status |
|---|---|---|---|
| 2 | The sensitised **linear** frailty index was flat at both ends: a dead zone (`a = 1.0`) zeroed all deviation within one robust-scale unit of the reference, and a cap (`c = 3.0`) flattened everything beyond three | Made a linear index piecewise-thresholded, contradicting its stated design. The dead zone zeroed 58.3% of values in the band nearest the reference where mortality signal is present; the cap flattened 68.7% of waist-to-height values, the largest single contributor to the index | **Corrected 5 Aug 2026.** `FI_METHODS.md` Amendment 1; severity is now the adverse excursion itself, with no free parameters. 484,000 values (43.8% of the matrix) previously flattened now carry information; 18.3% of participants move more than ten percentile points in index rank |

Neither defect is detectable by code review, testing or hash verification. Both are visible from the estimation-population and purpose declarations that each section below is required to state.

---

### §S0. Data preparation

Everything downstream rests on this section. It documents the chain from raw NHANES files to the two scored coordinate systems, in the order the chain runs.

Code: `data_preparation/` · executable run order in `RUN_ORDER.md`

#### Pipeline roadmap

Read this table first. It gives the order of operations, where each stage is documented below, and what it produces. **The subsection numbers are labels, not an ordering** — §S0.1 and §S0.2 were written first and describe stages 9 and 11.

| # | Stage | Documented in | Principal output |
|---|---|---|---|
| 1 | Raw file inventory and manifest | §S0.3 | `provenance/raw_file_manifest.csv` |
| 2 | Mortality linkage, lossless parse | §S0.4 | `01_imported/mortality.csv` |
| 3 | Eligible adult cohort (34,785) | §S0.5 | `04_eligibility/eligible_cohort.cs` |
| 4 | Biomarker import and harmonisation, 8 domains | §S0.6 | `02_harmonised/*.csv` |
| 5 | Locked raw 38-marker panel (29,053) | §S0.6 | `05_panel/final_38_panel.csv` |
| 6 | Self-reported disease reconstruction | §S0.7 | `02_harmonised/self_reported_disea` |
| 7 | Behavioural and socioeconomic variables | §S0.8 | `02_harmonised/{smoking_status,alc` |
| 8 | Transformation selection on all 29,053 | §S0.9 | `06_transform/transformed_panel.cs` |

| # | Stage | Documented in | Principal output |
|---|---|---|---|
| 9 | Young-healthy reference definition (1,238) | **§S0.1** | `08_reference/healthy_reference_me` |
| 10 | Adverse-direction specification, locked and gated | **§S5** | `config/fi_scoring_spec.csv` |
| 11 | Sex-specific standardisation; FI and HD scoring | §S9.3–S9.5 | `11_fi/`, `12_hd/` |
| 12 | Distributional verification | **§S0.2** | `10_validation/fi_distribution_san` |

Three rules hold throughout and are not relaxed anywhere in the chain.

**Raw files are read, never modified.** Every derived file is written beneath `data_preparation/outputs/`. The raw manifest records a SHA-256 for each source file so that any later change is detectable.

**Structural absence is not missingness.** Where NHANES does not field a question for a participant — a skip pattern, a cycle in which an assay was not run — the value remains missing rather than being coded as a negative response. This distinction is the single most common source of silent error in NHANES harmonisation and is enforced by test in every affected domain.

**Reconstruction is audited against the historical build, and disagreement fails visibly.** Where a certified historical output exists, the rebuilt file is compared participant by participant. Code is never adjusted to reach a remembered count.

#### S0.1 Sex-specific young-healthy reference states

**Aim** Every coordinate in this analysis has the form $z = (\,\text{transform}(x) - \text{centre}\,)/\text{scale}$, where the centre and scale come from a young-healthy reference population. Whether that reference is estimated **pooled across sex** or **separately within sex** is therefore a decision that propagates into every downstream quantity — HD, the FI, the mortality direction, the reduced panel and both recruitment screens.

This section tests, per marker, whether men and women share a young-healthy state closely enough to justify one pooled reference.

**Methodological approach and why** **Equivalence testing, not difference testing.** With 547 healthy men and 691 healthy women, a conventional difference test rejects the null for nearly every marker, because sample size makes trivial gaps significant. That answers the wrong question. The decision requires the opposite: can we *affirm* the sexes are close enough to pool? That is a two-one-sided-test (TOST) equivalence procedure against a margin fixed in advance, and the decision is driven by effect size rather than by p.

**Prespecified margin: ±0.20 pooled SD, α = 0.05.** The quantity tested is

$$\delta = \frac{\bar{x}_F - \bar{x}_M}{s_{\text{pooled}}}$$

which has a direct interpretation: **δ is the artificial separation, in z-units, that a pooled centre would inject between the sexes** before any biology is considered. Scale is reported alongside as $s_F/s_M$ with Levene's test.

**Computed on the transformed scale**, because that is the scale the coordinates live on and where pooling would actually occur.

**Outcome-blind by construction.** No mortality or disease variable is read.

**Estimation-population declaration**

| Quantity | Estimated on | Why | Used for |
|---|---|---|---|
| Location gap δ, scale ratio | Young-healthy reference, split by sex (547 M, 691 F) | The question *is* whether these two populations differ | The pooling decision |
| Reference median, robust scale | Same-sex young-healthy reference | Robust set point for the portable FI | FI deficits (§S9.5) |
| Reference mean, SD, covariance | Same-sex young-healthy reference | Moment estimators for Mahalanobis distance | HD (§S9.4) |

**Results** **Thirty-four of thirty-eight markers cannot be affirmed as poolable.** Four met the equivalence criterion; 17 markers showed location gaps exceeding 0.5 pooled SD and 3 exceeded 1.0.

| Marker | Unit | Male median | Female median | δ (pooled SD) | Decision |
|---|---|---|---|---|---|
| Haemoglobin | g/dL | 15.4 | 13.4 | **−2.008** | sex-specific |
| Uric acid | mg/dL | 5.6 | 4.2 | **−1.533** | sex-specific |
| Serum albumin | g/dL | 4.6 | 4.4 | **−1.020** | sex-specific |
| Bicarbonate | mmol/L | 26.0 | 24.0 | −0.930 | sex-specific |
| Alanine amino-transferase | U/L | 20.0 | 15.0 | +0.790 | sex-specific |
| Systolic blood pressure | mmHg | 114 | 106 | −0.781 | sex-specific |
| Alkaline phosphatase | U/L | 67.0 | 53.0 | −0.727 | sex-specific |
| HDL cholesterol | mg/dL | 52.0 | 60.0 | +0.702 | sex-specific |

Haemoglobin alone would carry a **2.01 pooled-SD artificial sex separation** under a pooled reference — a healthy woman would be scored as substantially deficient against a standard that is largely male. Three markers in the locked ten-marker panel — alanine aminotransferase, gamma-glutamyl transferase and serum albumin — are among those requiring sex-specific referencing, so the panel itself depends on this decision.

These values reproduce an independent earlier implementation of the same test to three decimal places (haemoglobin −2.008 against −2.003; uric acid −1.533 against −1.527; serum albumin −1.020 against −1.017), which is external corroboration that the reference is being selected correctly here.

Because the reference defines a single coordinate system, a mixed reference is not adopted; 34 failing markers determine the system-level decision.

**Conclusion** Men and women do not share a young-healthy physiological state on most routine biomarkers, and on several the difference is large enough that a pooled reference would manufacture a sex difference before any biology was measured. **Sex-specific referencing is therefore not a stylistic choice but a requirement**, and every subsequent sex comparison in this paper measures displacement from each sex's own healthy state rather than from a common standard.

This also constrains interpretation in the opposite direction: because each sex is referenced to itself, none of the sex differences reported later can be artefacts of raw between-sex differences in biomarker level. They are differences in *displacement from one's own healthy norm*.

**Code and outputs** **Code:** `data_preparation/src/10_validate/reference_sex_equivalence.py`

**Outputs:** `outputs/10_validation/reference_sex_equivalence.csv` (all 38 markers, with raw-unit reference intervals, δ, scale ratio, TOST, Welch and Levene results, and the FI and HD reference parameters actually used downstream), `reference_sex_equivalence_summary.csv`

**SUPPLEMENTARY TABLE S1 HERE** — `outputs/10_validation/reference_sex_equivalence.csv`
*Young-healthy reference intervals by sex for all 38 biomarkers, with the equivalence test and the reference parameters used to generate the sex-specific FI and HD coordinates.*

#### S0.2 Distributional sanity check after transformation and normalisation

**Aim** Verify by inspection that the transformation and sex-specific normalisation did what they claim: that each marker's distribution is approximately symmetric on the working scale, that the healthy reference sits where it should, and that the direction of each marker's adverse axis is correct.

This is not a formality. Both conceptual defects in this project's history (§0.4) were invisible to hash verification and unit tests, because in both cases the code correctly implemented what the specification said. They were visible only to a reader asking whether the step served its purpose. This figure is the visual counterpart of the estimation-population declaration: a wrong transformation, a mis-signed orientation, or a reference estimated on the wrong population would all be obvious on sight here.

**Methodological approach and why** **What is plotted.** The raw-oriented robust coordinates from FI Step 4, which is where transformation and normalisation end and the directionality and deficit logic begin. This is the last point at which the data are still a plain distribution rather than a constructed index.

**Why a shared x-axis is the point.** Because every marker is divided by its own same-sex young-healthy robust scale, all 38 markers occupy one common axis in interpretable units — young-healthy scale units from that sex's healthy median. A grid of separately scaled panels would hide the single most useful comparison: which markers the ageing population has drifted furthest on. The markers are ordered by that drift, so the ones dominating the index appear first.

**Why densities rather than histograms.** Bin-edge placement can create or hide apparent modes. Kernel density estimation with a common grid is bin-free and directly comparable between the sexes. Densities are computed on a seeded subsample of at most 5,000 per group for tractability; the estimate is visually indistinguishable from the full-sample version and is deterministic.

**Orientation convention.** Positive always means "above the healthy median on the *original raw-value* axis", because the orientation multiplier restores the raw direction after any reciprocal transformation. Without this, reciprocal- transformed markers would appear reversed and the figure would mislead.

**Outcome-blind.** No mortality or disease variable is read.

**Estimation-population declaration**

| Quantity | Estimated on | Why | Used for |
|---|---|---|---|
| Reference median, robust scale | Same-sex young-healthy reference | Defines the zero and unit of the axis | Coordinate construction |
| Reference 2.5th–97.5th interval | Same-sex young-healthy reference | The healthy range the cohort is compared against | Shaded band |
| Skew, kurtosis, D'Agostino $K^2$ | Full cohort, by sex | Distribution shape is a property of the data the analysis runs on, not of the reference | Diagnostic table |
| Density curves | Full cohort, by sex | Shows where the analysed population actually sits | Figure |

**Results** **The construction behaves as specified.** Estimated glomerular filtration rate and serum albumin sit left of the healthy median, as they must — both decline with age and the orientation convention maps decline to negative. All other markers sit predominantly right of zero. No marker points the wrong way.

**The young-healthy reference is of normal width.** The median reference 2.5th–97.5th interval spans **4.08 scale units**, against the 3.92 expected for a normal distribution given that the robust scale equals the standard deviation under normality. The reference is therefore neither unusually tight nor unusually broad on the working scale: the normalisation is doing what it claims.

**Population drift is visible, unequal, and concentrated.** Across all markers a median of **4.6%** of the cohort exceeds its sex's young-healthy 97.5th percentile — close to the 2.5% that would obtain if the ageing cohort resembled the reference. Waist-to-height ratio is the extreme departure: **68.9% of men and 63.0% of women** exceed the young-healthy 97.5th percentile, and the distribution is centred near +3 scale units with a tail beyond +10. This is the direct visual form of the capping problem that motivated Amendment 1 (§0.4). Glycated haemoglobin (30.8% of women, 21.6% of men) and systolic blood pressure (29.1% of women) follow. Across all 38 markers only 0.14% of values fall outside the plotted range.

**Sex differences appear where physiology predicts.** Alkaline phosphatase and lactate dehydrogenase are displaced further in women; potassium and chloride further in men; serum albumin further in men.

**The raw-scale companion figures resolve an ambiguity the normalised figure cannot.** A wide reference band on the normalised scale could mean either a genuinely heterogeneous reference population or a small robust scale. Plotting raw values with the reference overlaid shows the young-healthy reference is only modestly tighter than the full adult cohort: the **median reference-to-cohort width ratio is 0.827**, and for a few markers — total bilirubin (1.29 in men), eosinophil count, alkaline phosphatase, red cell distribution width — the reference is actually *wider*. Being young and healthy does not imply physiological homogeneity, which is directly relevant to the manuscript's claim that clinically normal physiology is not physiologically uniform.

**Limitation 1 — residual skew.** Transformation reduced but did not eliminate asymmetry. Eight of 76 marker-by-sex combinations exceed |skew| = 1 and twelve exceed |excess kurtosis| = 3. The worst are the urine albumin-creatinine ratio (skew 2.00 in men, 1.82 in women) and glycated haemoglobin (1.64 in women, 1.47 in men), both visible as unusually wide reference bands. Distances and projections involving these markers carry more tail influence than a symmetric marker would contribute.

**Limitation 2 — measurement granularity (previously unrecorded).** Several markers are reported at a resolution that is coarse relative to the young-healthy spread they are divided by, so their "continuous" coordinates are effectively ordinal. Bicarbonate is the extreme case: **24 distinct values across 29,053 participants, with one measurement step spanning 0.67 young-healthy scale units.** Ten of 38 markers have a step exceeding 0.10 scale units — after bicarbonate, total calcium (0.39), serum albumin (0.26), globulin (0.23), pulse rate (0.21), serum phosphorus (0.18) and total bilirubin (0.18). This appears in the figure as visibly stepped densities for those markers.

This interacts with the superseded version-1 deficit rule: under a one-unit dead zone, bicarbonate could occupy only a very small number of distinct deficit levels. Removing the dead zone does not remove the granularity, but it stops the coarse low end being discarded entirely.

**Conclusion** The transformation and sex-specific normalisation behave as specified, and no marker is misoriented. Two limitations are recorded that no automated check would have raised: residual skew in a small number of markers, and measurement granularity coarse enough in ten markers that their coordinates should be understood as finely ordinal rather than strictly continuous. Neither invalidates the construction; both constrain how finely differences in those markers should be interpreted.

**Code and outputs** **Code:** `fi_distribution_sanity_check.py` (normalised scale) and `raw_distribution_sanity_check.py` (raw scale)

**Outputs:** `outputs/10_validation/fi_distribution_sanity_check.csv` (76 rows: skew, excess kurtosis, D'Agostino $K^2$ and p, young-healthy reference intervals in raw units, and cohort drift on the normalised scale, for each marker and sex), raw_distribution_sanity_check.csv (reference and cohort intervals with their width ratio), and three figures under outputs/10_validation/figures/.

**SUPPLEMENTARY FIGURE S1 HERE** — Supplementary_Figure_FI_distributions.png *Distribution of all 38 biomarkers by sex after transformation and sex-specific normalisation, on a shared axis of young-healthy scale units. Dashed line, same-sex young-healthy median; shaded band, young-healthy 2.5th–97.5th percentile interval. Markers ordered by population drift.*

**SUPPLEMENTARY FIGURE S2 HERE** — Supplementary_Figure_raw_distributions_men.png and Supplementary_Figure_raw_distributions_women.png *Raw biomarker distributions by sex (green) with the same-sex young-healthy reference overlaid (purple), one panel per marker. Purple bar, reference 2.5th–97.5th percentile in raw units. Log axis where the locked transformation is a log, so the display scale matches the scale on which that marker is approximately symmetric.*

**SUPPLEMENTARY TABLE S2 HERE** — outputs/10_validation/fi_distribution_sanity_check.csv *Distribution shape and young-healthy reference intervals by marker and sex.*

#### S0.3 Source data, inventory and provenance

**Aim** Establish an auditable boundary between data we received and data we created, so that any reviewer can confirm no raw file was altered and can identify exactly which source file every derived value came from.

**Method and why** NHANES 1999–2018 comprises ten two-year cycles distributed as separate SAS transport files per cycle and per assay. A study drawing 38 markers from eight measurement domains therefore reads on the order of a hundred distinct files, and the risk is not that one is misread but that a substitution passes unnoticed.

Before any import, build_raw_manifest.py walks the raw-data root and records each file's path, size and SHA-256. Domain-specific manifests are built for each measurement chunk before that chunk is imported. Import scripts read only files present in a manifest.

*Why a manifest rather than trusting the directory:* NHANES filenames repeat across cycles with only a suffix distinguishing them (L40_B, L40_C), and a mis-globbed pattern silently substitutes one cycle for another. A hash manifest converts that failure from invisible to fatal.

**Raw files are opened read-only.** All generated files are written beneath data_preparation/outputs/, and provenance/output_manifest.csv carries a SHA-256 for each so that downstream modules can lock their inputs (§S9.12).

**Code and outputs** **Code:** src/00_inventory/ — build_raw_manifest.py plus one manifest builder per measurement domain

**Outputs:** provenance/raw_file_manifest.csv, provenance/output_manifest.csv

#### S0.4 Mortality linkage and endpoint definition

**Aim** Attach follow-up and cause of death to each participant without discarding information that later analyses might need, and define the study endpoint once.

**Method and why** Source files are the ten 2019 Public-Use Linked Mortality Files, with follow-up through **31 December 2019**. The fixed-width records are parsed to retain linkage eligibility (ELIGSTAT), vital status, the public underlying-cause recode (UCOD_LEADING), multiple-cause-of-death flags, and **both** follow-up clocks — time from interview and time from examination.

**The parse is deliberately lossless.** Accidental deaths (UCOD_LEADING == 4) are retained rather than dropped, missing causes are recoded honestly as missing rather than as a residual category, and no endpoint exclusion happens at import.

*Why losslessly:* an earlier build applied the endpoint definition during import, which made accidental deaths unrecoverable and forced a re-import whenever an analysis needed a different censoring rule. Endpoint construction is a modelling decision and belongs with the model, not with the data. Every survival analysis in this paper therefore applies its own exclusion to a complete file, and the exclusions are visible in the analysis code rather than baked into the input.

**Endpoint definition, applied downstream and identically throughout.** The primary outcome is **disease death**: death from any cause other than accident. Accidental deaths are **censored at their recorded time of death**, not excluded from the risk set, because a participant who died accidentally was at risk of disease death until that moment. Cause-specific analyses treat death from the cause of interest as the event and censor other causes at their observed times.

**Time scale** is examination follow-up in §S1 and attained age with delayed entry from §S3 onward; both are stated per analysis and the choice is reported as a sensitivity wherever it could matter (§S9.6).

**Results** Linkage was left-joined onto the fixed 29,053-person analysis master. Within that subset there are **3,028 deaths, of which 106 are accidental.** Both follow-up clocks are preserved on every record.

**Code and outputs** **Code:** `src/01_import/import_mortality.py`, `src/10_validate/compare_mortality.py` · specification in `MORTALITY_CLEANING.md`

**Outputs:** `outputs/01_imported/mortality.csv`

#### S0.5 The eligible adult cohort

**Aim** Define the broadest defensible adult population before any biomarker availability is considered, so that attrition to the analysed samples is measurable rather than implicit.

**Method and why** Eligibility is assessed on demographic and screening grounds only: adults aged **18–79** years, not pregnant, and not classified as acutely unwell at examination. This yields **34,785 participants**.

*Why eligibility precedes biomarker completeness:* requiring all 38 markers selects a healthier, better-attended subgroup. Establishing the eligible population first makes that selection visible as an attrition step with a recorded count, rather than absorbing it into the definition of the cohort. §S1 exploits this directly by running the descriptive sex–mortality contrast on the broad cohort rather than the panel-complete one, so that a population contrast is not confounded with panel availability.

Age is capped at 79 because NHANES top-codes age above that threshold, so ages beyond it are not resolvable.

**Code and outputs** **Code:** `src/04_eligibility/build_eligible_cohort.py`, `src/01_import/import_demographics.py`, `src/01_import/import_recent_illness.py`

**Outputs:** `outputs/04_cohorts/eligible_adult_cohort.csv`, `outputs/audit/cohort_attrition.csv`

#### S0.6 Biomarker panel assembly and harmonisation

**Aim** Assemble 38 routinely available measurements, on consistent units and definitions, for every participant in whom all 38 are present.

**Method and why** Markers were imported and harmonised in eight domain chunks, each with its own manifest, harmonisation script and comparison audit. The domains and their contribution to the final panel are:

| Domain | n | Markers |
|---|---|---|
| Liver | 7 | albumin, ALT, AST, total bilirubin, LDH, GGT, alkaline phosphatase |
| Immune | 6 | lymphocytes, neutrophils, monocytes, eosinophils, basophils, globulin |
| Red blood cell | 5 | haemoglobin, MCV, MCHC, RDW, RBC folate |
| Electrolytes | 4 | sodium, potassium, chloride, bicarbonate |
| Kidney | 4 | urea nitrogen, uric acid, albumin-to-creatinine ratio, eGFR |
| Cardiovascular | 3 | systolic pressure, diastolic pressure, pulse rate |
| Lipids | 3 | total cholesterol, HDL cholesterol, triglycerides |
| Calcium–phosphate | 2 | total calcium, phosphorus |
| Platelet | 2 | platelet count, mean platelet volume |
| Metabolism | 2 | glycated haemoglobin, waist-to-height ratio |

**Units are harmonised across cycles and confirmed implausible values set to missing.** Several conventions are recorded because they are decisions rather than mechanics:

- **Blood pressure.** All four readings are retained in the intermediate file; the harmonised value is the mean of readings **2–4**. *Why:* the first reading is systematically elevated by the measurement itself, and discarding it is standard practice — but retaining it in the intermediate preserves the ability to test that choice.
- **Complete blood count.** Measured and mathematically derived quantities are kept in **separate layers**. *Why:* several CBC indices are algebraic functions of others, and mixing them invites a panel containing its own arithmetic.
- **eGFR** is derived and therefore depends on Chunks 1 and 2, requiring age, sex and calibrated serum creatinine. It is a derived feature, flagged as such.
- **Triglycerides.** A dedicated fasting-subsample reconstruction was built for audit, but the panel uses the full-sample `LBXSTR`. *Why:* restricting to the fasting subsample would have cost roughly half the cohort for one marker.

**Panel membership is complete-case across all 38 markers**, giving **29,053** participants of the 34,785 eligible. No imputation is used anywhere in the principal analyses.

*Why complete-case rather than imputation:* the downstream geometry requires a covariance matrix estimated on the young-healthy reference and a per-participant 38-vector. Imputed coordinates would enter both the covariance structure and the distance, and a distance partly composed of model predictions is not the quantity the paper claims to measure.

**Code and outputs** **Code:** `src/01_import/`, `src/02_harmonise/`, `src/03_derive/`, `src/05_panel/build_final_38_panel.p` · one `compare_{mortality,smoking,alcohol,hei}.py` audit per domain in `src/10_validate/`

**Outputs:** `outputs/05_panel/final_38_biomarker_panel_raw.csv`, `config/final_38_panel_spec.csv` (marker, subsystem, locked transformation)

#### S0.7 Self-reported disease reconstruction

**Aim** Rebuild the questionnaire-derived chronic-condition flags used for the morbidity contrast in §S1, and verify the reconstruction against the certified historical build.

**Method and why** Thirteen doctor-diagnosed conditions form the historical morbidity count, and a broader candidate set is reconstructed alongside it for the expanded audit.

**Structural non-coverage remains missing.** Where a condition was not fielded in a cycle, or a participant was skipped past the item, the flag is missing rather than zero. *Why this matters more here than elsewhere:* a morbidity **count** sums across conditions, so coding unasked items as absent produces a count that is biased downward by exactly the number of unfielded items — and unevenly across cycles. The count in §S1 is therefore computed only among participants with complete responses to all thirteen.

**Results All fourteen reconstructed fields agree with the certified historical outputs with zero participant-level mismatches.**

**Code and outputs** **Code:** `src/01_import/import_self_reported_disease.py`, `src/02_harmonise/harmonise_self_repo[illegible]` `src/10_validate/{compare_self_reported_disease,audit_disease_candidates}.py` · audit report in `DISEASE_RECONSTRUCTION_AND_AUDIT.md`

**Outputs:** `outputs/02_harmonised/self_reported_disease_flags.csv`

#### S0.8 Behavioural and socioeconomic variables

**Aim** Derive the covariates used in the §S1 adjustment cascade and the §S7 exposure analyses, on definitions consistent across ten cycles of changing instruments.

**Method and why** **Smoking** is classified never / former / current from `SMQ020` and `SMQ040`. Structurally skipped `SMQ040` values among never-smokers are **not** treated as missing; refused, unknown, inconsistent and genuinely missing responses remain unclassified rather than being assigned a level.

**Alcohol** required reconciling two different instruments: a count-and-unit question fielded 1999–2016 and a categorical-frequency question from 2017–2018. Both are converted to average drinks per week, and the **uncapped estimate is preserved alongside the harmonised one** so that the effect of capping is measurable rather than assumed.

**Physical activity** is published in **two deliberate versions**, and this is the one place where the rebuild does not simply reproduce the historical rule. `physical_activity.csv` reproduces it exactly, including whole-record exclusion of respondents reporting above 10,000 MET-minutes per week. `physical_activity_inclusion_preserving.csv` retains those respondents' binary activity classification and caps only the continuous dose.

*Why both exist:* excluding an entire record because one continuous field is implausible discards that participant's usable binary classification, and those participants are not a random subset. The inclusion-preserving version is the corrected candidate and is the one used downstream; the historical version is retained so the difference is auditable. **This correction moved the analytic cohort** — see §0.2 and the superseded counts recorded there.

**Diet** is the HEI-2020 total score, reproduced from frozen CRAN archives under `provenance/hei_packages/` so the calculation is not hostage to a package update. It covers **2005–2018 only**, because dietary recall begins in that cycle, and is **never a completeness condition for the headline mortality cohort** — requiring it would have cut the cohort by a third for a variable used in one Discussion-tier analysis.

**Socioeconomic variables** — race/ethnicity, education, family income-to-poverty ratio — were imported for the §S7 adjustment models into `outputs/01_imported/socioeconomic.csv`, a new file, so that no existing harmonised output or hash was disturbed.

**Code and outputs** **Code:** `src/01_import/import_{smoking,alcohol,physical_activity,hei_package_data}.py`, `src/02_harmonise/harmonise_{smoking,alcohol,physical_activity}.py`, `src/02_harmonise/calculate_hei_2020.py` · audits in `src/10_validate/compare_{mortality,smoking,alcohol,physical_activity,hei,eligible_cohort,final_3` · tests in `tests/test_{smoking,alcohol,physical_activity,hei}.py`

**Outputs:** `outputs/02_harmonised/{smoking_status,alcohol_drinks_per_week,physical_activity,hei_2020_total}.` `outputs/01_imported/socioeconomic.csv`

#### S0.9 Transformation selection

**Aim** Place each marker on a scale where a unit of deviation means approximately the same thing across the range, which is the precondition for both the unweighted mean in the FI (§S9.5) and the covariance estimate in HD (§S9.4).

**Method and why** One transformation per marker is selected from a fixed candidate set by minimising absolute skewness. Selection is **unsupervised**: age, sex, reference status and mortality are not selection variables.

**Selection is performed on the complete 29,053-person panel.** This is the correction of defect 1 (§0.4). The superseded version selected transformations on the young-healthy reference subgroup, which normalised that subgroup rather than the distribution the transformation was actually applied to. Two of 38 choices changed on correction — eosinophils and glycated haemoglobin.

*Why the analysis cohort and not the reference:* a transformation is a property of the distribution it is applied to. Choosing it on a subgroup optimises symmetry where the analysis does not happen and can leave the analysed distribution skewed. The estimation-population declaration (§S9.11) is what makes this visible; no test or hash would have caught it, because the code correctly implemented a specification that was itself wrong.

The stage deliberately **stops before standardisation**, which requires the reference definition (§S0.1) and is documented at §S9.3.

**Limitations** Transformation reduced but did not eliminate asymmetry; residual skew and measurement granularity are quantified in §S0.2.

**Code and outputs** **Code:** `src/06_transform/` — `assess_transformation_need.py`, `select_full_cohort_transformatio` `assess_hd_readiness.py`, `assess_trimming_impact.py` · protocol in `TRANSFORMATION_PROTOCOL.md`, results in `TRANSFORMATION_RESULTS.md`, trimming decision in `TRIMMING_ASSESSMENT.md`

**Outputs:** `outputs/06_transform/final_38_panel_transformed_unstandardised.csv`, `config/final_38_panel_spec.cs` (locked transformation per marker)

> **On trimming.** `TRIMMING_ASSESSMENT.md` records the decision **not** to remove extreme values, because excluded participants were substantially enriched for mortality (death-rate ratio 2.42): deletion would have removed informative pathological observations rather than noise. That decision is load-bearing for §0.4 defect 2 — the superseded FI cap flattened those same mortality-enriched values one step later, which is inconsistent with retaining them here.

---

### §S1. The sex–mortality gap

> Manuscript section: *"Women had lower disease mortality despite greater reported morbidity"*

#### S1.1 Hypothesis and aim

No formal hypothesis is tested here. This section establishes the **biological contrast that motivates the study**: women report more diagnosed disease than men yet die of disease substantially less often. The aim is to demonstrate that this gap is (a) real in a large, contemporary, nationally representative sample, (b) not an artefact of age structure, survey design, socioeconomic position or health behaviour, and (c) not confined to a single cause of death.

Establishing a gap that survives conventional explanation is what creates the need for a physiological explanation in §S2 onward.

#### S1.2 Methodological approach and why

**Cohort construction.** The analysis begins from the **broad eligible adult cohort**, not the biomarker-complete cohort. This is deliberate: restricting to participants with all 38 biomarkers would select a healthier, better-attended subgroup and would confound a descriptive population contrast with panel availability. Participants are required to have complete demographic, mortality, smoking, alcohol and physical-activity data — the covariates used in the adjustment cascade — and nothing more.

**Survival models.** Cox proportional-hazards regression with Efron ties. Accidental deaths are censored because the study concerns disease biology, not external causes. The primary time scale is follow-up from examination with age adjustment; attained age with delayed entry is reported as a sensitivity, since the choice of time scale is a standard methodological challenge.

**Adjustment cascade rather than one adjusted model.** Covariates are added in interpretable blocks — demographic, then socioeconomic, then behavioural, then morbidity — and the female hazard ratio is reported at each step. A single fully adjusted model would hide *which* explanation fails to account for the gap; the cascade shows the estimate is stable against each in turn.

**Survey weighting.** Reported as a comparator, not as the primary. Unweighted estimates are the primary because the target is a biological association within the sampled population rather than a population-level prevalence estimate; the weighted estimate is reported to demonstrate that the design does not drive it.

**Menopause stratification.** The sex-hormone explanation is the most common alternative account of the female survival advantage, so it is tested directly by splitting at a data-derived menopause boundary and testing the sex × menopause interaction. See §S1.3 for the boundary's provenance.

#### S1.3 Estimation-population declaration

*Read this table first when auditing. Ask of every row: is this the right population for this use?*

| Quantity | Estimated on | Why that population | Used for |
|---|---|---|---|
| Cox coefficients | Survival cohort, n = 29,248 | The population the estimate describes | All §S1 hazard ratios |
| Comorbidity count | Complete-covariate cohort with complete responses to all 13 conditions | Condition-specific denominators differ; a complete-case count avoids treating unasked items as absent | Table 1 morbidity row |
| Menopause boundary (age 52) | RHQ natural-menopause respondents, hysterectomy excluded, ages 30–60 | The boundary must describe natural menopause; including surgical menopause shifts the median down by 3–4 years | §S1 menopause split |
| Survey weights | NHANES design variables as published | Not estimated here | Weighted comparator |

**Menopause boundary provenance.** Median age at natural menopause was 50 (IQR 45–52); the split uses the upper perimenopausal bound (Q3 = 52) so that the post-menopausal stratum coincides with the age ≥ 52 frame used from §S3 onward. The boundary is derived from questionnaire data **first**; it was subsequently found to coincide with a change in red-cell marker behaviour, which is corroboration and must be reported in that order. The cutpoint is not load-bearing: the sex × menopause interaction is non-significant at every cut from 45 to 55.

**The red-cell concordance, quantified.** The claim that the boundary "coincides with a change in red-cell behaviour" is now backed by an artefact rather than asserted (`04b_redcell_boundary_concordance.py`). The estimated quantity is the rate at which the female-minus-male difference closes, within three age windows fixed in advance — reproductive 35–45, transition 45–55 (the questionnaire's own interquartile range widened to a decade), post-transition 55–70 — as the female × age interaction in a model with survey-cycle fixed effects and HC3 standard errors.

| Marker | 35–45 | 45–55 | 55–70 | Windows differ |
|---|---|---|---|---|
| Haemoglobin, g/dL per year | +0.008 (−0.012 to +0.028) | **+0.061 (+0.040 to +0.082)** | +0.019 (+0.006 to +0.032) | $P = 5.1 \times 10^{-4}$ |
| Red cell distribution width, % per year | +0.016 (−0.005 to +0.037) | **−0.068 (−0.090 to −0.045)** | −0.008 (−0.018 to +0.003) | $P = 1.6 \times 10^{-7}$ |
| Mean corpuscular volume, fL per year | +0.045 (−0.053 to +0.144) | +0.019 (−0.078 to +0.116) | −0.029 (−0.085 to +0.027) | $P = 0.37$ |

The movement is confined to the transition window in both haemoglobin and red cell distribution width, and the **direction is sex-specific**: within that window women reverse (haemoglobin +0.055, RDW −0.044 per year) while men continue as before (haemoglobin −0.006, RDW +0.024), and outside it the male trends run in one direction throughout. Mean corpuscular volume shows nothing and is reported as the null it is rather than dropped.

**Two honesty constraints on how far this goes.** First, it locates the change in the **45–55 decade, not at age 52**. A single-knot broken-stick regression was fitted first and is deliberately **not reported**: bootstrap intervals for the knot spanned the whole search grid and several estimates sat on a grid boundary, because both moving markers are three-segment shapes that one knot cannot represent. These data corroborate the perimenopausal band; they do not select 52 within it, and the questionnaire is what selects 52. Second, the comparison uses the same cohort and age is age at examination, so it is convergent internal evidence rather than independent validation, and cohort effects, period effects and differential survival are not separable from ageing.

#### S1.4 Results

**Morbidity is higher in women; mortality is lower.** The cohort comprised 29,250 participants (14,190 women, 15,060 men), closely matched on age, race/ethnicity and income but differing in behaviour — men smoked more and drank more, women were more often sedentary. Among participants with complete responses to all 13 prespecified conditions, mean comorbidity count was **0.87 in women versus 0.73 in men**, and **48.2% of women versus 39.2% of men** reported at least one condition. Disease death nevertheless occurred in **9.5% of women versus 13.4% of men** (male-to-female risk ratio 1.40, 95% CI 1.32–1.50).

**The gap is not explained by age.** Across the adult age range the estimated probability of disease death was consistently lower in women, the absolute difference widening with age (Figure 1A). Among 29,248 participants with positive follow-up, 3,369 disease deaths occurred. Age-adjusted female HR **0.644 (95% CI 0.601–0.690)**; survey-weighted **0.66 (0.60–0.71)**.

**The gap spans several major causes** (Figure 1B): heart disease 0.535 (0.465–0.616), cancer 0.641 (0.560–0.733), cerebrovascular disease 0.607 (0.451–0.818), diabetes as any mention 0.66 (0.54–0.80), hypertension as any mention 0.78 (0.66–0.92). Chronic lower respiratory disease 0.876 (0.663–1.158), influenza/pneumonia and Alzheimer disease were compatible with no sex difference.

**The gap survives the adjustment cascade** (Figure 1C): age-adjusted 0.64 → 0.60 with race/ethnicity, education and income → 0.63 with smoking, alcohol and physical activity → 0.61 with the 13-condition morbidity count. All-cause mortality 0.64 (0.60–0.69); attained-age time scale 0.64 (0.60–0.69).

**The gap does not close after menopause.** Pre-menopausal (age < 52) HR 0.673 (0.559–0.811); post-menopausal (age ≥ 52) 0.639 (0.593–0.688); interaction non-significant. The advantage persists across the menopausal transition and is robust to the cutpoint.

#### S1.5 Conclusion

Women carry more diagnosed morbidity yet die of disease substantially less often, and the gap is not attributable to age structure, survey design, socioeconomic position, health behaviour, reported morbidity, cause-of-death distribution, or the menopausal transition. A conventional epidemiological account does not close it. This motivates the physiological analysis that follows: if the difference is not in *how much* disease burden each sex carries, it may be in *what kind*.

#### S1.6 Code and outputs

**Code:** `analysis/03_sex_mortality_frailty_gap/code/` · cohort build `01_build_analysis_cohort.py` · cause-specific `03_cause_specific.py` · menopause `04a_menopause_hr.py` · red-cell concordance `04b_redcell_boundary_concordance.py` · Table 1 `11_build_table_1.py` · combined figure `07_render_combined_figure_1.py`

**Source data:** `outputs/figures/figure1a_source_data.csv`, `figure1c_source_data.csv`, `cause_specific_female_hr.cs`[illegible] `menopause_hr_table.csv`, `outputs/data/redcell_age_profiles.csv`, `outputs/data/redcell_boundary_breakpoints.c`[illegible] `outputs/audit/cohort_attrition.csv`

**Tests:** `tests/test_table_1.py`, `tests/test_cohort_outputs.py`

> **FIGURE 1 HERE** — `outputs/figures/Figure_1_ABC_combined.png` *(A) modelled probability of disease death by age and sex; (B) cause-specific female hazard ratios; (C) adjustment-cascade robustness.*
>
> **TABLE 1 HERE** — `outputs/tables/Table_1_manuscript_display.csv`
>
> **SUPPLEMENTARY FIGURE 1 HERE** — `outputs/figures/Supplementary_Figure_1_mortality_robustness.png`
>
> **SUPPLEMENTARY FIGURE 2 HERE** — `outputs/figures/Supplementary_Figure_redcell_boundary.png` *Red-cell indices by age and sex, with the questionnaire-derived boundary and perimenopausal band marked; legend in `redcell_boundary_legend.txt`.*
>
> **SUPPLEMENTARY TABLE 1 HERE** — `outputs/tables/Supplementary_Table_1_human_readable.csv`

---

### §S1b. Prognostic movement within the young-healthy reference ranges

> Supports the manuscript Discussion section *“Clinically normal physiology was not physiologically homogeneous”*, and is the empirical justification for the sensitised-linear construction (§0.4 defect 2).

#### S1b.1 Hypothesis and aim

**Claim tested:** a substantial fraction of the prognostic signal in routine biomarkers lies *within* the sex-specific young-healthy reference range — that is, mortality-associated physiology is detectable before any marker becomes clinically abnormal.

This is an obligation rather than an option. The sensitised FI is linear precisely because deviations near the reference are held to carry information; if they do not, the superseded dead zone was defensible and the correction in §0.4 was unnecessary. The section therefore tests the premise the instrument rests on.

#### S1b.2 Methodological approach and why

**Signal is located, not merely quantified.** For each held-out participant and marker, the mortality-associated contribution is the product of that participant's coordinate and the corresponding training-derived mortality coefficient. Positive contributions — those pointing toward greater fitted mortality — are then classified by the participant's **raw** biomarker value into three mutually exclusive zones:

1. within the sex-specific empirical young-healthy 2.5th–97.5th percentile interval;
2. outside that interval but within conventional clinical limits;
3. clinically abnormal.

Where empirical and clinical intervals overlap imperfectly, clinically abnormal classification takes precedence, so the within-limits figure is conservative.

**Cross-fitted, not in-sample.** Twenty seeded five-versus-five cycle partitions are drawn. In each, coefficients are fitted separately in both halves and then **swapped**: every participant is scored with coefficients estimated on cycles they did not contribute to. Contributions are aggregated within partition and summarised as medians with 2.5th–97.5th split percentiles.

*Why cross-fitting is essential here:* an in-sample fit assigns each participant a coefficient partly determined by their own outcome, which inflates the apparent contribution of exactly the near-reference values the claim is about. The superseded preliminary calculation had this defect and could not be reported.

*Why contribution mass rather than a hazard ratio:* the question is where prognostic information is *located across the population*, not how steeply risk rises within a stratum. A zone decomposition answers the former directly.

#### S1b.3 Estimation-population declaration

| Quantity | Estimated on | Why | Used for |
|---|---|---|---|
| Mortality coefficients | Training half of each cycle partition | The scored participant must not have contributed | Contribution products |
| Young-healthy zone boundaries | Same-sex young-healthy reference | The comparison the claim is about | Raw-value classification |
| Clinical limits | External reference intervals and guideline targets; not estimated | Conventional interpretation is the comparator | Zone classification |
| Contribution shares | Held-out participants, age 52–79 | The population the estimate describes | Reported percentages |

#### S1b.4 Results

**The majority of positive mortality-associated signal arises from values inside the young-healthy interval itself.** Medians across 20 partitions:

| Zone | HD | FI |
|---|---:|---:|
| Inside same-sex young-healthy 2.5th–97.5th interval | **57.4%** | **57.1%** |
| Outside that interval, within clinical limits | 10.8% | 8.8% |
| **Within clinical limits (sum)** | **68.1%** | **66.0%** |
| Clinically abnormal | 31.9% | 34.0% |

Roughly two thirds of positive mortality-associated signal therefore lies within conventional clinical limits, and the larger part of that is not merely clinically normal but *young*-normal. Prognostic information is not concentrated just outside abnormality thresholds, where clinical attention would fall; it is distributed throughout the normal region including its centre.

**The two encodings now agree**, differing by 2.1 percentage points on the within-limits total. An earlier version, using the superseded thresholded FI, put the FI figure at 53.0% and made the result look like an artefact of one scoring choice. Removing the dead zone removed the discrepancy — the signal inside the normal range is a property of the measurements, not of how they were scored. This is the most direct evidence that §0.4 defect 2 was a defect and not a preference.

**Sensitivity to the clinical-limit source.** A four-variant sensitivity on the lactate dehydrogenase range (as published, two alternative intervals, and limits removed) moved the headline by at most 0.6 percentage points and always upward, so the correction recorded in §0.4 was conservative.

#### S1b.5 Conclusion

Mortality-associated physiological change is measurable well before conventional abnormality, and most of it occurs while individual values remain inside the range observed in young healthy adults of the same sex. A threshold-based deficit instrument discards this by construction, which is why the index used here is linear with no dead zone.

**What this does not establish.** Near-reference values contribute a majority of signal *mass* because there are many of them, not because any individual near-reference participant is as informative as a clinically abnormal one. The result concerns where information sits in a population, not individual risk. It also does not imply that young-healthy intervals should replace clinical reference ranges, which serve a different purpose. The young-healthy centre is in any case not claimed to coincide with any marker's mortality-minimising value: the coordinate system measures displacement from youth, which is what is intended, and must not be read as displacement from optimum (§S5).

**Not discharged:** a held-out hazard contrast *within* the least-displaced stratum was specified as a complementary analysis and has not been rebuilt. It answers a different question — whether the projection still discriminates among people who have barely moved — and is not required for the claim above. See `MANUSCRIPT_DEFECTS.md` Part 3 for the adjudication.

#### S1b.6 Code and outputs

**Code:** `analysis/04_.../code/03_signal_zones.py` — cross-fitting at line 185, cohort reconciliation asserted at line 58

**Outputs:** `outputs/data/signal_zone_summary.csv`, `signal_zone_partitions.csv`, `signal_zone_marker_summary.csv`, `signal_zone_classification_audit.csv`, `cycle_partition_manifest.csv`, `clinical_alert_sensitivity_summary.csv`

> **FIGURE 2F** — location of positive mortality-associated signal relative to clinical limits. *Note: the Figure 2 legend currently carries pre-correction values (56.94/56.79, 67.60/65.58) and must be regenerated from `signal_zone_summary.csv`.*

---

### §S2. Dysregulation magnitude

> Manuscript section: *"Physiological dysregulation magnitude did not explain the female survival advantage"* · **Figure 2, Table 2**

#### S2.1 Hypothesis and aim

**Hypothesis:** if women survive longer because they accumulate less physiological dysregulation, then a scalar measure of dysregulation should be lower in women and should attenuate the female survival advantage on adjustment.

This is the first of the two candidate physiological explanations raised by §S1, and the section is written to be capable of confirming it. It does not.

#### S2.2 Methodological approach and why

**Two scalar measures, constructed differently on purpose.** Homeostatic dysregulation (HD) is Mahalanobis distance from the same-sex young-healthy centroid (§S9.4); the sensitised frailty index (FI) is the unweighted mean adverse excursion across 38 markers (§S9.5). Both are computed without reference to mortality.

*Why two:* HD accounts for the correlation structure of healthy physiology, so a set of individually unremarkable but jointly improbable values registers as distant. The FI cannot do this but is portable and computable from a lookup table. A conclusion holding in only one would be a property of the scoring choice. The two share participants, markers, reference population and outcome, so **agreement between them establishes robustness to encoding, not independent replication.**

**Three questions are asked in sequence:** whether each measure predicts mortality; whether women differ from men on it; and whether adjusting for it attenuates the female hazard ratio. The third is the actual test — a measure can be strongly prognostic and still explain none of a group difference.

**Adjusted sex contrasts** use ordinary least squares with a four-degree-of-freedom cubic B-spline in age and survey-cycle fixed effects, with HC3 robust intervals. Outcomes are standardised within the analysis population, so coefficients are in cohort SD units.

*Why a spline rather than linear age:* both measures rise nonlinearly with age and the sexes have slightly different age distributions; a linear term would leave curvature in the residual and load it onto the sex coefficient.

**Coefficient changes after adjustment are descriptive and are not interpreted as causal mediation.** The exposure is not randomised and the adjustment variable is a downstream physiological state.

#### S2.3 Estimation-population declaration

| Quantity | Estimated on | Why | Used for |
|---|---|---|---|
| HD reference mean, SD, covariance | Young-healthy reference, **by sex** | Distance measured relative to healthy correlation structure | HD |
| FI reference median, robust scale | Young-healthy reference, **by sex** | Robust set point for a portable index | FI deficits |
| Cox coefficients | Mortality cohort, n = 29,023 | The population the estimate describes | Table 2 hazard ratios |
| Standardisation mean, SD | The analysis population of each model | Per-SD effects must be in that population's units | All per-SD estimates |
| Adjusted sex contrast | Age ≥52 stratum and full cohort, reported separately | The mortality analyses run in the ≥52 stratum | Figure 2C |

#### S2.4 Results

**Both measures predict mortality strongly.** Per cohort SD, in 29,023 participants with 2,922 disease deaths: FI **1.71** (95% CI, 1.65–1.78) and HD **1.67** (1.61–1.73). Neither shows a sex interaction ($P$ = 0.273 and 0.746). Entered jointly they retain independent association (FI 1.29, HD 1.38; apparent C-index 0.843), indicating related but non-identical aspects of state.

**The two measures disagree about whether women are more dysregulated.** Adjusted female-minus-male difference in the age ≥52 stratum: FI **+0.190 SD** (95% CI, 0.154 to 0.225) but HD **+0.010 SD** (95% CI, −0.026 to 0.045), which is compatible with no difference.

This dissociation is itself informative, and is the first indication that "amount of dysregulation" is not a single quantity. The FI weights each marker's adverse excursion equally; HD discounts deviations that are expected given the healthy correlation structure. Women's excursions are larger in unweighted terms but not after accounting for how healthy physiology co-varies.

> **Terminology, load-bearing.** A third quantity — the Euclidean norm of the 38-coordinate vector, *m* — is reported in §S3 and is **not** the same as HD. Women are higher on *m* (+0.232 SD in HD coordinates) while showing no difference on covariance-adjusted HD (+0.010 SD). Both are true. Any sentence using "physiological distance" or "farther from young-health" must state which is meant.

**Neither measure explains the female survival advantage.** The age-adjusted female hazard ratio was **0.624**; after additional adjustment for the FI it was **0.565** (0.524–0.609), and for HD **0.609** (0.566–0.656). Both estimates move *away* from unity or barely move toward it. Adjusting for how much dysregulation a person carries does not reduce the female advantage — in the FI case it widens it, because women carry more measured burden while dying less.

**Signal location** relative to clinical limits is reported at §S1b, which uses the same module.

#### S2.5 Conclusion

Physiological dysregulation magnitude is strongly prognostic and does not explain the female survival advantage. The result is robust to the choice of scalar: a covariance-aware distance and an unweighted deficit mean agree on the conclusion while disagreeing on whether women are more dysregulated at all.

**What this establishes negatively is the point.** A scalar collapses a vector. Two participants equally far from young-health may differ in which markers moved, how those deviations covary, and whether the resulting configuration resembles the physiology associated with death. §S3 tests that possibility directly.

**What this does not establish.** It does not show that physiological distance is unimportant — both measures are among the strongest mortality predictors in the study. It shows that distance does not carry the sex difference.

#### S2.6 Code and outputs

**Code:** `analysis/04_physiological_dysregulation_magnitude/` · ranges `01_build_ranges.py` · descriptives and sex contrasts `02_descriptive_magnitude.py` · signal location `03_signal_zones.py` · survival models `04_mortality_models.py` · reporting under `code/reporting/`

*Reporting is separated from fitting by design:* `run_all.py` fits models only, so a figure or table can be rebuilt without re-running the analysis.

**Outputs:** `outputs/tables/Table_2_dysregulation_mortality.csv`, `Supplementary_Table_magnitude_mortality_full.` `outputs/figures/Figure_2_candidate_compressed.{png,pdf,svg}`, `outputs/data/biomarker_range_table.csv`

**Protocol and QC verdict:** `PROTOCOL_AND_FERMI_VERDICT.md`

> **FIGURE 2 HERE** — `outputs/figures/Figure_2_candidate_compressed.png` *Six panels: HD and FI distributions against the young-healthy reference, the adjusted sex contrast, age trajectories, the female-gap cascade, and signal location relative to clinical limits.*
>
> **TABLE 2 HERE** — `outputs/tables/Table_2_dysregulation_mortality.csv`
>
> **SUPPLEMENTARY TABLE S10 HERE** — `Supplementary_Table_magnitude_mortality_full.csv` *Magnitude–mortality models with sensitivities.*

---

### §S3. Mortality-associated direction

> Manuscript section: *"Women were less aligned with a shared mortality-associated physiological direction"* · **Figure 3**

#### S3.1 Hypothesis and aim

**Hypothesis:** if the female survival advantage is not explained by *how far* physiology has moved (§S2), it may be explained by *which way* it has moved — that is, by how much of a person's displacement lies along the multivariable direction associated with disease mortality.

The aim is to construct that direction without outcome leakage, decompose each participant's displacement against it, and test whether the resulting projection carries the sex difference that magnitude could not.

#### S3.2 Methodological approach and why

**The geometry.** For a participant with coordinate vector $\mathbf{v}$ and unit mortality direction $\mathbf{u} = \beta/||\beta||$:

$$m = ||\mathbf{v}||, \qquad a = \frac{\mathbf{v}^\top \mathbf{u}}{||\mathbf{v}||}, \qquad q = \mathbf{v}^\top \mathbf{u} = m\,a, \qquad R = \sqrt{\max(m^2 - q^2,\, 0)}$$

*Magnitude $m$* is how far the participant has moved; *alignment $a$* is the cosine of the angle between that movement and the fitted direction; *projection $q$* is how much of the movement lies along it; *orthogonal displacement $R$* is the remainder.

*Why decompose at all:* §S2 showed that a scalar summary of displacement does not carry the sex difference. The decomposition is what makes "how far" and "which way" separately measurable, which is the central claim of the paper.

$R$ **requires the full 38-dimensional space.** Computed inside a panel selected for alignment with $\mathbf{u}$, there is almost no orthogonal remainder by construction, so $R$ is only interpretable at full dimension.

**Preventing circularity — the central methodological problem of this section.** The direction is fitted *against the outcome it is then used to predict*. Three mechanisms separate them:

1. **Leave-one-survey-cycle-out fitting.** Every participant's $q$, $a$ and $R$ are computed from a direction estimated on the other nine cycles. No participant's coordinates derive from any outcome in their own cycle.
2. **Whole cycles, not individuals, as the grouping unit.** Splitting by individual would let the model exploit cycle-specific assay and calibration artefacts, which inflates apparent generalisation. Grouped splitting forces transfer across a change of measurement era.
3. **Nothing is refitted at evaluation.** The direction, its scaling, and every derived coordinate are frozen before the held-out outcome is read.

*What this does and does not buy.* It removes resubstitution optimism and direct outcome leakage. It does **not** make $q$ outcome-independent, causal, or a validated surrogate. $q$ is proportional to the biomarker component of a fitted Cox linear predictor; it is not an estimate of absolute risk.

**Two representations throughout.** The entire analysis runs in both HD and signed-FI coordinates. As in §S2 they share participants, markers, reference and outcome, so agreement is robustness to encoding rather than replication.

**Time scale.** Attained age with delayed entry at examination, cycle-stratified baseline hazards, Efron ties. *Why attained age:* age is the dominant determinant of hazard, so using it as the time scale compares participants at the same age rather than the same follow-up duration and removes the need to model the age–hazard relationship parametrically.

**Uncertainty.** 500 participant bootstrap replicates, resampled within sex-by-cycle strata, with **all leave-one-cycle-out directions refitted inside every replicate** — so the interval propagates uncertainty in the direction itself, not only in the final coefficient. Split stability is assessed separately across all 126 unique five-versus-five cycle partitions; those percentiles describe sensitivity to cycle allocation and **are not confidence intervals**.

**Two intervals exist for the sex contrasts, and the manuscript quotes the wider one.** `sex_contrasts.csv` carries HC3 intervals *conditional on the cross-fitted direction* — they treat the fitted direction as if it were known in advance. `bootstrap_summary.csv` carries the direction-refit bootstrap. For magnitude the two agree, because magnitude does not depend on the direction at all; for alignment and projection the bootstrap is **four times wider**, because those quantities are defined relative to a direction that was itself estimated:

| Quantity (HD) | HC3, conditional | Direction-refit bootstrap | Width |
|---|---|---|---|
| Magnitude $m$ | (0.197, 0.268) | (0.199, 0.268) | 1.0× |
| Alignment $a$ | (−0.766, −0.703) | (−0.836, −0.569) | 4.2× |
| Projection $q$ | (−0.676, −0.610) | (−0.752, −0.488) | 4.0× |

The manuscript reports the bootstrap intervals. The sex-contrast panel (Figure 2c in the manuscript) is drawn with the conditional ones and its legend states this. The conclusion is identical either way — every interval excludes zero by a wide margin — but the conditional interval understates uncertainty for the two quantities that depend on the fitted direction, so it must not be the one quoted. **The `uncertainty_status` field inside `sex_contrasts.csv` still reads "full direction-refit bootstrap pending"; that string is stale, the bootstrap completed 500/500 valid replicates.**

#### S3.3 Estimation-population declaration

| Quantity | Estimated on | Why | Used for |
|---|---|---|---|
| Direction $\beta$, hence $\mathbf{u}$ | Nine training cycles, excluding the scored participant's cycle | Prevents outcome leakage into that participant's coordinates | $q$, $a$, $R$ |
| Score standardisation | Training cycles only | Test-set scaling would leak outcome-adjacent information | Per-SD estimates |
| Cox coefficients for reported hazard ratios | Full age 52–79 cohort, on frozen cross-fitted scores | The population the estimate describes | Table of associations |
| Bootstrap resampling | Participants within sex × cycle strata | Preserves the design that generated the data | Interval estimates |
| Split percentiles | 126 five-versus-five partitions | Sensitivity to cycle allocation | Stability ranges |

#### S3.4 Results

Cohort: **11,497** participants aged 52–79 (5,920 men, 5,577 women) with **2,494** disease deaths.

**Women are farther from young-health but less aligned with the mortality direction.** Adjusted female-minus-male differences, in cohort SD units:

| Quantity | HD | FI |
|---|---|---|
| Magnitude $m$ | **+0.232** (0.197 to 0.268) | **+0.140** (0.104 to 0.176) |
| Alignment $a$ | **−0.735** (−0.766 to −0.703) | **−0.743** (−0.776 to −0.711) |
| Projection $q$ | **−0.643** (−0.676 to −0.610) | **−0.578** (−0.612 to −0.545) |

The configuration is preserved in **every one of the 252 directed cycle partitions** in both encodings; none reverses it. Note that $m$ here is the Euclidean coordinate norm and is **not** the covariance-adjusted HD of §S2, on which women and men do not differ; both statements are true of the same participants.

**Both magnitude and projection predict mortality, projection more strongly.** Per cohort SD in HD coordinates: $m$ **1.63** (1.57–1.69), $q$ **1.84** (1.77–1.90).

**Adjustment for projection attenuates most of the female–male log-hazard difference; adjustment for magnitude attenuates none of it.** Female hazard ratio in the same cohort. Intervals are direction-refit participant bootstrap intervals, the uncertainty specified in Methods — **not** the conditional Cox intervals, which treat the fitted direction as if it were known in advance and are narrower:

| Model | Female HR (95% bootstrap interval) | log-HR |
|---|---|---|
| Sex only | 0.624 (0.573–0.678) | −0.472 |
| + magnitude $m$ | 0.532 (0.484–0.578) | −0.631 |
| + magnitude and alignment | 0.813 (0.703–0.890) | −0.207 |
| **+ projection $q$** | **0.941 (0.815–1.047)** | **−0.061** |

*Read the attenuation as a coefficient change, not as mediation.* Projection is outcome-supervised and may lie downstream of both sex and mortality-related processes, so the change on adjustment is descriptive. It does not estimate the proportion of the survival advantage explained, and non-collapsibility alone can move a hazard ratio on adjustment.

Adjustment for magnitude moves the estimate **away** from unity, reproducing §S2 in the direction analysis. Adjustment for projection removes **87%** of the sex difference on the log-hazard scale in HD coordinates (−0.472 → −0.061) and 78% in FI coordinates, where the post-projection hazard ratio is 0.900 with a bootstrap interval of [0.783, **0.997**] — still excluding 1, but a boundary case rather than a clear residual.

**The direction is substantially shared between the sexes.** Directions estimated separately within each sex, in sex-specific young-healthy relative coordinates, induce scores in held-out participants correlating at **0.868 to 0.910**. Applying the direction trained in the *other* sex retained **98.6% to 104.7%** of the own-sex log-hazard association, and allowing sex-specific directions did not improve pooled held-out partial log likelihood. Estimating one direction for both sexes cost nothing detectable.

This is the result that makes a single screening instrument coherent. It does not imply identical physiology: women and men differ in raw distributions, disease prevalence, exposures, baseline hazard and their positions within the space. **Shared direction, different position.**

**Two qualifications on how $q$ should be read.** Its association with mortality is stronger in women than men (per-SD hazard ratios 2.068 against 1.764; multiplicative ratio 1.174 in both encodings), so women sit lower on the axis but respond to it more steeply. And the association declines with attained age during follow-up, from 2.427 in the 52–64 band to 1.331 at ages 85 and above, remaining above unity throughout. Proportional-hazards diagnostics detected age-varying projection effects, so the reported per-SD hazard ratios are **cohort-averaged associations rather than constant multipliers**.

**Post-correction changes.** After the FI correction (§0.4 defect 2) the projection association is nonlinear in **both** representations (FI $P = 3.8 \times 10^{-7}$; it had previously been reported as HD-only), and HD–FI participant-level agreement rose sharply — correlations ∼0.83 to ∼0.98, four-region kappa 0.55 to 0.80. The version-1 thresholding had been *understating* agreement between representations. No conclusion reversed.

#### S3.5 Conclusion

Women and men move within a substantially shared mortality-associated physiological geometry, but women occupy a region of it with less displacement along the mortality-associated direction. That projection accounts for most of the female survival advantage, where total displacement accounts for none of it.

**What this does not establish.** The direction is predictive, not mechanistic. It may reflect causal ageing processes, accumulated disease, treatment, exposure history, preclinical illness, compensation, or unmeasured confounding. Lower alignment must not be equated with good health in an absolute sense: displacement lying away from the fitted direction may still produce disease, disability or mortality through pathways the model does not represent. The study identifies *where* the dominant sex contrast lies within measured physiology; it does not establish *why* women and men occupy different positions.

#### S3.6 Code and outputs

**Code:** `analysis/05_mortality_associated_direction/` · contract in `ANALYSIS_CONTRACT.md` (Amendment 1 adopted the corrected FI input) · `08_split_stability.py`, `09_bootstrap_uncertainty.py`, `10_sensitivities.py`

**Outputs:** `outputs/data/sex_contrasts.csv`, `mortality_associations.csv`, `cross_sex_direction_transfer.csv`, `loco_participant_geometry.csv`, `loco_direction_coefficients.csv`, `bootstrap_summary.csv`, `ph_diagnostics.csv`, `prespecified_sensitivity_summary.csv`, `representation_region_agreement.csv`

**Prose and checkpoints:** `outputs/REVISED_RESULTS_SECTION_DRAFT.md`, plus `SHARED_DIRECTION_`, `SPLIT_STABILITY_`, `SENSITIVITY_` and `BOOTSTRAP_UNCERTAINTY_CHECKPOINT.md`; gate assessment in `GATE_14_EVALUATION.md`

**Verification:** 11/11 scripts exit 0, **49/49 tests pass**, 6/6 locked input hashes verify, 500/500 bootstrap replicates valid.

> **A silent-failure class was found and fixed here.** The bootstrap originally resumed on replicate number alone, with no input-hash key, so a rerun would have silently reused all 500 superseded replicates and passed every check. Checkpoints now carry an input-lock fingerprint — see `IMPLEMENTATION_NOTE_001.md`. Any long-running resumable job should be checked for the same pattern before its resume is trusted.
>
> **FIGURE 3 HERE** — participant-level geometry, adjusted sex contrasts, shared-direction adequacy, mortality associations, and the female-advantage cascade.

---

### §S4. Reduced-panel development

> Manuscript section: *"A ten-marker panel reproduces the mortality-associated direction"* · **Figure 4, Table 3**

#### S4.1 Hypothesis and aim

A 38-assay panel is not a deployable screening instrument. The aim is to determine how much of the mortality-associated direction survives reduction to a routinely measurable subset, which markers carry it, and at what panel size the reduction becomes inadequate.

**The question is sufficiency, not ranking.** A marker may be strongly associated with mortality in isolation yet redundant once others are present, and a marker of modest individual association may be necessary to reconstruct the multivariate direction. Univariate screening answers a different question and is not used.

#### S4.2 Methodological approach and why

**Selection.** Elastic-net penalised Cox partial likelihood (§S9.7), with sex included unpenalised, markers ranked by absolute penalised coefficient and marker name as a deterministic tie-break.

*Why elastic net rather than lasso:* biomarkers are correlated, and pure $L_1$ selects arbitrarily among correlated predictors. The $L_2$ component stabilises that choice while $L_1$ still produces sparsity. The mixing fraction is tuned, not assumed.

*Why coefficient-magnitude ranking is legitimate here:* HD coordinates share a common scale — standard deviations of the same-sex young-healthy reference — so coefficients are comparable across markers. This would not hold for arbitrarily scaled predictors.

**Preventing overfitting — nested, cycle-grouped validation.** The ten survey cycles are divided into two halves of five intact cycles. Within the outer training half, **an inner grouped cross-validation tunes the penalty and mixing fraction, and the tuned model ranks the 38 markers**; the top-$k$ are refitted in the outer training data and applied unchanged to the outer test cycles. Every one of the 126 unique five-versus-five divisions is evaluated in both directions, giving **252 directed evaluations**.

Confined to the outer training cycles: marker selection, hyperparameter tuning, coefficient estimation, and score scaling. The held-out cycles contribute only their outcomes, and only at evaluation.

*Why nested rather than a single split:* tuning on the same data used to report performance is optimistically biased. The inner loop pays that cost internally, so the outer estimate is not inflated by hyperparameter search.

*Why the complete enumeration of 126 partitions rather than a sample:* it removes any possibility of a favourable draw and characterises the lower tail better. A 20-partition subset gives near-identical medians (0.9011 against 0.9018), so **the full enumeration did not change the answer** — its value is no-cherry-picking and a better lower tail (2.5th percentile 0.8677 against 0.8807), and it should be defended on those grounds rather than on a different result.

**Adequacy is decided by a frozen multi-criterion rule.** Eight criteria across four requirements, with thresholds fixed before any result was generated: reproduction of the full 38-marker projection (median and 2.5th percentile), transfer of that projection to the alternative encoding (median and 2.5th percentile), retention of the full panel's held-out mortality information, positive association within each sex separately, and preservation of the female–male contrast.

*Why thresholds were frozen first:* a narrow failure must return the larger panel rather than invite reconsideration of the threshold. This matters here, because the binding criterion passes by 0.0018.

**Panel size versus panel identity — a distinction the contract enforces.** Nested outer validation establishes the performance of *a selection algorithm at a size*, not an unbiased estimate for one exact marker combination. Only after the adequate size was fixed was the frozen algorithm applied **once** to the complete development cohort — tuning by ten-fold leave-one-cycle-out, refitting on all ten cycles, ranking all 38, and locking the top $k$. **The resulting identity is a locked development product requiring external validation; its apparent performance in the same cohort is not an unbiased estimate and is never presented as one.**

Cross-partition selection frequency is retained but is **descriptive only** and was barred from determining panel size. Any 90% line shown is for historical continuity and is labelled as such.

#### S4.3 Estimation-population declaration

| Quantity | Estimated on | Why | Used for |
|---|---|---|---|
| Penalty, mixing fraction | Inner grouped folds within outer training cycles | Tuning on evaluation data is optimistically biased | Marker ranking |
| Marker ranking and selection | Outer training cycles only | Held-out evaluation requires it | Candidate panels |
| Refitted coefficients | Outer training cycles only | Applied unchanged to test cycles | Held-out projections |
| Score scaling | Outer training cycles only | Test-set scaling leaks outcome-adjacent information | Reproduction correlations |
| Locked identity | **All ten cycles, once, after size was frozen** | §S9/§10 requires a full-development fit for identity | Table 3B |
| Selection frequency | 252 directed evaluations | Descriptive stability only | Figure 4D |

#### S4.4 Results

**Ten markers is the smallest adequate size; seven is not.** Reproduction improved monotonically with size. Seven markers reproduced the full projection at a median held-out correlation of 0.858 and retained 0.824 of its mortality information, failing three of the eight criteria. Ten satisfied all eight, reproducing at **0.902** (2.5th partition percentile 0.868) and retaining **0.913** of its mortality information (Table 3A).

**The binding margin is narrow and is reported openly:** median reproduction **0.9018** against a threshold of 0.90 — a margin of 0.0018. Every other criterion at size ten had substantial headroom. Because thresholds were fixed in advance, ten is the size the procedure returns; it is not established as a universal optimum.

**Run quality.** 252 of 252 tasks complete, **zero fit failures and zero solver warnings across 3,528 fits**, 174 CPU-hours.

**The locked panel**, in full-development rank order: alanine aminotransferase, red cell distribution width, mean corpuscular volume, neutrophil count, gamma-glutamyl transferase, urine albumin-to-creatinine ratio, platelet count, albumin, phosphorus, pulse rate (Table 3B; penalty 0.001, $L_1$ fraction 0.25). It spans liver, red cell, immune, renal, platelet, mineral and cardiovascular domains and requires nothing beyond routine biochemistry, haematology and urinalysis.

**Two independent calculations agree on where the panel stops being solid.** Ranks 1 to 8 of the full-development lock are exactly the eight markers selected in at least 90% of the 252 partitions; ranks 9 and 10 are exactly the two selected least consistently (phosphorus 56%, pulse 64%). Ranking all 38 markers by selection frequency and taking the top ten **recovers the locked ten exactly**, with the tenth marker selected 2.6 times more often than the eleventh (0.556 against 0.210). Rank order agrees at Spearman $\rho$ = 0.891, with disagreement confined to the two least stable slots, which swap.

*This is a consistency check, not replication.* Both calculations use the same algorithm, coordinates and data, and the full-development fit trains on a superset of every training half. What it establishes is that the single full-sample fit picked up nothing idiosyncratic to being run once on everything.

**Reduction amplified the sex contrast.** The adjusted female–male difference per standard deviation was larger for the reduced panel than for all 38 markers at every size, reaching **1.25 times** the full-panel contrast at ten. Attributing the full-panel contrast across markers accounts for it: the ten selected markers contributed **108%**, and the remaining twenty-eight contributed **−8%**, collectively displacing the contrast toward zero. The largest opposing markers — uric acid, alkaline phosphatase, chloride, lactate dehydrogenase, systolic pressure — are ones on which men deviate further from their own young-healthy reference than women do.

Reduction therefore did not merely discard uninformative markers; it removed markers that actively opposed the sex contrast. A panel assembled to maximise coverage of physiological change would have diluted precisely the signal this study concerns.

**The frailty-index encoding required fifteen markers, and the reason generalises.** Its smallest adequate panel was 15, not 10. This is not a transfer failure: the ten HD-selected markers reproduced the FI projection at a median held-out correlation of **0.887**. Nor did the encodings select different biology — **the ten are a strict subset of the fifteen**, and at equal size the two panels share eight of ten markers with identical first and second ranks, the FI substituting aspartate aminotransferase and red-cell folate for albumin-to-creatinine ratio and phosphorus. Sign differences on shared markers are entirely the FI orientation convention; there are **zero substantive sign conflicts**.

The difference is **informative-measurement coverage**. HD coordinates are signed, so every marker carries a value for every participant and ten markers supply ten measurements. The FI scores only adverse excursions, so a participant on the healthy side of a marker receives exactly zero and that marker contributes nothing to their score. Across all 38 markers, **21%** of participant-marker values are zero on this account. Across the ten markers the FI itself selects, the average participant carries **7.27** informative values; across its fifteen, **11.10** — the smallest panel at which the index delivers as many usable measurements per participant as ten HD markers. The additional five markers do not introduce new biology; they compensate for what the encoding discards.

*This is a real trade-off rather than a dominated choice.* Using all 38 markers, the adverse-only encoding predicted mortality marginally **better**. Rectification sharpens each marker while thinning each participant's profile, and these act in opposite directions on prediction and on compressibility.

> **State the panel size whenever quoting selection frequency.** ACR is selected in 100% of HD partitions but only 25% of FI partitions **at size 10**, which reads as severe representation-dependence —

yet at the FI's own adequate size of 15 it is selected in **94%** and is in the locked FI identity. The gap is mostly budget, not disagreement. `Table_S_marker_comparison.csv` carries both columns.

**Prespecified sensitivities — all six complete.**

| # | Sensitivity | Result |
|---|---|---|
| 12.1 | Age floor 50 / 52 / 55 | Reproduction 0.920 / 0.918 / 0.915; sex contrast negative throughout. **Age 52 is not load-bearing.** In-sample, so not comparable to Table 3A |
| 12.2 | Time scale | 0.918 under attained age, 0.906 under time-since-examination with a 4-df age spline; female > male preserved |
| 12.3 | Ridge-dominant penalty | No marker's selection frequency shifts by more than 0.016. Slots 9–10 are **not** an $L_1$-sparsity artefact |
| 12.4 | Leave-one-cycle-out geometry | **The two metrics move in opposite directions** — see below |
| 12.5 | Historical comparison | **9 of 10 markers shared**, hyperparameters identical. The sole disagreement is rank 9, which the historical output also flagged as unstable |
| 12.6 | Outcome-blind clusters | Cluster-level agreement (0.583) now exceeds marker-level (0.538), so correlated substitution explains part of the HD/FI disagreement — but absolute agreement fell relative to version 1. Report both halves |

**Sensitivity 12.4 is the one that constrains interpretation.** Under a nine-cycles-train, one-held-out design, reproduction is *better* (0.9169 against 0.9018) but retained mortality information is *worse* at every panel size (0.876 against 0.913 at size ten) — below the 0.90 primary threshold. The mechanism is that the five-versus-five geometry trains the full-38 comparator on only five cycles, so the comparator is noisy, its held-out gain is small, and the retained *ratio* is inflated. **The five-versus-five geometry flatters the retained-likelihood criterion by handicapping the comparator**, and the estimate reported here is therefore not conservative with respect to validation design. This is not an adequacy test — ten markers stands — but it must be stated.

#### S4.5 Conclusion

Ten routine measurements reproduce most of the mortality-associated direction defined by 38, retain most of its prognostic information, and express the sex contrast more strongly than the panel they were drawn from. Two encodings converge on a nested set of markers from different scoring rules and different adequate sizes, which argues the selected biology is a property of the data rather than of the scoring choice.

**Known limitations, stated rather than discovered.** Panel *size* was validated in held-out cycles; the exact ten-marker *identity* was not, and its apparent performance here is not unbiased. Candidate sizes were locked at 3, 5, 7, 10, 12 and 15, so **sizes 8 and 9 were never tested** and the adequacy boundary between 7 and 10 is unresolved — while the stable core is 8. Testing them now would be post-hoc and could not become the primary result. Selection used one algorithm; a different penalty or ranking rule could reach adequacy at a different size.

Marker coefficients are attributions within a fitted multivariable model and do not identify causal biomarkers, mechanisms or intervention targets.

#### S4.6 Code and outputs

**Code:** analysis/06_biomarker_selection/ · contract `ANALYSIS_CONTRACT.md` · `02_nested_selection.py` (parallel, resumable) · `code/reporting/02_full_development_lock.py` (identity lock, `--verify` mode) · four implementation notes

*Reporting code lives in `code/reporting/` by design:* the analysis fingerprint globs `code/*.py` non-recursively, so this keeps the fingerprint keyed to the fitting computation and leaves the 252 checkpoints addressable.

**Outputs:** `Table_4_adequacy_by_panel_size.csv`, `Table_4_adequacy_decision.csv`, `Table_4_locked_identity{,_FI}.cs`[illegible] `Table_S_{marker,subsystem}_comparison.csv`, `Table_S_cross_transfer.csv`, `selection_stability.csv`, `Supplementary_Table_S_sensitivities.csv`, `Figure_4_biomarker_selection.{png,pdf,svg}`

**Verification:** fingerprint `0B542088E190536BFEC3`; 11/11 tests pass; `02_full_development_lock.py --verify` re-applies the re-expressed tuning rule to the stored per-cell likelihoods of all 252 HD evaluations and reproduces **252/252** stored hyperparameter choices exactly.

> **Reuse of invariant computation was gated, not assumed.** The rerun on corrected FI input reused the HD branch's `tune_and_rank` output, cutting 176.9 CPU-hours to 1.92. Reuse is permitted only by `outputs/audit/selection_reuse_certificate.json`, produced by a script that recomputes `tune_and_rank` from scratch and compares decision-bearing columns exactly and refitted floats within 1e-10. Verdict `reuse_permitted`, largest deviation 3.55e-15. `IMPLEMENTATION_NOTE_004.md` records **two mis-specified versions of that test that wrongly refused valid reuse** — read it before writing another verification of this kind.
>
> **TABLE 3 HERE** — (A) adequacy by panel size, (B) the locked ten-marker panel.
>
> **FIGURE 4 HERE** — design, adequacy by size, transfer between encodings, marker stability and the final panels.
>
> **SUPPLEMENTARY TABLES S2–S5, S11 HERE** — selection stability, subsystem span, cross-transfer, sensitivities, and the HD/FI marker comparison at sizes 10 and 15.

---

### §S5. Adverse-direction assignment and its validation

> Underpins the sensitised frailty index (§S9.5) and every quantity derived from it. No manuscript Results section; the assignment is described in Methods under *"Sensitised frailty index"*.

#### S5.1 Why this section exists

The sensitised FI scores the **adverse excursion** of each marker, which requires every marker to be designated high-adverse, low-adverse or bidirectional before any score is computed. Every deficit-based index makes this assumption, and it is usually left implicit.

It is stated explicitly here because it is load-bearing in two directions. It determines which half of each marker's distribution contributes to a participant's score, and — because the FI is one of the two representations the whole analysis is run in — it determines whether agreement between representations means anything. An assignment derived from the outcome would make the "independent" second encoding a second look at the same fit.

#### S5.2 Methodological approach and why

**Adverse direction is a clinical prior, fixed before scoring.** Each of the 38 markers carries four locked fields in `config/fi_scoring_spec.csv`:

| Field | Meaning |
|---|---|
| `adverse_direction` | `high_only`, `low_only` or `two_sided` |
| `orientation` | `−1` where the locked transformation reverses the raw-value axis, `+1` otherwise |
| `physiological_system` | locked subsystem assignment |
| `direction_basis` | a written physiological justification for the assigned direction |

The resulting distribution is **19 high-only, 15 two-sided and 4 low-only**, with five markers carrying `orientation = −1`.

**The governing rule, quoted from the locked methods document:**

> *Directionality must not be selected or revised using associations with age, sex, mortality, disease outcomes or the observed transformed-data distribution. Any correction made during review must be justified from the biomarker's physiological interpretation and documented before FI scores are calculated.*

*Why this rule and not an empirical one:* deriving adverse direction from mortality in this cohort would make the coordinates outcome-selected before any cross-validation began. Every held-out evaluation downstream would then be reporting performance for a coordinate system that had already seen the outcome, and no amount of subsequent cycle-grouping would recover the independence. The assignment is therefore made on physiological grounds and is **outcome-blind by construction**, which is a weaker claim than "correct" but is the claim that keeps the analysis honest.

**Direction is defined on the original measurement scale**, not the transformed one, and is not re-estimated when a transformation changes. During standardisation, multiplication by the locked `orientation` restores transformed coordinates to the raw-value axis before the adverse-tail rule is applied (§S9.3), so a reciprocal-transformed marker is not silently reversed.

**Every assignment carries a written basis.** Examples, quoted verbatim from the specification:

- *high gamma-glutamyl transferase reflects cholestasis or alcohol-related injury*
- *hyponatraemia and hypernatraemia are both adverse*
- *high pulse reflects sympathetic overdrive or poor fitness; the low tail is clinically ambiguous without fitness or rhythm data*
- *hypoalbuminaemia reflects inflammatory or hepatic dysfunction; high albumin commonly reflects dehydration*

The complete set of 38 is Supplementary Table S15.

#### S5.3 Decision record

Reconciling the reviewed FI specification against the general biomarker registry found **two disagreements**, both resolved before scoring and both recorded:

| Marker | Registry | FI specification | Resolution and basis |
|---|---|---|---|
| Diastolic blood pressure | `two_sided` | `high_only` | Retained `high_only` — hypertension is the prespecified diastolic disease axis |
| Pulse rate | `two_sided` | `high_only` | Retained `high_only` — low resting pulse is clinically ambiguous without fitness and rhythm information |

Both are physiological, outcome-blind decisions and are recorded in the specification's `direction_basis` field.

#### S5.4 Validation

The specification is locked by SHA-256 and checked by an executable gate before any FI value is calculated. The gate verifies that the transformed panel and transformation recipe are the expected locked files, that all 38 markers are present in panel order, that every marker has exactly one valid `adverse_direction` category and one subsystem, and that transformation and orientation match the locked transformation recipe rather than the superseded registry field.

**Code:** `src/10_validate/validate_fi_scoring_spec.py`, with regression tests in `tests/test_fi_scoring_spec.py`. Passing the gate completes the specification step; it does not calculate reference parameters or participant FI values.

#### S5.5 Conclusion, and an acknowledged limitation

Adverse directions in this study are prespecified clinical priors, individually justified, hash-locked, validated by an executable gate, and estimated from no outcome. That is what makes the frailty-index encoding usable as a second representation rather than a restatement of the first.

**The limitation is real and should not be glossed.** A clinical prior is an assumption, not a measurement. Whether these particular assignments match the empirical hazard relationship in an older population is a separate question, and one this study is not positioned to answer without circularity: the only outcome available to test them against is the outcome the index is used to predict.

**An empirical diagnostic of exactly this question was performed and is reported in the companion manuscript**, where nothing depends on its result. It is not reported here, and no conclusion in this paper depends on it. Readers should treat the directionality assignments as prespecified priors with stated physiological justification, and should note that the young-healthy reference centre is **not** claimed to coincide with any marker's mortality-minimising value — the coordinate system measures displacement from youth, which is what is intended, and must not be read as displacement from optimum.

#### S5.6 Code and outputs

**Specification:** `data_preparation/config/fi_scoring_spec.csv` (38 rows, six fields, SHA-256 locked)

**Documentation:** `FI_METHODS.md` Step 1 — requirement, governing rule, validation gate and decision record

**Validation:** `src/10_validate/validate_fi_scoring_spec.py`, `tests/test_fi_scoring_spec.py`

**Applied at:** `src/11_fi/build_fi_adverse_excursions.py` (§S9.5)

> **SUPPLEMENTARY TABLE S15 HERE** — `data_preparation/config/fi_scoring_spec.csv`
> *Adverse-direction assignment for all 38 biomarkers, with transformation, orientation, physiological system, and the written physiological basis for each assigned direction.*

---

### §S6. Trial-arm sorting

Manuscript section: *"A single biomarker geometry sorts participants into two trial arms"* · **Figure 5, Table S7**

**This section replaces the two separate recruitment screens described in earlier drafts.** It supersedes `github/section_6_panel_screening` and `github/healthspan_screen`, neither of which was rebuilt and both of which predate the FI correction.

#### S6.1 Hypothesis and aim

Lifespan and healthspan trials face different recruitment bottlenecks. The first needs participants who will accrue events within a feasible horizon; the second needs participants whose physiology is already disturbed but who are not close to death, so that there is function left to preserve.

The geometry of §S3 separates exactly these two quantities. The aim is to test whether **one** classification of the $(q, R)$ plane can serve both, and to characterise the populations it produces.

#### S6.2 Methodological approach and why

**The rule**, fixed before any result was generated:

| Assignment | Rule |
|---|---|
| **Lifespan arm** | $q \geq$ training 90th percentile |
| **Healthspan arm** | $q \leq$ training 60th percentile **and** $R \geq$ training 60th percentile |
| **Neither** | all remaining participants |

Thresholds are estimated within training cycles only and applied unchanged to the complementary held-out cycles, across the same 126 partitions in both directions used for panel selection. The arms cannot overlap.

**Thresholds were carried over, not searched.** $Q_{\mathrm{LOW}}$ and $R_{\mathrm{HIGH}}$ come from a healthspan rule that had already passed all nine of its criteria, and $Q_{\mathrm{HIGH}}$ from the percentile every training half independently chose in the earlier lifespan screen. *Why this matters:* searching thresholds against the outcome would convert a frozen rule into a fitted one, and the enrichment reported would then be an in-sample maximum.

**No second scoring layer.** The sort uses the cross-fitted geometry directly; no weighted marker score is fitted on top of the panel. This differs from the historical lifespan screen, which thresholded a fitted score, so the lifespan arm here is a related but **not identical** rule — any divergence in enrichment is a consequence of that simplification and is reported as such rather than repaired.

**What is and is not independent of mortality — stated explicitly.** The transformed values, the young-healthy reference parameters and the magnitude $m$ are not derived from mortality. The direction $\mathbf{u}$, and hence $q$ and $R$, **are**. The lifespan/healthspan distinction *is* the distinction between displacement toward that direction and away from it, so it cannot be constructed without it.

**The defensible claim is therefore leakage-freedom, not mortality-independence:** directions are fitted leave-one-survey-cycle-out, so no participant's coordinates or assignment derive from outcomes in their own cycle, and thresholds are set in training cycles and transferred unchanged.

**The sort is purely geometric.** Clinical eligibility is applied afterwards as a transparent filter rather than built into the classifier, so both the geometric assignment and the post-filter cohort are visible.

**Sex was measured after assignment** and was never used to select or adjust a threshold, a panel or a representation.

**Five-year disease mortality is an incidence rate, not a cumulative-incidence probability.** The numerator is disease deaths occurring within five years of examination; the denominator is person-time truncated at five years, so a participant contributes $\min(extfollow - up, 5)$ person-years. Follow-up is measured from examination (`permth_exm`). Arm-versus-cohort comparisons are therefore **rate ratios**, computed with the same estimator in numerator and denominator.

*Why a rate rather than a proportion:* **23.8% of the cohort has less than five years of follow-up**, because later survey cycles were linked to mortality for a shorter period. A crude deaths-per-participant proportion would count those participants as full denominators and understate the rate. Person-time truncation handles administrative censoring exactly.

*Competing events.* Accidental deaths are not events; they contribute person-time up to the moment of death and none afterwards, which is the correct handling for a rate and requires no competing-risks estimator. The empirical impact is negligible: of 2,539 deaths in the cohort, **45 are accidental and 10 of those occur within five years**, removing **22 person-years of 52,363 — 0.042% of the denominator**. No death in this cohort has a missing underlying cause, so the requirement that cause be known excludes nobody.

#### S6.3 Estimation-population declaration

| Quantity | Estimated on | Why | Used for |
|---|---|---|---|
| Participant $q$, $R$ | Direction fitted leave-one-cycle-out | No assignment uses an own-cycle outcome | Plane coordinates |
| Percentile thresholds | Training cycles only | Applied unchanged to held-out cycles | Arm assignment |
| Arm event rates | Held-out participants | The estimate must not describe the fitting data | Enrichment |
| Sex composition | Held-out participants, **after assignment** | It is an outcome of the sort, not a criterion | Reported composition |

#### S6.4 Results

**The classification separates the arms cleanly.** The lifespan arm comprised approximately 10% of the eligible cohort in every configuration (median n = 573–582) at a five-year disease-mortality rate **3.7 to 4.3 times** the cohort. The healthspan arm comprised 13–17% (median n = 745–990) at **0.44 to 0.50 times** — actively depleted of near-term death — while containing a median of **three to five** displaced physiological systems. The healthspan arm was correctly positioned on both axes in 99–100% of the 252 partitions, and a usable rule was produced in **100%** of them.

**The arms have opposite sex composition.** Against a cohort that was 48.5% women, the lifespan arm was **15–28% women** and the healthspan arm **53–60%**. Neither was constructed with reference to sex.

This follows from §S3: women sit lower on $q$ at equivalent total displacement, so a threshold on $q$ selects proportionally fewer of them, while a rule selecting displacement orthogonal to $q$ selects proportionally more. The effect scales with how sharply a configuration resolves the axis — the HD panels, whose sex contrast on $q$ is the larger, produced the most male lifespan arms (14.9% and 16.7% women) while the FI panels produced 25.9% and 27.8%. **The configurations that measure the sex difference most precisely are those that most exclude women from a mortality-enriched arm.**

**This is not the screen failing in women.** Within the lifespan arm under the ten-marker HD panel, selected women had a five-year rate of **81.0** per 1,000 person-years against **51.9** in selected men, and were enriched **8.2-fold** over cohort women against **3.1-fold** for men — 0.31 five-year deaths per participant recruited against 0.22. A pooled threshold on a distribution where the sexes differ takes only the extreme tail of the lower-sitting sex, so the women it returns are the highest-risk participants in the cohort.

**A within-sex threshold restores balance at almost no cost in events.** Applying the same 90th percentile within each sex — post-hoc, reported as a sensitivity, and **barred by the specification from becoming primary** — produced arms that were 48.2–48.6% women at the same total size (median n = 577). The composition of risk changed rather than its quantity: selected women became less extreme (rate 43.1, enrichment 4.3-fold) and selected men more extreme (67.3, 4.0-fold), because the pooled rule had been admitting roughly 17% of men rather than 10%. Total expected events were nearly unchanged — **131** against 133.

#### S6.5 Conclusion

One geometric rule allocates participants to a mortality-enriched arm and a mortality-depleted, physiologically displaced arm, reproducibly and without outcome-specific tuning of either.

**What this does not establish.** Mortality enrichment does not imply treatment responsiveness: selected participants may not share an intervention's target mechanism, their risk may not be reversible, and their inclusion does not guarantee a larger treatment effect. **Whether allocation by this geometry predicts differential response is the assumption the entire enrichment strategy rests on, and it is untested.** The healthspan arm's orthogonal displacement is unassociated with mortality *in this model, cohort and panel* — it is not thereby shown to be biologically benign, and the observed low mortality does not establish intervention safety. Assignment is a relative, analysis-defined classification, not a clinical eligibility determination. Thresholds are percentiles of a training distribution and would require recalibration in any new population. Participants below age 52 were outside the geometry and were not evaluated. Five-year event counts are projections from observational follow-up, not trial accrual.

#### S6.6 Why a pooled threshold produces a male arm

The sex composition of the lifespan arm is the manuscript's strongest trial-design result, and it is arithmetic rather than biological. This subsection records the quantity that makes it legible.

**A pooled percentile is not the same percentile in each sex.** Women sit lower on the projection axis by 0.64 standard deviations, so a single cut placed at the pooled 90th percentile falls at a different point within each sex's own distribution. Computed per partition across all 252 partitions in all four configurations, with none dropped:

| Configuration | Pooled cut, as a percentile among **men** | ...among **women** | Men admitted | Women admitted |
|---|---|---|---|---|
| HD, 10 markers | 83.6 | 96.9 | 16.4% | 3.1% |
| HD, 15 markers | 83.7 | 96.5 | 16.3% | 3.5% |
| FI, 10 markers | 86.1 | 94.3 | 13.9% | 5.8% |
| FI, 15 markers | 85.8 | 94.7 | 14.2% | 5.3% |

**A rule that never mentions sex therefore acts as a sex filter.** It asks who is highest on the axis; on an axis where the sexes are separated, that question is partly a question about sex. This is a property of the threshold, not a defect of the instrument — selected women have a *higher* event rate than selected men under the pooled rule (81.0 against 51.9 per 1,000 person-years).

**The within-sex remedy, and its price.** Applying the same percentile within each sex admits exactly 10% of each, restoring the cohort's own sex composition. The exchange is explicit: marginal men leave the arm and women enter it in almost equal number (HD-10: 192.5 men out, 195.0 women in, medians across partitions). Because those women remain enriched 4.3-fold over cohort women, the arm loses only a little of its expected event count:

| Configuration | Expected 5-year deaths, pooled | within-sex | change |
|---|---|---|---|
| HD, 10 markers | 133 | 131 | −1.5% |
| HD, 15 markers | 136 | 132 | −2.9% |
| FI, 10 markers | 120 | 118 | −1.7% |
| FI, 15 markers | 133 | 128 | −3.8% |

**The loss is consistent and small, but it is a loss.** It is not zero, and the near-conservation is an empirical result in this cohort rather than a theorem; the size of the trade depends on the shape of the hazard–projection relationship. All four configurations lose between 1.5% and 3.8%.

**This analysis is post-hoc.** The specification barred a sex-specific threshold from becoming the primary rule, and that bar stands. It is reported as a design option to be pre-specified in a future study, not as a validated screen. Two constraints travel with it: within-sex selection admits women at lower absolute risk than men (43.1 against 67.3 per 1,000 person-years), so the arms are not exchangeable at baseline and randomisation should be stratified by sex with any sex-by-treatment interaction pre-specified; and the total screening burden is unchanged, merely redistributed evenly across the sexes.

**Two conventions for the event counts differ by up to one death**, because a median of sums is not a sum of medians. The table above uses sum-of-medians, matching the module's own published summary and Figure 6e. Both are written to `pooled_threshold_event_ledger.csv` deliberately, so that the figure and the prose cannot drift apart unnoticed.

#### S6.7 Code and outputs

**Code:** `analysis/08_trial_sorting/` · specification signed off before execution in `SPECIFICATION.md`, with five decisions and their rationale recorded

**Outputs** (in `analysis/08_trial_sorting/outputs/`): `trial_sorting_summary.csv`, `trial_sorting_by_partition.csv`, `sensitivity_sex_specific_{summary,threshold}.csv`, `pooled_threshold_sex_filter.csv`, `pooled_threshold_event_`[illegible] `Table_S_trial_arm_composition.csv`, `Figure_5_trial_arm_sorting.{png,pdf,svg}`, `Figure_5_legend.md`

The two `pooled_threshold_*` files are produced by `code/04_pooled_threshold_sex_filter.py` and support §S6.6. Note that `Figure_5_trial_arm_sorting` is the module's own build; the display item published as **Figure 6** is rebuilt in house style by `npj_aging/figures/build_figures_5_6.py`.

> **FIGURE 5 HERE** — the rule shown for one partition; arm size and sex composition; sex composition against the cohort; mortality enrichment and depletion.
>
> **SUPPLEMENTARY TABLE S7 HERE** — trial-arm composition across configurations.

---

### §S7. Lifestyle construct validity

> Manuscript section: *"Established adverse exposures displace participants along the mortality direction"* · **Figure 6, Table S12**

#### S7.1 Hypothesis and aim

The mortality direction was estimated from prospective disease mortality and nothing else. An axis fitted to an outcome can in principle reflect the outcome model rather than physiology, so it has not yet been shown to correspond to anything independently known to damage health.

**Two questions, with predictions recorded before any result existed.** First, do three exposures established to shorten life — smoking, physical inactivity, poorer diet — displace participants along the axis? Second, if mortality risk depends on the *direction* of displacement rather than its magnitude, does the geometry distinguish exposures by which of the two they move?

#### S7.2 Methodological approach and why

**None of the three exposures contributed to biomarker selection, to estimation of the direction, or to the choice of panel size or identity.** All were fixed before these analyses were specified. *An axis that had seen smoking data would be expected to recover smoking, which is precisely why it was not permitted to.* This is what makes the test a construct-validity test rather than a circular one.

Eight outcomes per exposure: magnitude $m$, alignment $a$ and projection $q$ in each representation, plus the cross-fitted projections of the ten- and fifteen-marker panels — all in analysis-cohort standard deviations so that quantities are comparable.

Models adjust for sex, a four-degree-of-freedom spline in age and survey cycle, with heteroscedasticity-robust intervals. A socioeconomic model adds race/ethnicity, education and family income-to-poverty ratio, **fitted on identical rows to its own comparator** so that adjustment and sample change are not confounded.

**Predictions were recorded in advance and their outcomes are reported including the miss.** Six predictions; five confirmed, one wrong.

> **Amendment 1, 7 August 2026.** The age spline was originally specified as `cr(age, df=4)`, whose basis spans the constant function and is therefore collinear with the model intercept — leaving the design matrix rank 16 of 17 and the robust covariance computed from a singular inverse. Point estimates remained correct via pseudo-inverse, but 72 of 232 standard errors were meaningless. The spline is now centred, restoring full rank, and a rank guard raises rather than returning corrupted intervals silently. Point estimates moved by a median of 0.0003 SD; no conclusion changed. Full record in `MANUSCRIPT_DEFECTS.md` Part 4.

#### S7.3 Results

**All three exposures displaced participants along the direction**, at 0.65 SD (95% CI, 0.60 to 0.70), 0.28 SD (0.24 to 0.32) and 0.17 SD (0.15 to 0.19) respectively in HD coordinates. The direction recovered all three despite never having seen them.

**They separated by which quantity they moved.** Current smoking acted almost entirely on direction — 0.72 SD alignment (0.67 to 0.77) against only 0.15 SD magnitude (0.10 to 0.20) — and after socioeconomic adjustment the magnitude association was abolished (−0.003 SD; −0.06 to 0.05) while alignment held at 0.62 SD (0.57 to 0.67). Mutual adjustment drove magnitude negative (−0.10 SD; −0.17 to −0.03) with alignment still 0.57 SD. Physical inactivity showed the opposite ordering, 0.36 SD magnitude against 0.24 SD alignment. Poorer diet moved both about equally and weakly, 0.17 and 0.16 SD. Former smokers sat close to never smokers at 0.09 SD on both, with no separation between the quantities.

**Not an artefact of prevalent disease:** among 1,950 participants reporting no chronic condition the smoking alignment association was *larger*, at 0.88 SD (0.76 to 0.99).

**Both encodings and both reduced panels reproduced the pattern**, the panels at full strength despite having been selected without any lifestyle information.

**The prediction that missed, reported as a miss.** Physical inactivity had been predicted to move both quantities comparably and instead moved magnitude appreciably more. The geometry separated the three exposures more sharply than anticipated — one acting on direction, one on magnitude, one on both.

#### S7.4 Conclusion

An axis derived solely from mortality recovers exposures it never saw, which is evidence that it indexes physiological deterioration rather than a property of the outcome model. More informatively, it does not treat those exposures as equivalent.

**What this does not establish.** These are cross-sectional associations. They are not intervention effects, evidence of longitudinal responsiveness, or demonstrations that changing an exposure moves a person along the axis. Exposures are self-reported; for smoking, misclassification would attenuate rather than create the observed contrast. Socioeconomic adjustment reduces but does not remove confounding by unmeasured circumstance, and the abolition of smoking's magnitude association under it indicates shared socioeconomic patterning rather than a clean decomposition. Dietary data exist only from 2005 onward. Establishing responsiveness is a precondition for treating the axis as a trial endpoint rather than only as a screening instrument.

#### S7.5 Code and outputs

**Code:** `analysis/09_lifestyle_construct_validity/` · `SPECIFICATION.md` (six predictions recorded in advance; Amendment 1) · `code/lifestyle_associations.py`, `code/build_figure.py`

**Outputs:** `lifestyle_associations.csv`, `Figure_6_lifestyle_construct_validity.{png,pdf,svg}`, `Figure_6_legend.md`; superseded version-1 outputs preserved at `outputs/_v1_superseded_20260807/`

> **FIGURE 6 HERE** — which quantity each exposure moves; agreement between encodings; reduced panels; smoking across model specifications.
>
> **SUPPLEMENTARY TABLE S12 HERE** — all exposure associations, all models.

### §S8. Functional measurement and system coverage

Manuscript section: *"The panel captures inflammation and glycaemia, but not pulmonary function"* · **Figure 7**

#### S8.1 Hypothesis and aim

The panel was assembled from routine blood chemistry, haematology and urinalysis. It was never expected to represent every physiological system, and it contains no direct measure of any organ's mechanical capacity. Two questions follow: which widely used measurements does the panel make **redundant**, and which systems can it **not see**?

#### S8.2 Methodological approach and why

**Candidates were chosen for measurement rationale and statistical power before any result was examined**, and four were excluded with reasons recorded in advance so that the omissions are not silent: cardiorespiratory fitness (measured only to age 49 — zero participants in this cohort), grip and knee-extensor strength (two cycles each, too few events for a test at a comparable standard), and apolipoprotein B (three cycles, and correlated at $r \approx 0.93$ with non-HDL cholesterol, which the panel already contains). **None is claimed uninformative; each is untestable here.**

Each candidate is evaluated in its own eligible cycles, since no common complete-case cohort exists across them, and **every comparison is made on identical rows** — models with and without the candidate are never fitted on different participants.

**The evidence of added information is held-out partial log-likelihood change**, by leave-one-cycle-out within that candidate's cycles, together with the number of folds improved. The conditional hazard ratio is reported alongside.

*Why the unconditional association is not evidence.* A measurement can be strongly associated with mortality alone and add nothing once the panel is present, which is exactly what C-reactive protein does. The unconditional estimate is reported only as a reference quantity.

Candidates are oriented so higher is more adverse ($FEV_1$ reversed). C-reactive protein and insulin are log-transformed, and C-reactive protein is standardised **within assay era**, because the measurement changed between 2009–2010 and 2015–2016 with no data in 2011–2014. Spirometry is restricted to quality grades A and B and adjusted for standing height.

#### S8.3 Results

| Candidate | Alone | Given all 38 | Given ten | Folds improved | Verdict |
|---|---|---|---|---|---|
| hsCRP | 1.31 | **0.98** | **1.01** | 4/7, 2/7 | Redundant; held-out likelihood *falls* |
| Fasting glucose | 1.14 | 1.02 | **1.06** | 5/9, 6/9 | Nothing beyond 38; small gain over ten |
| Fasting insulin | 1.08 | 1.01 | 1.01 | 4/9, 4/9 | Redundant |
| HOMA-IR | 1.12 | 1.01 | 1.03 | 3/9, 5/9 | Redundant |

| Candidate | Alone | Given all 38 | Given ten | Folds improved | Verdict |
|---|---|---|---|---|---|
| **$FEV_1$ (reversed)** | 2.13 | **1.57** | **1.63** | **3/3, 3/3** | **Adds substantially** |

**Inflammation is already captured.** hsCRP was strongly associated alone (1.31; 95% CI, 1.23–1.38) and absent conditional on either comparator, with held-out likelihood falling under both. The most widely used inflammatory marker in ageing research adds nothing to this panel. Its unconditional association is real; it is simply not independent of what the routine markers already measure.

**Fasting glycaemia is captured by the full panel but not fully by the reduced one.** Fasting glucose added nothing beyond all 38 but retained a small association with the ten-marker panel (1.06; 1.02–1.11), held-out likelihood rising 3.4 units across 6 of 9 folds. The reduced panel contains no glycated haemoglobin, which the full panel does, and fasting glucose partially recovers what that omission costs. A stricter eight-hour-fasting definition gave a similar conditional estimate (1.07; 1.00–1.14) but with likelihood no longer improving and only 3 of 6 folds better, so the finding is **suggestive rather than established.** It is nonetheless a specific, reportable limitation of the ten-marker identity, and it falsified a prediction recorded in advance.

**Pulmonary function is not captured at all.** Reduced $FEV_1$ remained strongly associated conditional on both comparators — 1.57 (1.34–1.83) given all 38 and 1.63 (1.39–1.90) given the ten — with held-out likelihood rising 13.2 and 16.2 units and improving in **all three** available folds under both. A direct measure of pulmonary mechanical capacity carries mortality information that 38 routine blood and urine markers do not contain.

**Effect size and replication run in opposite directions, and both facts must travel together.** The $FEV_1$ result is the largest and rests on three folds; the hsCRP null is the best replicated at seven.

**Three of four advance predictions confirmed.** The fourth — that the reduced panel would behave like the full panel for every candidate — was falsified by fasting glucose.

#### S8.4 Conclusion

The panel's coverage is uneven in a way that is informative rather than arbitrary. It already contains the information a widely used inflammatory marker and a fasting glycaemic measure supply; it does not contain, and cannot substitute for, a measurement of how well the lungs work. For a reduced screening panel, adding a blood marker from a system the panel already represents is unlikely to help, whereas a cheap non-invasive functional measurement addresses a genuine gap.

**What this does not establish.** The three candidates were evaluated in different cycles and their estimates are not directly comparable with one another. Examination-based measurements select for the ability to complete a protocol — spirometry excludes participants with contraindications and the fasting subsample is not random — so these estimates apply to participants able and willing to undergo the procedure. The hsCRP analysis spans an assay change with a four-year measurement gap, handled by within-era standardisation but not eliminated by it. **Absence of added information in this cohort does not establish that a measurement is uninformative in general — only that it adds nothing to this panel, in these participants, for this endpoint, over this follow-up.** These are exploratory measurement-boundary tests and did not retrospectively alter the locked panel or the sorting rule.

#### S8.5 Code and outputs

**Code:** `analysis/10_system_coverage/` · `SPECIFICATION.md` (scope decision and four predictions recorded in advance) · `code/import_candidates.py` (raw candidates imported into the module, not into `data_preparation`, because they are inputs to one diagnostic question rather than to the panel)

**Outputs:** `system_coverage_results.csv`, `candidate_{crp,glucose,spirometry}_raw.csv`, `Figure_7_system_coverage.{png,pdf,svg}`, `Figure_7_legend.md`

**FIGURE 7 HERE** — conditional hazard ratios under three specifications, and change in held-out partial log-likelihood with folds improved.

---

### §S8b. External replication in NHANES III (1988–1994)

Manuscript section: *"The finding replicated in an independent cohort under predictions registered in advance"* · **Figure 5**, **Table 3**

#### S8b.1 Hypothesis and aim

Everything to this point is estimated within one survey programme. The obvious alternative explanation is that the geometry describes NHANES 1999–2018 rather than human physiology. NHANES III is a **disjoint** sample of the same population, measured on **different assay platforms** up to three decades earlier, with mortality linkage through 2019. If the finding is a property of physiology it should reappear; if it is a property of the survey it should not.

**Eight predictions, each with a falsification criterion, were recorded and signed off before any mortality-linked analysis was run** — see `NHANES_III/SPECIFICATION.md`. They are reproduced verbatim in Table 3 of the manuscript. This section documents how they were tested and what happened.

#### S8b.2 Methodological approach and why

**Everything was rebuilt inside the replication cohort** — the young-healthy reference, transformation selection, standardisation, and the fitted mortality direction. Only two things transfer: the locked marker *identity* (needed to test transport at all, prediction P7) and the adverse-direction specification, which is a clinical prior rather than an estimate. Transformation *orientation* was recomputed, because it encodes whether the selected transformation reverses the raw-value axis and is therefore a property of the data, not of the prior. Agreement consequently reflects physiology rather than a transported model.

**Thirty-four of the 38 markers were available.** The five-part automated differential is replaced by the three-part Coulter differential, so absolute neutrophil count exists only as total granulocytes; gamma-glutamyl transferase and globulin were fielded in fewer than half of Phase 1 participants. All four were excluded rather than imputed. **Panel transport is therefore evaluated on eight of the locked ten markers**, and that is stated wherever P7 is reported rather than left for a reader to infer.

NHANES III has two survey phases rather than ten cycles, so cross-fitting is grouped ten-fold by **survey primary sampling unit**, preserving the principle applied throughout that correlated sampling units are never split across a fold.

**The 20-year follow-up truncation was pre-specified** in Decision 5 of the specification, before any result existed. This matters: it is the sensitivity that rescues P5, and a reader is entitled to know it was not selected after seeing the primary estimate.

#### S8b.3 Estimation-population declaration

| Quantity | Population | n | Deaths |
|---|---|---:|---:|
| P1, sex–mortality gap | adults 18–79 with mortality linkage | 16,040 | 5,747 |
| P1, panel-complete subset | complete on 34 markers | 13,185 | 4,557 |
| P2–P7b | panel-complete, aged ≥52 at examination | 4,474 | 3,401 |
| Young-healthy reference | rebuilt within NHANES III | 1,053 | — |

The panel-complete cohort is **13,195** participants; **13,185** enter the age-adjusted Cox model, ten being lost to follow-up. Both numbers are correct and appear in the manuscript in their respective places. The young-healthy reference comprises 467 men and 586 women, a mean of 31.0 participants per marker against 32.6 in the development cohort — comparable, and built by the identical cascade (`reference_cascade.csv`). Among those aged ≥52, 51.2% are women.

**S8b.4 Results**

**Seven of eight predictions were confirmed.**

| | Prediction | Criterion fixed in advance | Observed | Verdict |
|---|---|---|---|---|
| P1 | Lower female disease mortality | HR < 1, interval excluding 1 | 0.714 (0.678–0.753) | Confirmed |
| P2 | Women not less displaced | $\Delta m \geq 0$ | +0.005 (HD), +0.070 (FI) | Confirmed |
| P3 | Women lower in alignment and projection | both < 0, excluding 0 | $a$ −0.526, $q$ −0.453 (HD); −0.594, −0.438 (FI) | Confirmed |
| P4 | Magnitude does not explain the gap | HR moves away from 1, or unchanged | 0.707 → 0.689 | Confirmed |
| P5 | Projection removes >50% of the gap | >50% of the sex difference in log hazard | **46.3%** full follow-up; 52.4% at 20 years | **Partially met** |
| P6 | Most signal within clinical limits | >50% | 59.6% | Confirmed |
| P7 | Locked panel transports | above the 95th percentile of a random eight-marker null | 0.8488 vs null p95 0.7774 | Confirmed |
| P7b | Approaches a natively selected panel | ≥90% of the native ceiling | 95.7% of 0.8868 | Confirmed |

**P5 is a partial replication with a measured mechanism, not an excuse.** The attained-age gradient replicates almost exactly: the hazard ratio per standard deviation of projection falls from 2.243 (52–65) through 1.855 and 1.577 to 1.213 (85+), against 2.427 falling to 1.331 in the development cohort. NHANES III follows participants for up to 26 years, so a large share of person-time accrues above age 75 where the axis is weakest. Truncating follow-up recovers the prediction monotonically — 46.3% (full) → 52.4% (20 y) → 59.3% (15 y) → 66.6% (10 y) — and the median exit age falls from 82.8 to 73.6 across those horizons. **Report the primary as primary**; the sensitivity explains it, it does not replace it.

**A striking secondary result.** Given no knowledge of the locked panel, an elastic-net selection run natively within NHANES III independently recovers red cell distribution width, urine albumin-to-creatinine ratio, mean corpuscular volume, serum albumin and granulocytes — five of the locked ten, and the only available neutrophil proxy — in **10 of 10 folds**, with resting pulse in 5 of 10.

**Two deviations are recorded rather than smoothed over.**

1. **The random null used 200 draws, not the 1,000 specified.** A `timeout` wrapper terminated the run. Two hundred draws are sufficient for the p95 decision, and the bootstrap interval on that percentile (0.7669–0.8241) confirms the locked panel clears its upper bound; but the deviation is a deviation and is logged in `claim5_panel_transport.csv` itself.

2. **One random eight-marker panel in a hundred beat the locked eight.** The panel sits at the 99th percentile of arbitrary draws, not beyond chance. The honest claim is that the locked identity is very good, not that it is unique.

Signal-zone location (P6) is computed on the 32 of 34 markers for which clinical limits are defined; 45.9% of positive mortality-associated contribution arises within the young-healthy interval, 13.7% in the corridor between that interval and the clinical limit, and 40.4% beyond the limit.

#### S8b.5 Conclusion

The central finding is not an artefact of one survey programme. In a disjoint cohort, on different assay platforms, up to three decades earlier, women are again no less displaced from young-health than men, again markedly less aligned with the mortality direction, and again die of disease less often; adjustment for magnitude again moves the female hazard ratio *away* from unity; most mortality-associated signal again arises within conventional clinical limits; and a panel selected in one cohort transports to the other far better than chance.

What does not fully replicate is the *magnitude* of the projection adjustment, and the reason is measured rather than assumed: this cohort is followed into a much older age range, where the axis is weaker. That is a boundary on the finding's reach — the geometry is most informative in the decades where geroscience trials would recruit — and it is reported as such.

#### S8b.6 Code and outputs

**Code:** `NHANES_III/` · `SPECIFICATION.md` (eight predictions and falsification criteria, signed off before execution; Decision 5 pre-specifies the truncation) · `NHANES_III_ORIENTATION.md` (file-format guide for readers who have only used continuous NHANES) · `code/01_build_panel.py`, `code/02_select_transformations.py`, `code/04_build_reference.py`, `code/05_build_coordinates.py`, `code/06_sex_mortality_gap.py`, `code/07_direction_geometry.py`, `code/08_followup_sensitivity.py`, `code/09_panel_transport.py`, `code/10_signal_zones.py`, `code/11_functional_measures.py`

**Outputs** (in `NHANES_III/outputs/`): `claim1_female_hazard_ratios.csv`, `claim1_cause_specific.csv`, `claim1_table1_equivalent.csv`, `claim23_sex_contrasts.csv`, `claim23_female_hr_cascade.csv`, `claim4_signal_zones.csv`, `claim4_signal_zones_by_marker.csv`, `claim5_panel_transport.csv`, `claim5_random_null.csv`, `sensitivity_followup_truncation.csv`, `sensitivity_attained_age_gradient.csv`, `reference_cascade.csv`, `reference_parameters.csv`, `transformation_selected.csv`

> **FIGURE 5 HERE** — sex contrasts in both cohorts; the adjustment cascade; signal location; panel transport against the random null; the pre-specified truncation; the attained-age gradient.
>
> **TABLE 3 HERE** — the eight predictions, their criteria and their verdicts.

Raw NHANES III survey files are not redistributed with the analysis compendium; they are permanently hosted by NCHS and their use is governed by the NCHS data use agreement. Download locations are given in `NHANES_III_ORIENTATION.md`.

---

### §S9. Mathematical methods

*Complete specification. A condensed version appears in the main manuscript Methods. Every formula below is implemented in the file named beside it; where the implementation and this text disagree, that is a defect to be reported.*

**Citations marked † must be verified against `manuscript_library.enl` before submission.** They are recorded here from working knowledge and page numbers in particular should not be trusted without checking.

#### S9.1 Notation

| Symbol | Meaning |
|---|---|
| $x_{ij}$ | raw value, participant $i$, biomarker $j$ |
| $T_j(\cdot)$ | locked transformation for marker $j$ |
| $m_j$ | orientation, −1 for reciprocal transformations, +1 otherwise |
| $s$ | sex, male or female |
| $\mathbf{z}_i$ | 38-vector of standardised coordinates for participant $i$ |
| $\beta$ | biomarker log-hazard coefficient vector |
| $\mathbf{u} = \beta/\|\beta\|$ | unit mortality-associated direction |

#### S9.2 Transformation

One transformation per marker is selected from a fixed candidate set by minimising absolute skewness, **on the complete 29,053-person panel**. Selection is unsupervised: age, sex, reference status and mortality are not selection variables.

*Why the full cohort:* the transformation must normalise the distribution it is actually applied to. Selecting on a subgroup normalises that subgroup and can leave the analysis distribution skewed. This was defect 1 (§0.4).

Library: `numpy`, `scipy.stats` · Code: `data_preparation/src/06_transform/`

#### S9.3 Reference standardisation

Two different standardisations are used, deliberately.

**Moment-based (HD branch).** For marker $j$ and sex $s$:

$$z_{ij} = \frac{T_j(x_{ij}) - \mu_{js}}{\sigma_{js}}$$

with $\mu_{js}, \sigma_{js}$ the mean and sample SD in the **same-sex young-healthy reference**.

**Robust (FI branch).** For the same quantity:

$$z_{ij}^{(r)} = m_j \, \frac{T_j(x_{ij}) - M_{js}}{S_{js}}, \qquad S_{js} = \frac{\mathrm{IQR}_{js}}{1.349}$$

with $M_{js}$ the same-sex reference median and $S_{js}$ the normal-consistent robust scale. The divisor 1.349 is $\Phi^{-1}(0.75) - \Phi^{-1}(0.25)$, making $S$ equal to $\sigma$ under normality while resisting tail influence.†

*Why two:* HD requires a covariance matrix, for which moment estimators are the natural and internally consistent choice. The FI is intended to be portable and computable from a lookup table, so it uses robust location and scale that are stable in small clinical reference samples. Multiplication by $m_j$ restores the raw-value axis so that a positive coordinate always means “above the same-sex healthy median” regardless of transformation.

Library: `numpy`, `pandas` · Code: `data_preparation/src/09_standardise/`, `src/11_fi/estimate_fi_reference_parameters`

#### S9.4 Homeostatic dysregulation (HD)

Classical Mahalanobis distance from the same-sex young-healthy centroid:

$$\mathrm{HD}_i = \sqrt{\mathbf{z}_i^{\top} \, \mathbf{C}_s^{-1} \, \mathbf{z}_i}$$

$\mathbf{C}_s$ is the **ordinary unbiased sample covariance (`ddof=1`) of the same-sex young-healthy reference**, inverted directly when full rank.

*Why Mahalanobis:* it is direction-neutral and scale-free, and it accounts for correlation between markers, so a set of individually unremarkable values that are jointly improbable registers as distant. This is the property a per-marker sum cannot reproduce, and it is why HD is retained alongside the FI.

*Sensitivities:* Ledoit–Wolf shrinkage covariance†, and leave-one-marker-out recomputation.

Source: Mahalanobis (1936)†; applied to physiological dysregulation by Cohen et al. (2013)†; shrinkage estimator Ledoit & Wolf (2004)† Library: `numpy.linalg`, `sklearn.covariance.LedoitWolf` Code: `data_preparation/src/12_hd/`

#### S9.5 Sensitised linear frailty index (FI)

Two steps. First the adverse excursion, applying the locked directionality:

$$e_{ij} = \begin{cases} \max(0,\ z_{ij}^{(r)}) & \text{high-only} \\ \max(0,\ -z_{ij}^{(r)}) & \text{low-only} \\ |z_{ij}^{(r)}| & \text{two-sided} \end{cases}$$

The deficit severity **is** that excursion, with no dead zone and no cap:

$$d_{ij} = e_{ij}$$

Then the index, an unweighted mean over the 38 markers, and the signed vector $g_{ij}$ retaining the direction of deviation for the FI direction analysis:

$$\mathrm{FI}_i = \frac{1}{38}\sum_j d_{ij}, \qquad g_{ij} = \mathrm{sign}(z_{ij}^{(r)})\, d_{ij}$$

**The index has no free parameters.** It reads as *"on average, this participant sits* FI *young-healthy scale units into the adverse tail across 38 markers."*

*Why strictly linear:* the instrument is defined as linear, so severity must be proportional to displacement at every magnitude. The superseded specification was flat at **both** ends — a dead zone $a = 1.0$ assigning exactly zero to any deviation within one scale unit, and a cap $c = 3.0$ assigning exactly one to everything beyond three, giving $d = \mathrm{clip}((e - a)/(c - a), 0, 1)$. Both were removed as defect 2 (§0.4). The dead zone zeroed 58.3% of values in the band nearest the reference, where mortality signal is present. The cap flattened 68.7% of waist-to-height values, making the largest single contributor to the index effectively binary for two thirds of the cohort, and contradicted the decision recorded in `TRIMMING_ASSESSMENT.md` not to remove extreme values because they are mortality-enriched.

*Why unboundedness is acceptable:* the $[0, 1]$ bound bought comparability with published frailty indices, which this instrument does not have in any case — its deficits are continuous and reference-relative rather than binary and clinically defined. **Scores are therefore not comparable to published frailty index values**, and a reader assuming a 0–1 proportion will misread them. Averaging across 38 markers prevents any single extreme dominating: the index has median 1.07, interquartile range 0.90–1.26 and maximum 2.67.

*Why transformation is required upstream:* the unweighted mean assumes one scale unit means comparable displacement across markers. Computed on raw values instead of transformed ones, right-skewed markers have small interquartile ranges relative to their tails, and the urine albumin-creatinine ratio alone would carry 18.4% of total burden with a largest excursion of 4,276 scale units against 30 on the transformed scale. Transformation is what makes the mean meaningful; it is not a stylistic step.

*Why unweighted:* an unweighted mean is the classical deficit-accumulation form and requires no fitted parameters, which is what makes it portable. Contributions are nonetheless unequal, since the mean implicitly weights each marker by population drift — waist-to-height carries 10.2% of burden and explains 23.9% of index variance. Mortality-weighted variants are used only where explicitly stated and are a different instrument.

*Deployment note:* the transformation need not be shipped to the point of care. A raw-unit lookup table giving, per marker and sex, the raw values at $z = 0, 1, 2, 3$ allows a clinician to locate a patient by interpolation without computing any transform.

*Prespecified sensitivity:* the superseded bounded grading is retained in code as a parameterised option so the effect of bounding is measured rather than argued.

Source: deficit accumulation, Mitnitski et al. (2001)†; standard construction procedure, Searle et al. (2008)†
Library: `numpy`, `pandas` · Code: `data_preparation/src/11_fi/`

#### S9.6 Survival models

Cox proportional hazards throughout:

$$h_i(t) = h_{0k}(t)\ \exp(\beta^\top \mathbf{x}_i)$$

with **Efron's approximation** for tied event times, **delayed entry** (left truncation) at examination, and **survey-cycle-stratified baseline hazards** $h_{0k}$.

*Why attained age as the primary time scale:* age is the dominant determinant of mortality hazard, so using it as the time scale compares participants at the same age rather than the same follow-up duration, and removes the need to model the age–hazard relationship parametrically. Delayed entry is then required, because participants enter the risk set only at their examination age.

*Why cycle stratification:* assay methods, calibration and population composition differ across survey cycles. Stratifying allows each cycle its own baseline hazard without assuming a common one.

*Why Efron:* more accurate than Breslow with the substantial tie counts produced by month-resolution follow-up, at negligible cost.

*Sensitivity:* time since examination with a four-degree-of-freedom natural cubic B-spline for baseline age. Proportional hazards assessed by scaled Schoenfeld residuals.†

Source: Cox (1972)†; Efron (1977)†; time-scale choice, Korn et al. (1997)†; diagnostics, Grambsch & Therneau (1994)†; Therneau & Grambsch (2000)† Library: `statsmodels.duration.hazard_regression.PHReg` (primary), `lifelines.CoxPHFitter` (regularised fits), `patsy.dmatrix` (splines)

#### S9.7 Regularised marker selection

Elastic-net penalised Cox partial likelihood:

$$\hat{\beta} = \arg\max_{\beta}\ \ell(\beta) - \lambda\left[\alpha\|\beta\|_1 + \tfrac{1-\alpha}{2}\|\beta\|_2^2\right]$$

Sex is included unpenalised. Markers are ranked by absolute penalised coefficient, with marker name as the deterministic tie-break.

*Why elastic net rather than lasso:* biomarkers are correlated, and pure $L_1$ selects arbitrarily among correlated predictors. The $L_2$ component stabilises that choice while $L_1$ still produces sparsity. The mixing fraction $\alpha$ is tuned rather than assumed.

*Why coefficient-magnitude ranking is legitimate here:* the HD coordinates share a common scale — standard deviations of the same-sex young-healthy reference — so coefficients are comparable across markers. This does not hold for arbitrarily scaled predictors.

Source: Zou & Hastie (2005)†; Cox-model coordinate descent, Simon et al. (2011)† Library: `statsmodels PHReg.fit_regularized`

#### S9.8 Geometric decomposition

For a participant with coordinates **z** and unit mortality direction **u**:

$$D = \|\mathbf{z}\|, \qquad L = \mathbf{z}^\top \mathbf{u}, \qquad R = \sqrt{\max(D^2 - L^2,\, 0)}$$

$D$ is total displacement from the young-healthy state, $L$ the component along the mortality-associated direction, and $R$ the orthogonal remainder, by Pythagoras.

*Why decompose:* it separates *how far* a participant has moved from *which way*. The central claim of the paper is that these are different quantities with different consequences, and the decomposition is what makes them separately measurable. $R$ **requires the full 38-dimensional space**: computed within a panel selected for alignment with **u**, there is almost no orthogonal remainder by construction.

Library: `numpy.linalg`

#### S9.9 Validation architecture

**Cycle-grouped splitting.** Training and test sets are whole survey cycles, never individuals.

*Why:* splitting by individual permits a model to exploit cycle-specific assay and calibration artefacts, which inflates apparent generalisation. Grouped splitting forces transfer across a change of measurement era, which is the condition any deployable instrument must meet.†

**Nested validation.** Marker selection, hyperparameter tuning, coefficient estimation and score scaling occur inside the training cycles only; the fitted object is applied unchanged to the held-out cycles.

*Why nested rather than single-split:* tuning on the same data used to report performance is optimistically biased. The inner loop pays that cost internally.

**Percentiles across splits.** Reported as 2.5th–97.5th cycle-split percentiles.

*Why these are not confidence intervals:* the 126 five-versus-five partitions of ten cycles overlap heavily, and each partition's two directions are complementary rather than independent. The spread describes sensitivity to cycle allocation only.

Library: `itertools.combinations`, custom · Code: `analysis/*/code/common.py`

#### S9.10 Uncertainty and multiplicity

- **Bootstrap:** participant resampling stratified by sex and survey cycle, with full refitting of the direction in each replicate.† Library: `numpy.random`
- **Binomial proportions:** Wilson score interval, preferred over the normal approximation at the small event counts encountered in the healthspan screen.† Library: `statsmodels.stats.proportion`
- **Multiplicity:** Benjamini–Hochberg false discovery rate across cause-specific endpoints.† Library: `statsmodels.stats.multitest`
- **Rank and linear association:** Spearman $\rho$ and Pearson $r$. Library: `scipy.stats`
- **Correlation clustering:** stable edge at $|\rho| \geq 0.50$ in $\geq 80\%$ of training halves; clusters are connected components of the resulting graph. Library: `scipy.sparse.csgraph.connected_components`

#### S9.11 Master estimation-population table

*The consolidated audit. Every estimated quantity in the analysis, and the population it is estimated on.*

| Quantity | Estimated on | Why | Used for |
|---|---|---|---|
| Transformation choice | All 29,053 panel members | Must normalise the analysis distribution | HD and FI coordinates |
| HD reference mean, SD | Young-healthy reference, **by sex** | Defines each sex's healthy set point | HD $z$ |
| HD covariance $\mathbf{C}_s$ | Young-healthy reference, **by sex** | Distance is measured relative to healthy correlation structure | HD |
| FI reference median, IQR | Young-healthy reference, **by sex** | Robust set point for a portable index | FI deficits |
| Deficit severity | Not estimated — no free parameters | Severity *is* the adverse excursion; the dead zone and cap were removed (§0.4 defect 2, §S9.5) | FI |
| Cox coefficients (descriptive) | Full analysis cohort | Population the estimate describes | §S1 |
| Cox coefficients (predictive) | **Training cycles only** | Held-out evaluation requires it | §S3–§S6 |
| Score scaling (mean, SD) | **Training cycles only** | Test-set scaling would leak outcome-adjacent information | §S3–§S6 |
| Cutoffs and thresholds | **Training cycles only** | Applied unchanged to held-out cycles | §S5–§S6 |

#### S9.12 Software

Python 3.10. Analyses are deterministic; seeds are fixed and recorded in each module's manifest.

| Package | Version | Used for |
|---|---|---|
| `numpy` | 1.26.4 | linear algebra, geometry |
| `pandas` | 2.2.3 | data handling |
| `scipy` | 1.15.3 | distributions, correlation, graph components |
| `statsmodels` | 0.14.5 | Cox models, regularised Cox, proportions, FDR |
| `lifelines` | 0.30.0 | regularised Cox fits |
| `patsy` | 1.0.2 | spline and design matrices |
| `scikit-learn` | 1.7.2 | Ledoit–Wolf shrinkage covariance |
| `matplotlib` | — | figures |

BLAS threading is capped (`OPENBLAS_NUM_THREADS=1`) during parallel execution to prevent oversubscription; this affects run time only, not results.

**Reproducibility.** Each module verifies SHA-256 hashes of its locked inputs before execution and refuses to run on changed inputs. Analysis fingerprints combine contract, configuration, input and code hashes so that results generated under different settings cannot be silently combined.

---

### Appendix A. Section template

Every section must contain, in this order:

1. **Hypothesis and aim** — what is being tested, or if descriptive, what contrast is being established and why it is needed.
2. **Methodological approach and why** — the mathematics, the data split and training scheme, and the **reason each was chosen over the alternative**. A method stated without its rationale does not satisfy this document.
3. **Estimation-population declaration** — one row per estimated parameter: what, estimated on which population, why that population, what it is used for.
4. **Results** — with the numbers, and with limitations stated inline rather than deferred.
5. **Conclusion** — what this establishes, and explicitly what it does not.
6. **Code and outputs** — links to scripts, source-data files, tests; figure and table insertion points.

---

### Appendix B. Display-item registry

*One authoritative list of every figure and table, what it shows, which section documents it, and which file generates it. This registry is canonical: where the manuscript and this table disagree, the manuscript is wrong.*

#### Main display items

*Numbering follows the manuscript as submitted (V2). Figures are built in house style by the scripts in `npj_aging/figures/`; where a module also produces its own version of a display, the module build is a working figure and the npj build is the published one.*

| Item | Content | Section | Built by |
|---|---|---|---|
| Table 1 | Cohort characteristics by sex | §S1 | `npj_aging/tables/build_tables.py` |
| Table 2 | The locked ten-marker panel | §S4 | `npj_aging/tables/build_tables.py` |
| Table 3 | NHANES III prediction register, P1–P7b | §S8b | `npj_aging/tables/build_tables.py` |
| Figure 1 | Sex–mortality gap; neither magnitude measure explains it | §S1, §S2 | `build_figures_1_3_S1.py` |
| Figure 2 | Displacement equal, direction less aligned | §S3 | `build_figure2.py` |
| Figure 3 | Compression to ten routine measurements | §S4 | `build_figures_1_3_S1.py` |
| Figure 4 | Lifestyle exposures recovered by the axis | §S7 | `build_figure4_lifestyle.py` |
| Figure 5 | External replication against registered predictions | §S8b | `build_figures_5_6.py` |
| Figure 6 | Trial-arm sorting, and the within-sex remedy | §S6 | `build_figures_5_6.py` |
| Figure 7 | System coverage, development cohort | §S8 | `build_figure7.py` |
| Figure 8 | Functional measurement, replication cohort | §S8, §S8b | `build_figures_1_3_S1.py` |

**Figure 6 has five panels.** Panel **d** shows why a pooled threshold produces a male arm (§S6.6); panel **e** is the pooled-versus-within-sex ledger. Earlier drafts had four panels and cited the ledger as 6d.

**Figure 8 was Supplementary Figure S1** until it was promoted into the main text. Output files built before 11 August 2026 carry the stem `FigureS1_functional`.

**Supplementary tables**

| # | Content | Section | Source file |
|---|---|---|---|
| S1 | Disease-condition mortality associations | §S1 | `03/outputs/tables/supplementary_tab` |
| S2 | Marker selection frequency across 252 partitions | §S4 | `06/outputs/data/selection_stability` |
| S3 | Panel subsystem span | §S4 | `06/outputs/tables/Table_S_subsyster` |
| S4 | HD→FI cross-transfer | §S4 | `06/outputs/tables/Table_S_cross_tra` |
| S5 | Panel sensitivities (age, time scale, ridge, LOCO) | §S4 | `06/outputs/tables/Supplementary_Tab` |
| S6 | FI informative-value coverage by marker count | §S4 | `06/outputs/data/representation_comp` |
| S7 | Trial-arm composition | §S6 | `08/outputs/Table_S_trial_arm_compos` |
| S8 | Young-healthy reference intervals and sex-equivalence test, 38 markers | §S0.1 | `10_validation/reference_sex_equival` |
| S9 | Distribution shape and reference intervals by marker and sex | §S0.2 | `10_validation/fi_distribution_sanit` |
| S10 | Magnitude–mortality models with sensitivities | §S2 | `04/outputs/tables/Supplementary_Tab` |
| S11 | HD/FI marker comparison at sizes 10 and 15 | §S4 | `06/outputs/tables/Table_S_marker_co` |
| S12 | Lifestyle exposure associations, all models | §S7 | `09/outputs/lifestyle_associations.c` |
| S13 | System-coverage candidate results | §S8 | `10/outputs/system_coverage_results.` |
| S14 | Cross-fitted signal-zone location by marker | §S1b | `04/outputs/data/signal_zone_marker_` |
| S15 | Adverse-direction assignment for all 38 markers, with physiological basis | §S5 | `data_preparation/config/fi_scoring_` |

**Supplementary figures**

| # | Content | Section | Source file |
|---|---|---|---|
| S1 | Mortality robustness forest | §S1 | `03/outputs/figures/Supplementary_Fi` |
| S2 | All 38 markers by sex, normalised scale | §S0.2 | `Supplementary_Figure_FI_distributio` |
| S3 | Raw distributions with reference overlay, men | §S0.2 | `Supplementary_Figure_raw_distributi` |

| # | Content | Section | Source file |
|---|---|---|---|
| S4 | Raw distributions with reference overlay, women | §S0.2 | `Supplementary_Figure_raw_distribut`[illegible] |

**Manuscript–supplement agreement**

The citation mismatches listed in earlier versions of this registry were applied to the manuscript on 11 August 2026 and are no longer outstanding. Two conventions remain worth stating, because they are the ones most likely to be mistaken for errors:

| Apparent inconsistency | Why it is correct |
|---|---|
| 13,195 versus 13,185 | 13,195 is the NHANES III panel-complete cohort; 13,185 is the subset entering the age-adjusted Cox model, ten having been lost to follow-up. Both appear, each in its own place |
| Eight markers, not ten, in P7 | Only eight of the locked ten exist in NHANES III; neutrophil count is available solely as total granulocytes, and gamma-glutamyl transferase and globulin were fielded in fewer than half of Phase 1 participants |

The directionality diagnostic (module 07) carries no display item in this manuscript. It is a companion-paper analysis and is deliberately absent here; see that module's README for why it must not be used to respecify the frailty index.

---

### Appendix C. Building this document

`SUPPLEMENTARY.md` is the editable source. The PDF is generated, never edited.

```
cd "Manuscript_One\github_final\manuscript and supplementary"
powershell -ExecutionPolicy Bypass -File .\build_supplementary.ps1
```

Add `-Docx` to emit a Word version for collaborator comments; equations become native Word OMML, which is adequate for commenting but typesets less well than the PDF.

**Toolchain.** pandoc 3.9.0.2 and TinyTeX (TeX Live 2026) at `%APPDATA%\TinyTeX`. Cambria and Cambria Math are specified deliberately: the pandoc default Latin Modern lacks the Greek and relational glyphs used in running text and drops them **silently**. The build script counts missing-character warnings and reports them; a clean build reports zero.

**If TinyTeX must be reinstalled**, note that `tinytex::install_tinytex()` fails in this environment — `yihui.org/tinytex/TinyTeX-1.zip` returns 404 and R's download methods are unavailable. Fetch the release asset directly from `github.com/rstudio/tinytex-releases/releases/latest` and extract it to `%APPDATA%`.
